# Whole-body single-cell atlas of an adult vertebrate in homeostasis and regeneration

**DOI:** 10.64898/2026.02.03.703562

**Authors:** Kutay Deniz Atabay, Patrick Aoude, Chanyoung Park, Mykola Kadobianskyi, Olivier H. Paugois, Benjamin Judkewitz, Peter W. Reddien

## Abstract

A complete cell-type transcriptome atlas of a vertebrate could promote understanding of animal cell-type composition, organization, and evolution. The miniaturized, transparent, and regenerative teleost *Danionella cerebrum* brings whole-vertebrate single-cell profiling experiments within reach. We performed region-stratified single-cell RNA sequencing across the adult *Danionella* body and mapped cell types and gene expression at single-cell resolution using spatial transcriptomics. We delineated spatially-distinct neural cell types based on their regional gene expression signatures. The body-wide atlas elucidated cell types harboring adult positional information, uncovered paedomorphic features, and revealed conserved body-region and appendage-specification programs in adult connective tissue. Comparative analyses revealed conserved neural cell types, and regeneration datasets uncovered expression dynamics during telencephalon regeneration. This whole-vertebrate transcriptome atlas yields a comprehensive resource for myriad questions in biology and neuroscience.

## Introduction

Early investigation into biology involved the description of major cell and tissue types (*1–6*). Modern molecular advances have made possible similar investigation of organism components, but now at the level of spatially mapped gene expression for cell types and organs (*7–14*). The molecular mapping of the cell transcriptomes comprising complete animal anatomy has not been achieved for an adult vertebrate. Vertebrates exhibit numerous specialized cell types and enhanced cell-type complexity through innovations such as neural crest-derived cell types, the adaptive immune system, sophisticated skeletal and connective tissue cell types and structures, specialized organs, and intricate nervous systems that share architectural similarities across major regions (*15–18*). Most vertebrates are also large –the complexity and size of current vertebrate models are barriers to complete molecular access to every adult cell type. The size of most vertebrates might also limit the scope of future whole-animal molecular studies following perturbations and drug treatments, across diverse physiological conditions, and across the lifespan. These challenges could, in principle, be overcome through investigations of miniaturized vertebrates.

*Danionella cerebrum* is a miniaturized teleost fish found in streams near the Bago Yoma mountain in Myanmar (*19*), is one of the world’s smallest vertebrates (∼12 mm) (*19*), and has the smallest vertebrate brain on record, containing ∼650,000 neurons within 0.6 mm^3^ (*20*) (∼4X the *Drosophila* brain, (*21*)), yet also contains all major vertebrate brain regions (*20, 22, 23*). *Danionella cerebrum* is genetically tractable and optically transparent (Fig. 1A) and displays robust behaviors, including associative learning, schooling, and audible vocalizations (*24–28*). Thanks to its optical transparency, *Danionella cerebrum* is arguably unparalleled for non-invasive live whole-brain and whole-body imaging at single-cell resolution in an adult vertebrate (*29*); work on *Danionella* has already enabled investigation of diverse neural dynamics topics (*26, 30–35*). *Danionella cerebrum* is also closely related to zebrafish, which allows for the transfer and use of an expansive molecular toolkit, and its genome, similar to many other teleosts, is ∼half the size of zebrafish (*36, 37*).

**Figure 1.**
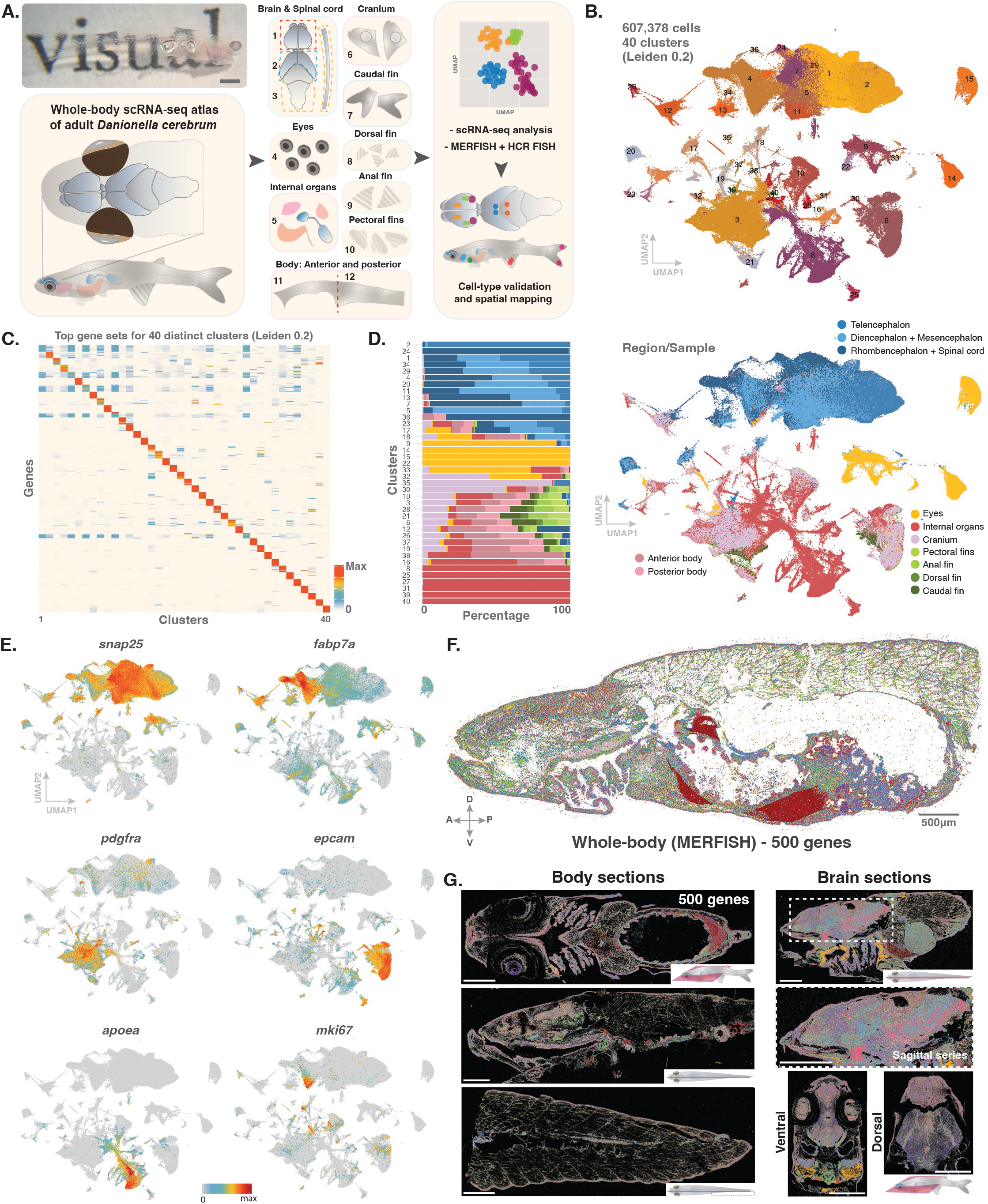
A whole-body single-cell transcriptome and spatial atlas of adult *Danionella cerebrum*. **(A)** Adult *Danionella cerebrum* (unpigmented, *tyr*^-/-^), (scale bar, 1mm), illustrations of the regions of the WT adult body used for single-cell RNA sequencing, and experimental approach summarized. **(B)** UMAP representation of all cells colored by general cell classes/clusters. **(C)** Heatmap showing the expression patterns of differentially expressed marker genes across 40 general cell classes (table S1). **(D)** Barplot showing distribution of experimental samples/body parts across different clusters (left); UMAP representation showing distribution of cells from experimental samples (body parts) (right). **(E)** UMAP feature plots of gene expression for cell-class markers for neurons (*snap25a*), neural stem cells/radial glia (fabp7a), fibroblasts (*pdgfra*), simple epithelial cells (*epcam*), intestinal enterocytes and hepato-intestinal cells (*apoea*), and dividing cells (*mki67*). **(F)** MERFISH of an anterior body section showing expression of a 500-gene panel, including specific cell-type and tissue markers. **(G)** MERFISH with representative body and brain sections (scale bars, 500 µm) showing expression of the 500-gene panel.

We utilized adult *Danionella cerebrum* to generate a complete vertebrate single-cell transcriptome atlas at saturation. We analyzed the regional distribution of cell types and performed trajectory and comparative analyses of neural stem cells, progenitors, and mature neuron classes. We found that *Danionella cerebrum* is broadly regenerative and performed scRNA-seq in the telencephalon after injury to define molecular and cellular features of neural regeneration. We also used whole-animal multiplexed high-throughput fluorescent in situ hybridization (MERFISH) and Hybridization Chain Reaction (HCR) FISH to spatially resolve cell types and gene expression domains, producing a spatial map of gene expression and cell types. Together, these analyses yielded a comprehensive spatial transcriptome atlas of an adult vertebrate.

### Single-cell RNA sequencing and profiling cell types of the whole organism

We dissected *Danionella cerebrum* into 12 samples: telencephalon, diencephalon and mesencephalon, rhombencephalon and spinal cord, eyes, internal organs, cranium, caudal, dorsal, anal, and pectoral fins; and anterior and posterior trunk (Fig. 1A). We used scRNA-seq (10X Genomics) to profile cells from each sample separately; 607,370 cells passed stringent quality control steps (fig. S1A-D). The resulting atlas was comprised of 40 broad tissue classes (Fig. 1B, C, table S1). The dissection of separate samples enabled mapping body cells across clusters (Fig. 1D-F, fig. S1E, table S1). We assembled a large-scale (500-gene) MERFISH panel to spatially resolve the expression patterns of select top marker genes from the atlas (*9, 10, 12, 38–46*) (table S2). Adult *Danionella* size allowed MERFISH to be applied in sections across whole-body axes, including a diverse set of neural cell types (Fig. 1G).

### Global classification of cells following binning into four main groups of tissues

We next sought to annotate an atlas for ∼all *Danionella* cell types. Our approach was to cluster cells from binned micro-dissected anatomical categories to identify similar tissue-class cells (e.g., neural) from different regions. We then merged these similar cell classes for subclustering to identify individual cell types (e.g., different classes of neurons). We binned cells from the 12 samples into five anatomically relevant categories: *(i)* central nervous system (excluding eyes): telencephalon; diencephalon and mesencephalon; rhombencephalon and spinal cord samples; *(ii)* eyes; *(iii)* internal organs; and *(iv)* core body tissues (i.e., skin and musculoskeletal system), which included cells from cranium and body, *(v)* all fins (Fig. 2A). We clustered cells within these five bins and annotated resultant clusters (fig. S2-6, table S3). From the clusters for each bin, we identified four major tissue classes: *(i)* neural, *(ii)* epithelial: organ-lining, secretory, parenchymal, and reproductive cells, *(iii)* connective, and *(iv)* muscle (Fig. 2B). We merged cells for each of these four tissue classes from the 5 anatomical bins and subclustered the data. For example, all neural cell types across the body were combined into a single Seurat object to facilitate the discovery of new cell types and core modules within this tissue class. In-depth annotations revealed a diversity of cell types for each functional cell class across the entire body (Fig. 2, table S3). MERFISH enabled visualization of the spatial distribution of selected markers across the body and within the central nervous system (Fig. 2).

**Figure 2.**
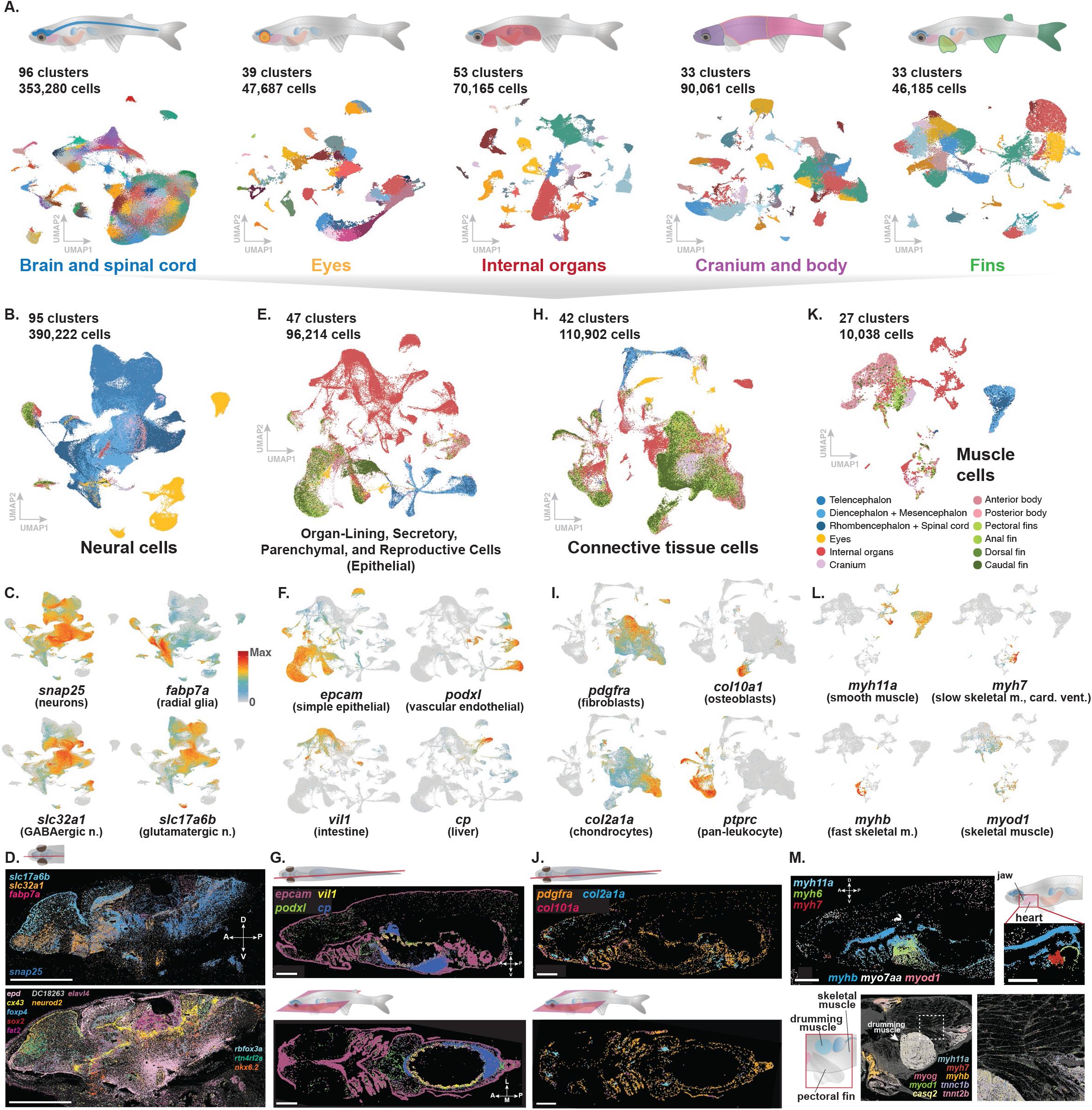
Cell type diversity revealed by regional/organ-level or tissue-type classification. **(A)** Cells subclustered based on their experimental sample identities, organized in different organs or body parts (cluster markers and annotations, tables S1, S3). **(B)** UMAP plot showing all neural cells of the whole body, including neurons, glia, neural stem cells, oligodendrocytes, and Schwann cells; colors indicate sample identities. **(C)** UMAP feature plots showing gene expression for mature neurons (*snap25a*), radial glia (*fabp7a*), GABAergic neurons (*slc32a1*), and glutamatergic neurons (*slc17a6a*). **(D)** MERFISH panel showing expression of genes in panel C (top), and a select set of genes showing regionally distributed expression patterns for leptomeningeal cells (*epd*), glia/radial glia (*cx43*), neural stem cells/radial glia (*sox2*), cerebellar granular layer (*fat2*), immature neurons (*neurod2*), subset of mature neurons (*elavl4*), *Danionella cerebrum* transcript ID: DC18263 (predicted best BLAST hit, *PLEKHG1*) marking a single central neuronal layer in the optic tectum, interneurons, optic tectum/ventral pallium (*foxp4*), mature neurons (*rbofox3a*), dorsal pallium, glutamatergic neurons (*rnt4rl2a*), and *nkx6.2*+ cells, across dien-, mesen-, and rhombencephalon. Scale bars, 500 µm **(E)** UMAP plot showing epithelial, organ-lining, parenchymal, and reproductive cells of the whole body. **(F)** Example UMAP feature plots showing gene expression for simple epithelium (*epcam*), vascular endothelial cells (*podxl*), intestinal cells (*vil1*), and liver cells (*cp*). **(G)** MERFISH showing expression of genes from panel F across the anterior sagittal and dorsoventral body sections. **(H)** UMAP plot showing all connective tissue cell types of the body. **(I)** Example UMAP feature plots showing gene expression for fibroblasts (*pdgfra*), osteoblasts (*col10a1*), chondrocytes (*col2a1a*), and leukocytes (*ptprc*). **(J)** MERFISH panels showing expression in fibroblasts, chondrocytes, and osteoblasts. **(K)** UMAP plot showing all muscle cell types of the body. **(L)** UMAP feature plots showing gene expression for smooth muscle (*myh11a*), slow skeletal and cardiac ventricular muscle (*myh7*), fast skeletal muscle (*myhb*), and general skeletal muscle. **(M)** MERFISH panels showing distinct muscle cell types across the body, including the jaw, heart, trunk, and drumming muscle (pool of genes regionally expressed in the drumming muscle); Panels G, J, M, scale bars: 1 mm.

### Neural tissues

The neural dataset harbored 390,222 cells in 95 molecularly distinct cell clusters, representing central and peripheral nervous systems (Fig. 2B-D; fig. S7). Consistent with transcriptome atlases from other species including human, marmoset, mouse, and other teleosts (*12, 13, 47, 48*), clustering, indicated that neuronal and glial cell lineages were mainly clustered based on their developmental origin or regional bias, rather than by neurotransmitter-defined identities (explored below). MERFISH confirmed the presence of *fabp7a*+ radial glia distributed throughout the brain, forming neurogenic niches (Fig. 2D) that coarsely resembled juvenile to 1-6-month-old zebrafish (*49–51*) with a potentially more continuous periventricular distribution of neural stem cells along the axes of the nervous system. Broadly, in the forebrain, *slc17a6b*+ glutamatergic neurons had a dorsal bias, whereas *slc32a1*+ GABAergic neurons showed a ventral bias. Furthermore, *emx3*, *foxg1a*, *sp8a*, and *tbr1b*+ radial glia and their lineages mainly localized in the forebrain and were clustered together in the UMAP space (*52*). Specifically, *eomesa*, *egr3*, *pbx3a*, and *stxbp1b*+ cells clustered together and were localized in the dorsal telencephalon (pallium), and separately, *dlx1/2/5/6*, *gsx1/2*, *lhx6*, *nkx2.1*, and *isl1*+ cells were localized in the ventral telencephalon (subpallium) (fig. S8). *eomesa* expression in the pallium, and *dlx1/2/5/6*, *lhx6*, *nkx2.1*, and *isl1* expression in the subpallium were previously observed in adult zebrafish (*42, 52–54*); however, pallial localization of *egr3*, *pbx3a*, and *stxbp1,* and subpallial expression of *gsx1/2* along with its colocalization with these other subpallial markers, have not been described in an adult teleost. We delineated a group of genes, including transcription factors such as *zeb2*, *her5*, *lhx9*, *hoxc4a*, *znf536, neurod1/2* linked with patterning, neurogenesis, and specification; along with receptors, ion channels, ligands, and neuropeptides that showed regional expression patterns in cerebellum, optic tectum, and hindbrain, marking functional and anatomical boundaries (fig. S9). Additionally, similar to previous observations in adult zebrafish (*42, 46*), *eomesa* and *emx2* expression were largely segregated to distinct cells in pallium and subpallium, respectively (fig. S10). Genes broadly marking glutamatergic and GABAergic function showed a coarse expression pattern across the CNS (fig S11). Cells from the dorsal and ventral telencephalon clustered away from cells of other brain regions and were segregated based on their dorsal-versus-ventral gene expression signatures across telencephalon clusters (fig. S8A, B). Cells from diencephalon and mesencephalon samples were clustered together and expressed key patterning and fate-specification factors, including *sp8a*, *en2a*, and *tcf7l2* (fig. 8C, D). Similarly, cells from the rhombencephalon clustered together in the UMAP space and expressed key transcription factors such as *hoxa4a* and *irx5a*, indicating persistent transcriptional signatures aligned with early patterning programs that underlie distinct brain regions (fig. S8C, D).

### Non-neural tissues

The epithelial dataset contained 92,214 cells in 47 distinct clusters across organs and body parts and involved organ-lining, secretory, parenchymal, and reproductive cell types (Fig. 2E-G, fig. S12, table S1). *epcam*+ cells were widely distributed throughout the body, including for different organ systems; *vil*+ and *cp*+ expression was restricted to cells of the intestine and liver, respectively; and *podxl*+ endothelial cells were distributed throughout the body, yet enriched in the brain and heart, with distribution along the trunk (Fig. 2G).

The connective tissue dataset contained 110,902 cells, 42 clusters, representing different fibroblast types, chondrocytes, osteoblasts/cytes, osteoclasts, and mesenchymal progenitors, as well as hematopoietic-system cells, including macrophages, mast cells, leukocytes, erythrocytes, and thrombocytes (Fig. 2H-J). Major populations contributing to the connective tissue dataset were *pdgfra*+ fibroblasts, *col2a1a*+ chondrocytes (primarily), *col10a1a*+ osteoblasts, and *ptprc*+ leukocytes (Fig. 2I, fig. S13). We observed prominent distribution of *pdgfra*+ fibroblasts throughout the organism (Fig. 2J).

The muscle dataset contained 10,038 cells and 27 clusters of distinct muscle cell types (Fig. 2K-M), including visceral and vascular smooth muscle cells (*myh11aa+*); cranial/jaw muscle (*myhb*); cardiac muscle (ventricular, *myh7*+, atrial, *myh6+*; myocardial, *myl7+*); differentiated skeletal muscle and its progenitors (*myod1*+) (Fig. 2P); differentiating myocytes (*myog)*; and slow (*tnnc1b+)* fibers. We also identified satellite cells and myoblasts across regions (fig. S14). Separately, visceral and vascular smooth muscle cells (mural cells) formed separate clusters for the internal organs and brain, indicating regionalized tissue-specific specialization for these cell types (fig. S14, table S1).

We performed a separate analysis for the drumming muscle that regulates male sonic output (fig. S15A). Using MERFISH, we observed prominent expression of *tnnc1b* and *actc1*, genes typically expressed in slow muscle, as well as expression of *casq2*, a gene associated with rapid calcium release and uptake, indicating rapid contractile capability. *fabp11a* and *plvapa* expression was consistent with specialized vascularization, possibly aiding metabolic demand. We also observed broad *sim1a* expression and posterior *hoxc8a* expression. Beneath the thin muscle fibers of the drumming muscle, at the core, we observed *acanb* and *lum* expression, genes primarily expressed in connective tissue, indicating a fibrocartilaginous element consistent with previous observations (*19*), and *scg3* expression, a gene primarily expressed in neuroendocrine cells (fig. S15A). *tnnt2b* was selectively expressed at the surface and near the core, facing *acanb*+ cells. Drumming muscle in *Danionella* and other teleost species with sonic muscles has been reported to be rich in mitochondrial genes (*27, 55, 56*). We isolated muscle clusters with high mitochondrial gene expression, and excluded myocardiocytes, which identified a candidate gene expression signature for drumming muscle (fig. S15B-D, table S1). To assess whether this atlas reached ∼complete cell-type saturation, covering even the rarest states, we examined *pth*, *rln3a*, *avp*, and *galn*+ neuropeptidergic neurons, which have overall numbers ranging from 30 to 150 cells across the entire CNS (fig. S16A, B). We also searched for other known rare cells/cell types, and cells with complex morphologies (e.g., ciliated) across different tissues, such as neuroepithelial cells of the gills (NECs), Kolmer-Agduhr neurons, and sensory hair cells, which were all detected in the atlas (fig. S16C). This indicated that the *Danionella* atlas includes even the sparsest cell types/states in the body that survived the experimental protocols, thereby reaching ∼saturation for nearly all major transcriptomic states in an adult vertebrate.

### Constitutive adult positional information and *Danionella cerebrum*

Teleosts exhibit remarkable regenerative capacity (*51, 57, 58*). Similarly, *Danionella* was able to rapidly regenerate caudal fins (fig. S17). Adult positional information exists in some regenerative species and can enable regeneration. For example, dozens of regionally expressed genes in planarian muscle regulate the regeneration of regional cell types (*59, 60*). Some factors are constitutively expressed to define positional memory in adult axolotl (*61–63*), and Hox genes are regionally expressed in adult human skin (*64*). Molecular access to all cell types across all adult body axes and organs enabled elucidation of cell types that display constitutive regional gene expression, including potential components of positional information. Comparative cell-type analyses showed that connective tissue, including fibroblasts, retained regional expression of genes that function in embryonic patterning (fig. S18). To explore whether canonical positional information genes, such as *Hox* genes that regulate the formation of anteroposterior body axis regions and proximal-distal (PD) limb pattern during development, are expressed regionally in the adult *Danionella cerebrum*, we isolated all fibroblasts (31,756 cells, 19 clusters), based on *pdgfra* expression, from anterior and posterior trunk samples (Fig. 3A). This included all fibroblasts from non-neural tissues and *pdgfra*+ glial progenitors and perivascular fibroblasts of the CNS (fig. S19A-C). DGE analyses showed that *Hox* genes were among the most differentially expressed in the anterior and posterior trunk regions (Fig. 3B-D). A separate analysis for multiple connective tissue lineages, including fibroblasts, chondrocytes, osteoblasts, as well as neural, muscle, and epithelial cells, showed that other connective tissue cells, neurons, radial glia, and muscle progenitors also retained regional expression patterns of *Hox* genes. By contrast, *epcam*+ epithelial cells and *myod1*+ muscle cells did not (fig. S18A). Regionally delineated *Hox* gene expression was strongest in connective tissue cells compared to neural and muscle cell types (Fig. 3E, fig. S18B, C, table S4). An expanded list of genes differentially expressed across regions (table S5) yields targets for investigating the maintenance of adult regional tissue identity.

**Figure 3.**
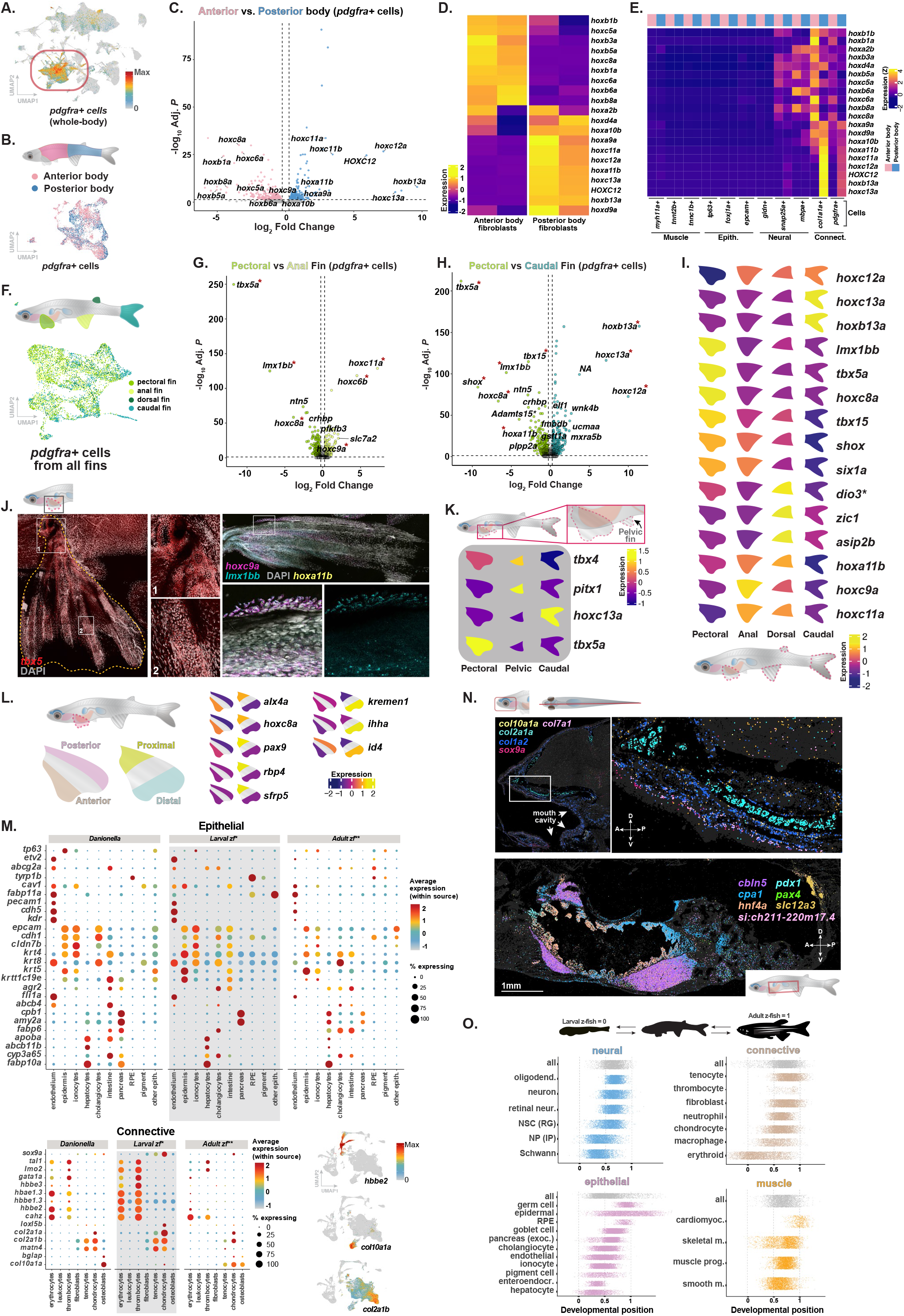
Positional memory and paedomorphic gene expression signatures in adult *Danionella cerebrum*. **(A)** UMAP feature plot showing all *pdgfra*+ cells (fibroblasts) across the body. **(B)** UMAP feature plot showing a subset of fibroblasts isolated from anterior and posterior trunk samples. **(C)** Volcano plot showing the top differentially expressed genes between anterior and posterior trunk fibroblasts, Hox genes are selectively labeled. **(D)** Heatmap showing an expanded list of Hox genes, known to be regionally expressed across the anteroposterior axis of the body, are differentially expressed in anterior and posterior trunk samples of the adult *Danionella cerebrum*. **(E)** Heatmap showing Hox gene expression in cells positive for key tissue markers across the anteroposterior body axis. **(F)** Illustration and UMAP representation of different fins and fin fibroblasts, respectively. **(G-H)** Volcano plots demonstrating the top differentially expressed genes between pectoral and anal fin fibroblasts, and between pectoral and caudal fin fibroblasts. Expression of conserved limb/appendage patterning genes *tbx5a*, *lmx1b*, *tbx15*, and *shox* is enriched in pectoral fin compared to anal or caudal fins. **(I)** Summary heatmap showing the top differentially expressed genes across fins, revealing fin-specific gene expression programs in the adult state. **(J)** HCR-FISH showing expression of *tbx5a*, *hoxc9a*, and *lmx1b* in the adult pectoral fin. **(K)** Summary heatmap showing the top differentially expressed genes across pelvic, pectoral and caudal fin fibroblasts, revealing pelvic fin-specific expression of *pitx1* and *tbx4*. **(L)** Summary heatmap showing anteriorly enriched expression patterns for *alx4a*, *hoxc8a* along with other proximally and distally enriched gene expression patterns across pectoral fin regions. **(M)** Comparison of adult *Danionella cerebrum* and larval and adult zebrafish gene expression patterns for a set of curated genes. *Larval zebrafish atlas: ZMAP (*80*); **adult zebrafish atlas: pooled dataset from 11 sources (*45, 46, 81–89*). Expression patterns indicate widespread expression of keratins across cell types; primarily embryonic/larval-like blood (larval hemoglobins: *hbbe2*, *hbae1.3*) and cartilogenic program (*col2a1b*, *matn4*) along with expression of *col2a1a* (mature chondrocytes) and *col10a1a* (osteoblasts); UMAP plots show *hbbe2*, *col2a1b* and *col10a1a*. **(N)** Top MERFISH shows layered distribution of *sox9a*+ chondrocyte progenitors, *col7a1*+ dermal fibroblasts, *col1a2*+ interstitial fibroblasts and tenocytes, *col2a1a*+ chondrocytes, and *col10a1a*+ osteoblasts in the mouth cavity and upper jaw. The right panel shows a zoom-in of the white rectangular area. Bottom MERFISH panel shows markers expressed in internal organs. Scale bars: 1 mm. **(O)** Comparative analysis between adult *Danionella cerebrum,* larval zebrafish (data source*: ZMAP (*80*)), and adult zebrafish (data source**: pooled dataset from 11 sources (*45, 46, 81–89*), grouped by tissue class, with a pooled “all” cells (top rows; grey cells represent atlas cells in that tissue class summarizing that specific tissue class). Each dot is a single atlas cell along the larval (0) to adult (1) axis, colored by tissue class, indicating the “developmental” position of diverse *Danionella* cell types of different tissue classes (0: closer to larval zebrafish; 1: closer to adult zebrafish). Photoreceptor and amacrine cells are merged into a “retinal neuron” group, and all fibroblast subtypes into a single “fibroblasts” group, and re-scored against their merged reference. Cell types are ordered by ascending median. 3000 atlas cells per cell type are shown.

Because *Danionella* fins regenerate, we reasoned that some genes might be constitutively expressed associated with fin-type-appropriate regeneration. *tbx5a*, *lmx1b*, and *hoxc8a* were preferentially expressed in the pectoral fin (Fig. 3F-I; table S5). These results are notable for multiple reasons: *Tbx5* is the major factor promoting forelimb identity development in tetrapods, and the pectoral fin is likely orthologous to forelimbs (*65, 66*). *Lmx1b* specifies dorsal identity in limbs, and *hoxc8a* has a role in limb regionalization (*65, 67, 68*). Conversely, the anal fin preferentially expressed *hoxc6b*, *hoxc9a*, and *hoxc11a*. A comparison between pectoral and caudal fins revealed that the pectoral fin preferentially expressed *shox*, *tbx15a*, and *hoxa11b*. *shox* encodes a proximal mesenchyme transcription factor that primarily shapes stylopod/zeugopod growth and chondrogenesis (*69*). Similarly, both *tbx15a* and *hoxa11b* have been implicated in forelimb development in tetrapods (*70, 71*). By contrast, *hoxc12a*, *hoxb13a*, and *hoxc13a* were expressed more highly in the caudal fin than the pectoral fin, perhaps consistent with its posterior identity (*72, 73*), (Fig. 3I, fig. S19D, table S5). A separate analysis comparing dorsal and anal fins revealed *zic1* as a preferentially expressed gene in the dorsal fin (fig. S19D, table S5). *zic1* was previously shown to be expressed by the dorsal part of the caudal fin in developing medaka fish (*74*), and was implicated in dorsal patterning and neural crest development (*75*). Next, we isolated pelvic fins for scRNA-seq. A comparison of pectoral and pelvic fin fibroblasts revealed *tbx4* and *pitx1* as top genes in pelvic fins (Fig. 3K, fig. S19E, table S5). *Tbx4* and *Pitx1* are arguably the two most prominent tetrapod hindlimb-identity-specifying factors (*66, 76*), supporting the possibility that these factors represent a form of positional memory in adult pelvic fins.

We identified additional genes with enriched expression in fibroblasts from distinct fins, which can be investigated for roles in adult fin identity in regeneration in the future (table S5). Furthermore, we physically separated fin regions for additional scRNA-seq; revealing anterior and proximal *alx4a* and *hoxc8a* and posterior and distal *id4* enrichment for pectoral fins, among other regionally expressed genes (Fig 3L, fig. S20A-E). These findings extend the conserved gene regulatory networks that direct vertebrate appendage/limb formation into the adult teleost context, and reveal the persistent fin-specific deployment of ancestral appendage/limb tissue-patterning programs.

### Molecular analysis of paedomorphic features suggest a mosaic state in *Danionella cerebrum*

*Danionella* species display heterochrony in the form of prominent progenetic paedomorphosis, characterized by accelerated sexual maturation and substantial loss of adult skeletal structures, including dorsal skull bones, adult retention of larval-like features, loss of pigmentation, and absence of scales (*19, 20, 77–79*). To assess potential paedomorphic signatures in the adult *Danionella cerebrum*, we surveyed curated marker panels for connective, epithelial, muscle, and neural states alongside thyroid hormone signaling pathway components (fig. S21, 22), and performed three-way Spearman correlations between the *Danionella* atlas, a reference larval (ZMAP) atlas (*80*), and adult zebrafish data (constructed by pooling 11 single-cell transcriptome datasets (*45, 46, 81–89*)) (Fig 3M-O; fig. S23, 24, table S6). Globally, adult *Danionella* data positioned between larval and adult zebrafish, with both adult-like (i.e., cardiomyocytes, germ cells) and larval-like (i.e., erythroid cells, Schwann cells) clusters present for each tissue class. Consistent with this, mosaic signatures emerged in blood (embryonic hemoglobins), abundant cartilage (*matn4*, *col2a1b*), and internal organ, connective, and neural tissue cell types (Fig. 3M-O). TH receptors were broadly expressed but varied across neural and non-neural tissues, which could be investigated for connection to heterochrony. Details are in Supplementary text (Molecular analysis of paedomorphic features), in figs. S21-24 and table S6.

### Regional identity drives neuronal transcriptomic similarities

95 molecularly distinct cell clusters were distributed across the central and peripheral nervous systems (Fig. 4A, B, fig. S7). We grouped cells from this dataset into glutamatergic neurons, GABAergic neurons, neurons with other neurotransmitter (NT) identities (cholinergic, glycinergic, dopaminergic, serotonergic, noradrenergic, and histaminergic neurons), radial glia (*fabp7a*, *sox19a*, *atp1a1b*, *id1*, and *sox2*+) and intermediate progenitors (*ascl1a*+), oligodendroglia and Schwann cells (Fig. 4A, B, fig. S7B). We then subclustered glutamatergic neurons (57 clusters), GABAergic neurons (36 clusters), and other NT-class neurons (28 clusters) (Fig. 4A-F, fig. S25-27, table S1). In all cases, clusters appeared regionally biased, suggesting regional origin as a determinant of transcriptional similarity among neurons. Glutamatergic, GABAergic, and other NT neurons from the eye and the peripheral nervous system (PNS) formed distinct clusters (Fig. 4C-E). Analysis of transcription factors with expression in different neuronal classes revealed specific transcription-factor signatures for brain regions, as well as a peripheral signature (Fig. 4G, fig. S28) (*38, 40, 42*). MERFISH revealed anatomical boundaries and regionalization for general glutamatergic, GABAergic, and other neurotransmitter marker genes, whereas cell-type-determining transcription factors displayed regional expression patterns (Fig. 4H). Differential gene expression analysis between glutamatergic and GABAergic neurons, and radial glia across different regions of the CNS showed genes with regionally enriched expression patterns (Fig. 4I, fig. S29-31, table S7). This intricate spatial and molecular map of an entire adult nervous system defines its global organization and can serve as a resource for myriad future investigations into the development, function, and regeneration of vertebrate nervous systems.

**Figure 4.**
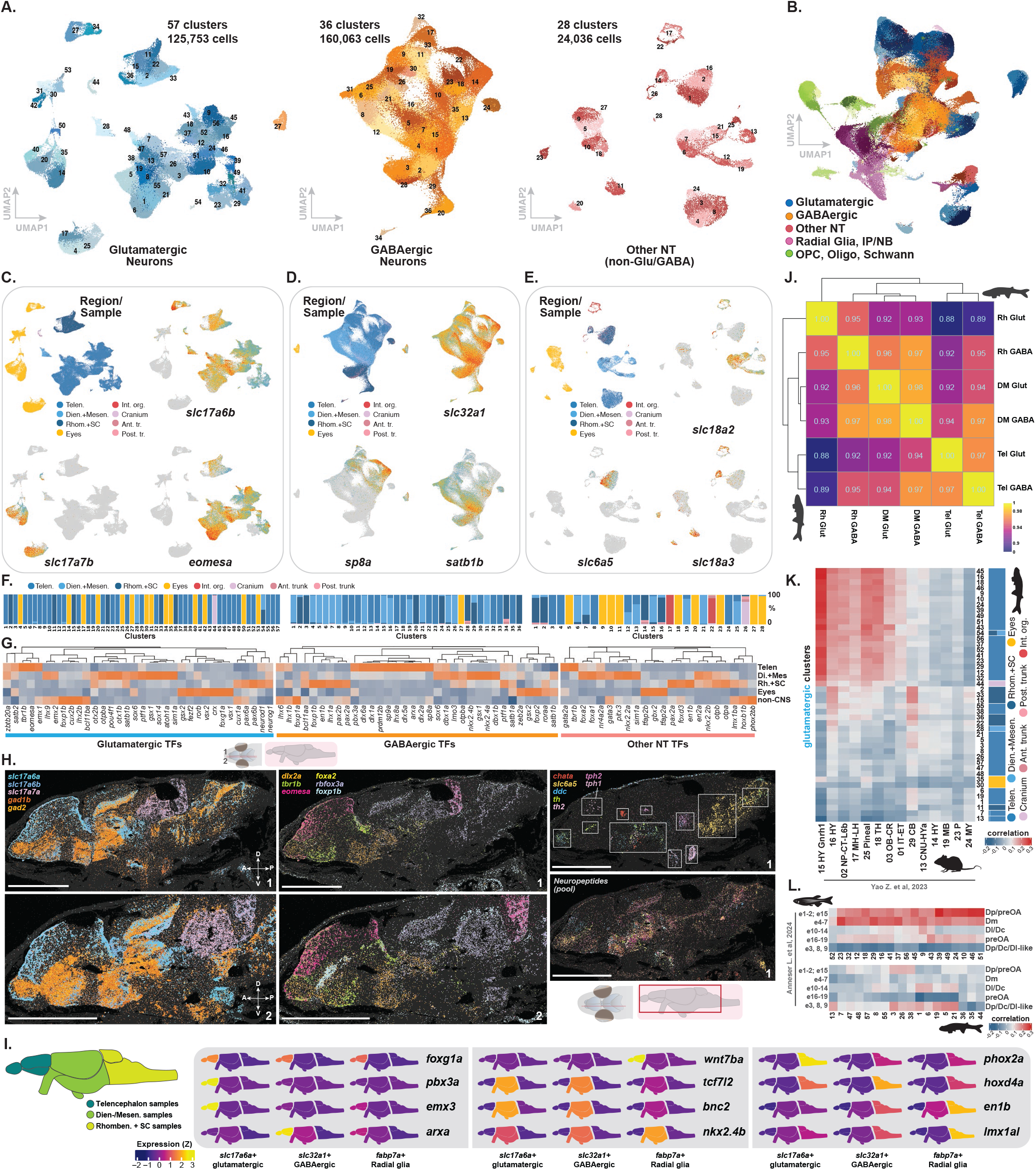
Cellular diversity across neural tissues in the adult central and peripheral nervous system. **(A)** UMAP plots showing diversity of glutamatergic, GABAergic neurons, and neurons with other neurotransmitter identities across the adult body. **(B)** UMAP representation of all neural cells colored based on neurotransmitter identities and cell class (neuronal and glial). **(C)** UMAP plots showing sample distributions across glutamatergic clusters and expression of *slc17a6a* (general glutamatergic neuron marker), *slc17a7a* (a subset of glutamatergic neurons of the eye), and *eomesa* (a forebrain glutamatergic marker). **(D)** UMAP plots showing sample distributions across GABAergic neuron clusters, and expression of *slc32a1* (pan-GABAergic neuron marker), *sp8a* (marking inhibitory neurons in the telencephalon/olfactory bulb), and *satb1b* (marking a subset of interneurons across dien-, mesen-, and rhombencephalon). **(E)** UMAP plots showing sample distributions across neuron clusters with other neurotransmitter identities, and expression of *slc18a2* (common in histaminergic, serotonergic, and dopaminergic neurons with other unique markers), *slc6a5* (glycinergic neurons), and *slc18a3* (cholinergic neurons). **(F)** Barplots showing distribution of experimental samples across glutamatergic, GABAergic, and other-NT clusters. **(G)** Heatmaps showing brain and body-region-specific expression of transcription factors known to define different subtypes of glutamatergic, GABAergic, and other NT neurons. **(H)** MERFISH showing expression for glutamatergic and GABAergic markers for two representative sagittal positions (left panels); regionally segregated expression of a select panel of transcription factors (middle panels); markers indicating distribution of other-NT neurons across the brain (right, top panel), and an expansive marker pool (gene lists is presented in table S7) labeling neuropeptidergic neurons across the brain (right, bottom panel). **(I)** Summary heatmap showing expression patterns for top differentially expressed genes for *slc17a6a*+ glutamatergic neurons, *slc32a1*+ GABAergic neurons, and *fabp7a*+ radial glia across regions (expanded heatmap in fig. S31). **(K)** Heatmap of comparative cross-species analysis (SAMap) between mouse (*13*) and *Danionella cerebrum* glutamatergic neurons. **(L)** Heatmap showing a comparative cross-species analysis (SAMap) between zebrafish (*42*) and *Danionella cerebrum* telencephalic glutamatergic neurons.

There was greater similarity between neurons from different neurotransmitter classes (glutamatergic and GABAergic) from the same brain region, when compared to the same neurotransmitter class from different regions (Fig 4J). To study what drives transcriptional similarities (i.e., region versus neurotransmitter identity), we collapsed each neuronal cell cluster/type into an average profile (metacells) and built a dendrogram to assess profile similarity (*47*). We performed hierarchical clustering of the metacells, generated a heatmap of pairwise correlations using highly variable genes, and annotated each metacell’s region, neurotransmitter identity, and experiment of origin. Major branches organized by region, with neurotransmitter classes only organizing within region-based blocks. Different experimental batches were assessed for impact, and we also assessed whether the clusters matched regional labels (using the adjusted RAND index, a measure of how well two groupings agree at different clustering resolutions). These tests suggested that region was the key determinant driving transcriptomic similarities between neurons in the adult *Danionella* CNS, especially for glutamatergic neurons (fig. S32). Analyses performed using both all genes and highly variable genes between neurons further indicated region/developmental origin as a driver of molecular similarity between neural cells.

We explored cell-type homology between *Danionella* and mouse neurons, radial glia, and neuronal progenitors using self-assembling manifold mapping (SAMap (*90*)) (Fig. 4K; fig. S33-35). *Danionella* telencephalon glutamatergic clusters showed broad similarity to mouse thalamic-habenular/diencephalic (18-TH, 17-MH-LH), pineal (25-Pineal), and neuroendocrine-associated clusters (16-HY and 15-HY-Gnrh1), whereas *Danionella* hindbrain clusters showed similarity to mouse cerebellar granule-like excitatory clusters (29-CB). GABAergic compartment showed similarity to mouse subpallial/basal ganglia and hindbrain/cerebellar GABAergic classes spanning telencephalon, dien-/mesencephalon and rhombencephalon (09-CNU-LGE, 07 CTX-MGE, 08-CNU-MGE, and 28-CB), consistent with broadly deployed medial/lateral ganglionic eminence (MGE/LGE)-like transcriptional programs. Radial glia clusters matched mouse glial classes (30 Astro-Epen and 32-OEC) and were broadly anti-correlated with neuronal classes (fig. S35B), supporting an epedymoglial/radial-glial identity, with coarse correlation to astro-ependymal tanycyte, and olfactory/telencephalic astroglial states, whereas region-specific radial glia matched to astroependymal/tanycte and Bergmann glia (fig. S35). Regional cross-species cluster similarities were more pronounced in glutamatergic neurons (table S8).

We also analyzed cell-type homology with zebrafish (42) and *Danionella* telencephalon using SAMap (Fig. 4L, fig. S36, table S8). *Danionella* glutamatergic neuron clusters mapped best onto the dorsal-pallial region (Dp), the olfactory recipient of the posterior zone, and the dorsal-medial zone (Dm). Another match extended to the preoptic area clusters and broadly to Dp/Dc/Dl-like zones. Overall, this analysis suggested that glutamatergic cells in the olfactory and limbic subdivisions are conserved between the two teleosts. Analysis of GABAergic clusters showed similarity to clusters of MGE, CGE, preoptic *galn*+ neurons, LGE-derived clusters and a ventromedial cluster, suggesting that the subpallial interneuron program is shared between the *Danionella* and zebrafish telencephalon.

### Neural stem cell and glial cell distribution and diversity in the *Danionella cerebrum* CNS

Teleost nervous systems are also broadly regenerative (*51, 58, 91*). We explored the diversity of radial glia and intermediate progenitors, after subclustering (49,423 cells and 33 clusters); this dataset also contained newborn and immature neurons across the organism (Fig. 5A, fig. S37, table S1). These progenitors also displayed regionally biased transcriptomic signatures across clusters (Fig. 5B, C). The most abundant cell type was radial glia, including both cells with dividing and quiescent signatures (Fig. 5D, E). MERFISH with sagittal brain sections revealed a broad distribution as well as regional localization of radial glia and intermediate progenitor markers across pallium, subpallium, ventral diencephalon, optic tectum, mesencephalon, and rhombencephalon (Fig. 5F).

**Figure 5.**
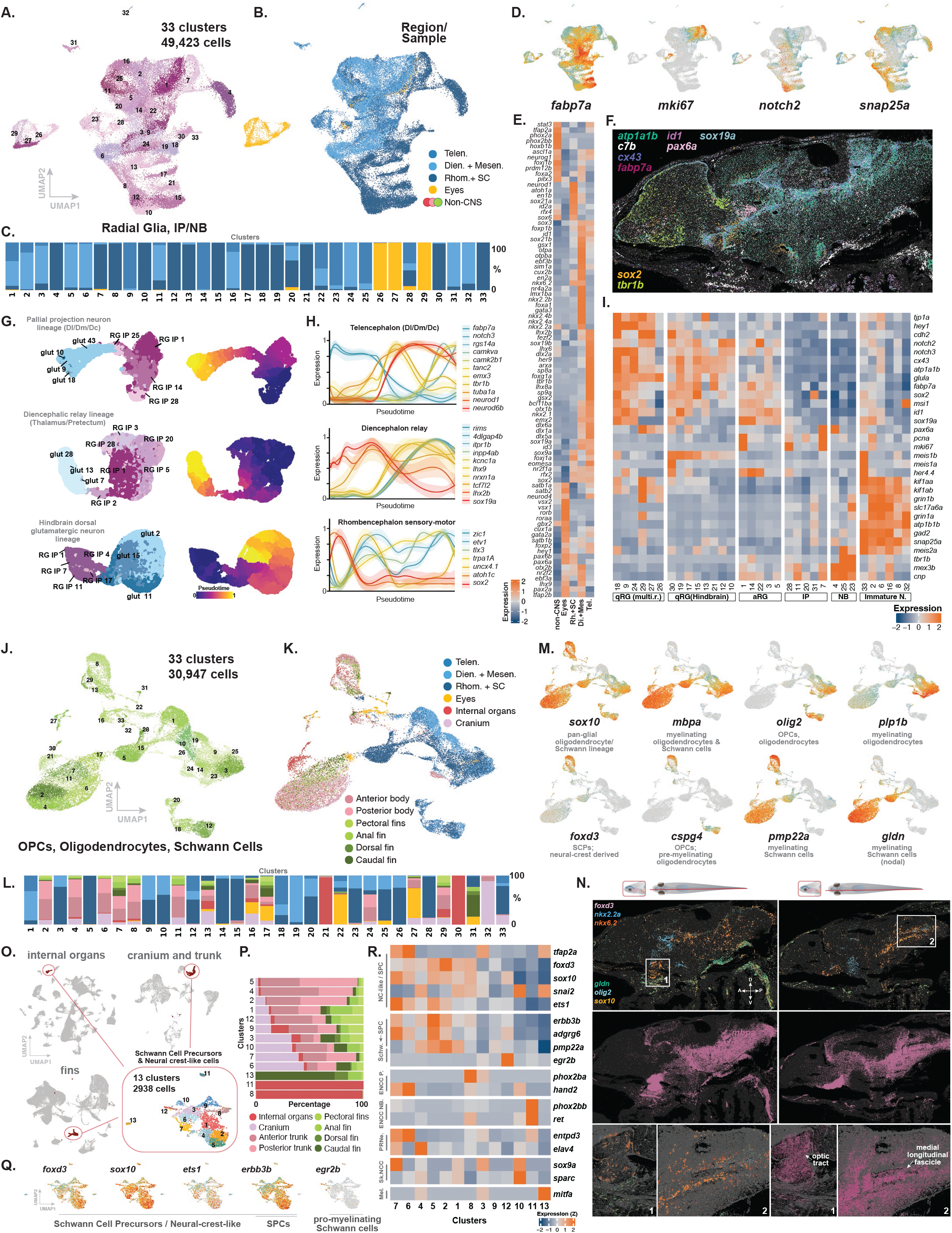
Neural stem cells and glial cell types across the adult body. **(A)** UMAP showing 33 distinct clusters containing different glial cell classes, including radial glia and newborn neurons. **(B)** UMAP representation showing distribution of experimental samples across glial clusters. **(C)** Barplot showing the percentage distribution of experimental samples across different glial clusters. **(D)** Feature plots showing radial glia (*fabp7a*), dividing cells (*mki67*), quiescent radial glia (*notch2a*), and newborn/immature neurons. **(E)** Heatmap showing transcription factors known to be associated with different neural progenitor classes/states, showing regional expression patterns across brain/body regions. **(F)** MERFISH showing expression of radial glia and intermediate progenitor markers in concert. **(G)** Pseudotime analysis showing predicted glutamatergic neuron differentiation continua for three separate brain regions: pallium, diencephalon, and rhombencephalon. Cells are colored based on parent glutamatergic and radial glia/intermediate progenitor colors in figures 4A-B and 5A. **(H)** Pseudotime gene expression analysis for each predicted differentiation continua, marking specific progenitor and newborn/immature neuron states. **(I)** Heatmap showing differential expression of a curated marker set shown to be linked with quiescent and active radial glia states, intermediate progenitors, and newborn/immature neurons. **(J)** UMAP showing 33 distinct clusters comprised of oligodendrocytes, oligodendrocyte progenitors, and Schwann cells. **(K)** UMAP representation showing distribution of experimental samples across oligodendrocyte and Schwann cell clusters. **(L)** Barplot showing the percentage distribution of experimental samples across oligodendrocyte and Schwann cell clusters. **(M)** UMAP feature plots showing gene expression associated with the pan-glial progenitor state and with different stages of oligodendrocyte and Schwann cell maturation. **(N)** MERFISH showing expression of markers of oligodendrocytes, oligodendrocyte progenitors, Schwann cells and their progenitors, and myelinated fiber tracts, including the optic nerve and medial longitudinal fascicle. **(O)** Illustration of the approach showing isolation and subclustering of Schwann cell precursors from distinct body samples, including internal organs, cranium, trunk, and fin samples. **(P)** Barplot showing distribution of experimental samples across clusters in O. **(Q)** UMAP feature plot showing expression of key marker genes for Schwann cell precursors (SPCs) as well as neural-crest-like cells. **(R)** Heatmap showing differential expression of a list of markers previously associated with SPCs and neural crest cells (SPC: Schwann cell precursor; NC-like: neural crest-like; Schw: Schwann cells; ENCC P: enteric neural crest cells - progenitor; ENCC NB: newborn enteric neurons; PRNe: Peripheral neurons; Sk. NCC: Skeletal neural crest cells; Mel: melanocytes)

We investigated neurogenesis in the adult *Danionella cerebrum* CNS using pseudotime-based inference. We prioritized glutamatergic neurons because of their regional production dynamics, as opposed to the complex, tangential, and long-range migratory behaviors of interneuron progenitors (*92, 93*). We analyzed telencephalon; diencephalon and mesencephalon; and rhombencephalon and spinal cord, separately. Using pseudotime inference, we identified cluster pairs with transcriptional similarity between the radial glia/neuroblast and glutamatergic neuron datasets using correlation plots (fig. S38). Selected pairs were used to connect radial glia to neuroblasts to neurons in representative trajectories per region (Fig. 5G, H, fig. S38, 39, table S1). Different modes of teleost neurogenesis have previously been observed (*51, 94–96*). We identified three distinct putative differentiation continua: a pallial projection neuron trajectory (representing dorsolateral, dorsomedial, and dorsal-central pallium identities); a diencephalic relay neuron trajectory with a thalamus/pre-tectum signature; and a hindbrain dorsal glutamatergic neuron trajectory. Differentially expressed genes identified by pseudotime analysis, along with known genes associated with specific neuronal trajectories (*52, 97–103*), exhibited distinct gene expression patterns along pseudotime (Fig. 5H).

We classified radial glia into quiescent radial glia (qRG) and *meis1b*+ posterior (i.e., hindbrain) qRG, expressing known maintenance factors *id1*, *notch2/3,* and *hey1* (*40, 99, 101, 104*). Active radial glia clusters were classified by *fabp7a*, *sox2*, and *sox19a,* and cycling cells by *pcna* and *mki67* expression. Intermediate progenitors/neuroblasts, and newborn/committing neurons expressed *mex3* and *tbr1b, respectively* (*40, 99, 105*). Finally, fate-committed immature neurons were annotated (Fig. 5I) (*40, 99*). Combined with regionally expressed transcription factors, cell-state markers also indicated heterogeneity and regional transcriptional variance in the radial glia and intermediate progenitor populations (fig. S30C) (*95, 99, 106*). We compared *fabp7a*+ quiescent and active radial glia from telencephalon, diencephalon, mesencephalon, and rhombencephalon samples and identified the top differentially expressed genes across these regions. This confirmed key regional distinctions, indicating expression of *foxg1a*, *emx3*, *wnt7ba*, and *eomesa* in the telencephalon; *crabp1a*, *pax7b*, *nkx4.2*, *pomca*, and *en2* in the dien-/mesencephalon samples, and *irx5a* and *nipal2* in the rhombencephalon and spinal cord samples (fig. S30C, table S7).

We also explored diversity within the oligodendrocyte and Schwann cell populations, comprising 30,947 cells and 33 distinct clusters distributed across the central and peripheral nervous systems (Fig. 5J, K, L, fig. S40, table S1). We detected distinct maturation stages, including their progenitors (Fig. 5M, N) (*107, 108*). Myelin signal was largely distributed along the fiber tracts across the dien-/mesencephalon and the rhombencephalon (Fig. 5N). Both OPCs and myelinating oligodendrocytes were enriched around the optic nerve and medial longitudinal fascicle in the hindbrain.

### Neural-crest-like cells in the adult *Danionella cerebrum*

Exploration of cell classes in internal organs, cranium, trunk, and fin samples revealed three distinct clusters of cells with transcriptomes resembling Schwann cell precursors (SCP) as well as a neural-crest-like program. We isolated and subclustered these cells to analyze heterogeneity within this notable cell population, as well as regional and tissue-of-origin differences among them (Fig. 5O, P). We identified this population based on the specific expression of *foxd3*, *sox10*, *erbb3b*, *ets1*, *gpm6ab*, and *ngfrb*, genes previously linked to Schwann cell precursor and neural crest states (*109–112*). A detailed analysis based on tissue origin of these cells indicated two populations: a dominant SCP population and a candidate neural-crest-like cell population, primarily associated with the enteric nervous system as well as skeletal structures (i.e., cranium and fins) (Fig. 5P-R, fig. S41, table S1). The presence of these populations raises the possibility of new incorporation of Schwann cells and possibly distinct neural crest-derived cell lineages into multiple anatomical regions, which can be investigated in the future.

### Regenerative dynamics in the *Danionella cerebrum* telencephalon

*Danionella cerebrum* exhibited nervous system regeneration, including missing parts of the telencephalon (fig. S42A). EdU-labeling showed widespread cellular incorporation of new cells into numerous tissues across the adult body (fig. S42B-E). In teleosts, the telencephalon is everted, with radial glia on the outside. generating neuronal lineages through stacking (*113, 114*). The teleost telencephalon regulates higher-order processing, spatial cognition, olfactory processing, learning, memory, and social dynamics (*115–117*). Comparative studies using single-cell transcriptome datasets have revealed molecular similarities between the cell types and anatomical regions of the teleost forebrain and those of tetrapods, including striatal, hippocampal, and cortical cell types (*114, 115, 118, 119*), rendering this structure an appealing target for exploring regenerative dynamics. We developed microsurgical techniques to remove tissue or induce an extensive injury down the anteroposterior axis within the telencephalon of adult *Danionella cerebrum*. Following a unilateral injury, we collected the injured and contralateral (uninjured) telencephalon 12 hours, 3 days, 7 days, 14 days, and 30 days post-injury (Fig. 6A). Injuries caused an immediate (seconds) and transient (hours) loss of transparency in the injured tissue, which was visibly cleared within 2-4 weeks, suggesting a mechanism maintains transparency (fig. S42F). Following regeneration (day 30), EdU+ cells were positioned along the injury site, restoring the overall tissue architecture and marking cells that incorporated into the telencephalon (Fig. 6B).

**Figure 6.**
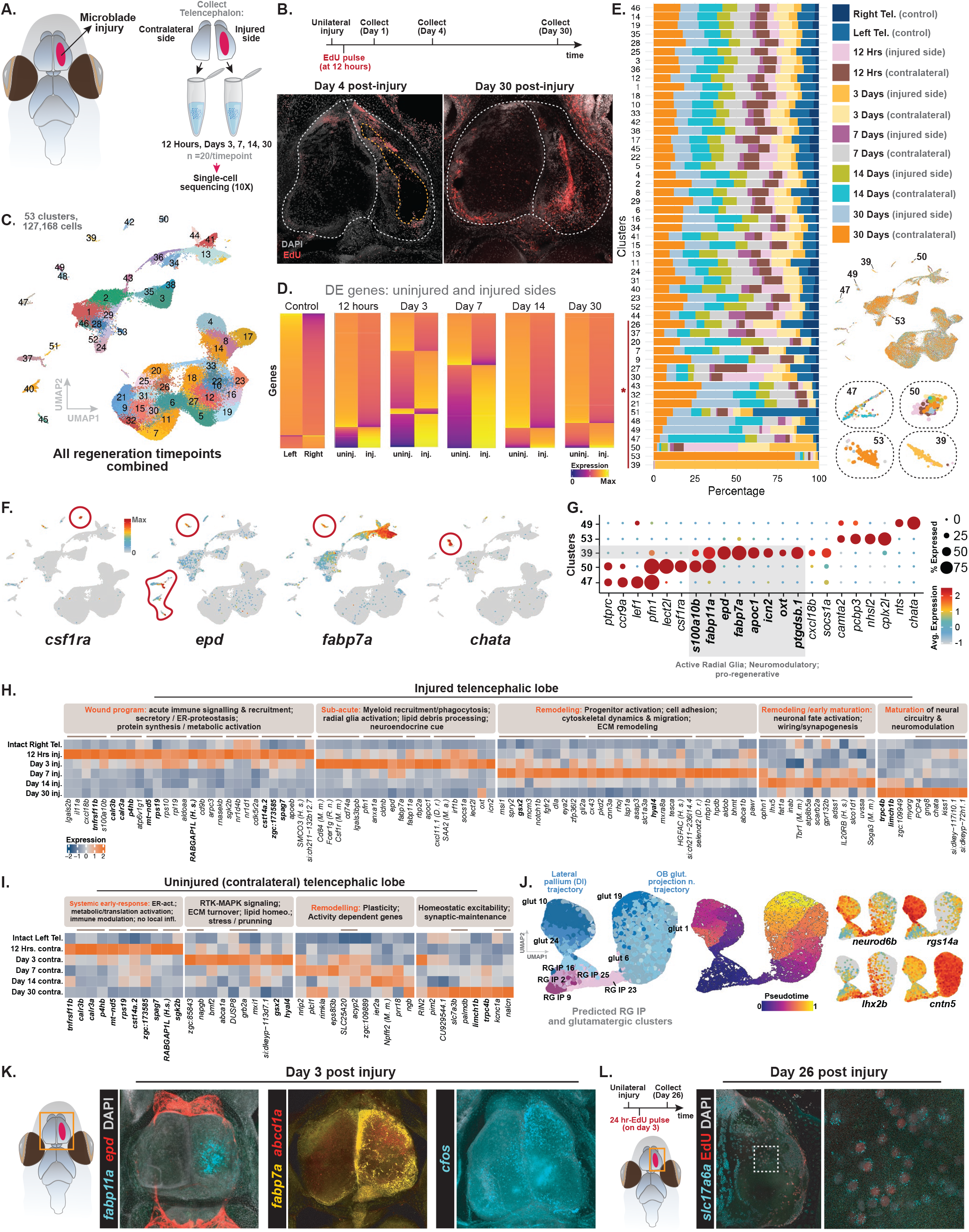
Telencephalon regeneration following injury in the adult *Danionella cerebrum*. **(A)** Illustration showing the experimental procedure used for unilateral telencephalon injury and tissue collection. **(B)** Schematic display of experimental timeline for EdU labeling of the injured telencephalon at day 30; two separate examples are shown. **(C)** UMAP plot of all time points pooled for the injured and contralateral (uninjured) telencephala. **(D)** Heatmaps showing global differential gene expression changes across different time points for the injured and contralateral (uninjured) sides of the telencephalon. Shown values are relative, and the order of the genes across time points is not fixed. **(E)** Barplot showing the percent contribution of each time point to UMAP clusters, indicating the presence of time-point-specific clusters: 49, 47, 50, 53, and 59. Sample-specific-clusters with high Jensen-Shannon divergence scores are marked with asterisks. **(F-G)** UMAP and dot plot showing top genes expressed in each time-point-specific cluster, indicating a wound response that is comprised of immune cell signaling and recruitment, radial glia activation, neuromodulatory signaling, neuronal remodeling, and maturation. **(H-I)** Timepoint-specific analysis indicating top genes expressed in each phase, comprised of wound response, regeneration, and neural maturation for both injured and uninjured (contralateral) telencephalic side. **(J)** Pseudotime analysis predicting regenerating glutamatergic neuron lineages are predominantly clusters 1, 6, 10, 19, and 24. UMAP plots are colored based on parent glutamatergic and radial glia/intermediate progenitor colors (left), and pseudotime (middle). Genes expressed in distinct newborn or immature neurons (*neurod6b*), progenitors (*lhx2b*), and mature neurons with dorsolateral or dorsomedial telencephalic identity (*rgs14a*+), which are predicted to be regenerating, are plotted (right). **(K)** HCR-FISH for day 3 enriched markers, *fabp11a*, *epd* and *fabp7a*; and *cfos*; n=4 **(L)** HCR-FISH performed on day-26 fixed brain samples following injury and 24-hour-long EdU-labeling on day 3; showing mature EdU+-*slc17a6a*+ glutamatergic neurons in the regenerating telencephalic hemisphere.

Analyses of telencephalon regeneration (127,168 cells, 53 clusters; 5 timepoints) revealed injury-specific cell states with acute changes occurring between 12 hours and 7 days following injury, and a differential response between injured and uninjured sides (Fig. 6C-G, fig. S42G, S43, table S9). Time-resolved expression signatures elucidated a stepwise program on the injured side: an acute wound response program and myeloid cell recruitment, a shift to immune modulation with radial glia proliferation and neurogenesis by day 3, structural remodeling and early neural maturation programs (Notch/FGF activation, markers of progenitors such as *fabp7a*) between days 3-14, and circuit/neuromodulation maturation by day 30. The contralateral side showed a parallel but distinct systemic response involving metabolic activation and stress modulation, a possible compensatory gene expression signature without myeloid cell recruitment (Fig. 6H, I). Specific clusters (c39, c50) linked to neurogenesis based on their gene expression signatures expressed pro-regenerative repair factors such as *epd* (ependymin) (*99*). This cluster also expressed *cxcl18b* and *socs1a*, suggesting an immune-reactive state (*120, 121*). Regeneration-related clusters expressed genes implicated in neuronal maturation and circuit re-establishment, such as *cplx2l*, *chata*, and *camta2* (*122–124*). We also observed expression of *nts* (neurotensin) and *oxt* (oxytocin) in injury-specific clusters, genes activated in the axolotl brain following tail injury (*125*). Pseudotime analyses of predicted regenerating glutamatergic lineages predominantly recovered dorsolateral pallium (*rgs14a*+) and olfactory projection neurons (*cntn5*+) (Fig. 6J; fig. S44). HCR-FISH confirmed expression of key upregulated genes (*fabp11a*, *fabp7a*, *abcd4a* and *epd*) potentially linked with a regeneration-associated program involving radial glia activity, fatty acid handling, and extracellular support (Fig. 6K). We also confirmed, through EdU labeling, integration and maturation of *slc17a6a*+ glutamatergic neurons in the injured telencephalon (Fig. 6L). Cluster-level markers and timepoint-specific gene expression signatures are discussed in detail in Supplementary Text (Telencephalon regeneration).

## Discussion

The miniaturized anatomy and transparency of *Danionella cerebrum* present unique opportunities for whole-vertebrate experiments with single-cell resolution, ranging from spatial mapping of cell types to high-throughput connectomics and imaging of neuronal activity across the nervous system. This miniaturization enabled us to generate a whole-body transcriptome atlas of all vertebrate cell types for the first time and will enable future studies across diverse conditions and specific perturbations – potentially including regeneration, pharmacological studies, mutant conditions, aging, and diverse physiological states. Even rare cell-type populations were captured at frequencies exceeding their anatomical abundance in the body in this atlas. This oversampling approach suggested the atlas essentially saturated coverage of cellular diversity in the adult state. *Danionella* is particularly powerful for neuroscience research, enabling both monitoring and manipulation of neural activity patterns in the adult brain. Furthermore, transparency can facilitate systems-level investigations into inter-organ communication, neuroimmune interactions, and gut-brain signaling in a living adult organism.

Studying adult fibroblasts revealed that the ancestral *Hox*-guided positional system remains active and molecularly harbored by connective tissue cell types in adult *Danionella cerebrum.* Our analyses show that conserved appendage/limb-patterning programs are also persistently deployed in an adult fin-specific manner in *Danionella cerebrum*. This included classic limb/appendage-initiating transcription factors, such as *Tbx5* (forelimb/pectoral fin) and *Tbx4* and *Pitx1* (hindlimb/pelvic fin) (*76, 126*). Multiple other genes with developmental patterning roles showed persistent adult regional expression, identifying genes that can be studied in adult tissue maintenance and repair. The retention of regional expression of key developmental factors, primarily in connective tissues, might reflect a persistent adult limb/fin patterning program that could enable tissue homeostasis and capacity for regeneration.

We observed a mixed pattern of paedomorphic signatures across *Danionella* tissues. Heterochronic features in *Danionella*, including agenesis of several bones (e.g., dorsal skull bone), lack of scales, life-long transparency, and a primarily cartilaginous skeletal system, have been previously reported (*19, 78, 127–129*). We observed persistent expression of embryonic hemoglobins in adult *Danionella* (*130*). Epithelial, connective, neural and muscle gene expression patterns indicated a mixed state, with both adult- and larval-like transcriptomic states across the cells and cluster of these tissue classes. The molecular hallmarks and mechanisms of paedomorphic features will be interesting to investigate using this molecular resource. It will be interesting to investigate whether paedomorphic features in adult *Danionella* impact regenerative attributes or cellular turnover, or are associated with adaptive benefits in their environment through larval-like traits such as small size and transparency. The *Danionella* genus comprises five species that exhibit additional morphological variation, providing potential for future comparative evolutionary studies (*19, 78, 79, 127, 128*).

Our analysis of neural cell types revealed regional patterns of gene expression in both neurons and neural progenitors. Adult teleost (zebrafish) radial glia are heterogeneous, with regard to their regional identity and degree of differentiation (*95, 131, 132*). Consistent with these observations, the pattern of transcription factor expression in radial glia, intermediate progenitors, and newborn neurons showed telencephalon-enriched expression modules (*foxg1a*/*emx3*/*eomesa*, along with *dlx2a*/*arxa*/*sp8a* and *wnt7ba*); diencephalon/mesencephalon-enriched expression modules (*pomca*, *en2a*, *nkx2*.*4b*, *zfpm2b*), and rhombencephalon and spinal cord-associated expression signatures (*irx5a*/*rfx4*/*atoh1a*), indicating regionally patterned, state-diverse progenitor composition in the adult *Danionella cerebrum* CNS. Overall, region explained more of the observed transcriptomic variation than neurotransmitter class across the analyses of mature neurons we performed. Collectively, these data suggested that adult transcriptomic identity of neuronal types is governed more by region of origin than by neurotransmitter identity, mirroring the pattern observed in mammals, and suggesting a conserved organizing principle of the vertebrate nervous system. Cross-species analyses using SAMap also suggested that regional or developmental identity is a primary driver of similarity across species. This atlas also provides a resource of transcriptional changes associated with neural regeneration. Telencephalon regeneration data also revealed transcriptional changes in the uninjured hemisphere; it will be interesting to determine if this is caused by disrupted connections between hemispheres or a more systemic response.

The data presented here collectively provide a broad resource of molecularly distinct cell populations across all major tissues of the *Danionella cerebrum* body (this resource is accessible at digifish.wi.mit.edu/atlas). These findings and resources position *Danionella cerebrum* as a powerful new model for systems neuroscience, stem cell biology and regeneration, biomechanics and collective behavior, and evolutionary biology.

## Materials and Methods

### Animal care and husbandry

Adult *Danionella cerebrum* were maintained in 3, 6, and 9-L tanks (Iwaki Aquatic Systems) at a density of 9-12 animals/liter, with a ratio of ∼2:1 male to female. The water chemistry conditions were as follows: pH, 6.9-7.2; conductivity, 150-350 µS/cm; and temperature, 25-27 °C. Animals were kept in a 12:12-hour circadian cycle under indirect light. We provided 4-5 cm-long, clear and dark blue silicon tubes for breeding, as well as artificial grass/leaves/rocks for tank enrichment. Larvae were maintained at 28 °C in an incubator or on a slide heater and were fed a pure paramecium culture for 15 days in 10cm glass dishes (Pyrex), with daily changes of E3 medium (Cold Spring Harbor Zebrafish E3 medium protocol, without methylene blue). After 15 days, the animals were placed in a water circulation system and fed 2-3 times a day with either a live Artemia culture or Gemma Micro 150 (Skretting, USA). Eggs were collected 1-2 times a day, usually within 2 hours following morning and early afternoon feedings.

### Tissue dissection and single-cell isolation

*Danionella* tissues were dissociated using either the Worthington Papain Dissociation System (for nervous system dissociations; Product No: LK003150) or Liberase-based (Sigma Aldrich, Product No: NC1179175) enzymatic digestion (non-CNS tissues). Nervous system samples were dissected under a dissection microscope using fine forceps. The telencephalon was separated from the diencephalon and mesencephalon with a perpendicular cut anterior to the diencephalon and mesencephalon. The rhombencephalon was separated from the diencephalon/mesencephalon with a perpendicular cut just posterior to the optic tectum, dorsoventrally separating the rhombencephalon from the mesencephalic area. The spinal cords were isolated by gently pulling them, with the help of the forceps, from the anterior part of the trunk, following the removal of the brain, and were combined with rhombencephalon samples. Media and enzyme solutions were prepared following the manufacturer’s guidelines. Specifically, Neurobasal (Gibco, Product No: 21103049) (or Earle’s Balanced Salt Solution, EBSS; Worthington Papain Dissociation System) was supplemented with B27 (Invitrogen, Product no: A3582801) (2%: 500µl B27 per 25ml medium) and was kept on ice. The papain solution was prepared by adding 5ml of Neurobasal (or EBSS) to Vial 2 of the Worthington Papain Dissociation System. DNAse was prepared by resuspending the contents of Vial 3 in 0.5ml of Neurobasal Medium, of which 250µl was transferred to the papain vial. The papain/DNAse mixture was gently inverted 5-7 times to mix and incubated at 32-34 °C in a water bath during tissue collection.

For tissue dissections, 1ml of Dulbecco’s phosphate-buffered saline (DPBS) containing calcium and magnesium (Gibco, Product no: 14040117) was aliquoted into 1.5ml microcentrifuge tubes and kept on ice for subsequent tissue collections. Adult *Danionella cerebrum* (4-6 months old) were anesthetized using Tricaine-S (MS-222; 200mg/L; Syndel USA) and transferred to a 93mm light-blue Sylgard® dish (Living Systems Instrumentation, DD-90) containing 15-20ml of ice-cold Neurobasal/B27 (or EBSS for non-neural tissues). The target tissues or organs were microdissected under a stereomicroscope, and each sample was then minced into smaller fragments using fine forceps. For brain tissues, 20 animals were pooled per sample or brain region. For non-neural tissues, 5-7 animals were sufficient to yield adequate cell numbers.

Tissue fragments were first washed by brief centrifugation in DPBS (with calcium and magnesium) and the supernatant was removed. The pellet was gently resuspended in 1ml ice-cold DPBS (without calcium or magnesium; Gibco, Product No: 14190144), and was briefly held at room temperature (3-5 minutes). 1ml of pre-warmed enzyme mixture (papain/DNase or Liberase/DNase) was added to each sample. For papain-mediated dissociation, tissues were incubated at 34 °C for 20 minutes (or up to 30 minutes for non-neural tissues). Tissues were then gently triturated using a P1000 micropipette set to 900µl. A second incubation at 34 °C for 5-7 minutes was performed, followed by an additional 15-25 gentle triturations. Incubation times were extended up to 10-15 minutes if visible tissue fragments were still present. For Liberase-based dissociations, incubation was performed at room temperature for 30 minutes rather than at 34 °C.

Following enzymatic digestion, cell suspensions were centrifuged at 500 x g for 5 minutes at 4 °C in a swinging-bucket rotor. During centrifugation, the protease inhibitor cocktail was prepared by adding 32ml of EBSS to the ovomucoid inhibitor vial (Vial 4, Worthington Papain Dissociation System). An inhibitor mix was then prepared by mixing 900µl of EBSS with 100µl of ovomucoid solution, and 50µl of DNase in a 1.5ml LoBind (Eppendorf) microcentrifuge tube. Supernatant containing the enzyme solution was gently removed, and the cell pellet was resuspended in 1ml of the ovomucoid inhibitor solution. Samples were centrifuged again at 500 x g for 5 minutes at 4°C, and the resulting pellet was gently resuspended in 1ml of ice-cold DPBS (without calcium or magnesium), supplemented with 2-3% (% w/v) ultrapure bovine serum albumin (Invitrogen; BSA; Product no: AM2618). All subsequent washes and resuspensions were performed in this BSA-supplemented DPBS buffer. Cell suspensions were filtered through a 40-µm cell strainer into pre-chilled LoBind tubes, followed by centrifugation at 500 x g for 5 minutes at 4 °C. If residual debris was detected, up to two additional washes were performed, noting that each wash may lead to a substantial reduction in total cell yield. The final supernatant was discarded, and the final pellet was resuspended in 200-500µl of ice-cold DPBS (without calcium or magnesium). The resulting single-cell suspensions were kept on ice until the subsequent experimental steps. Cell concentration and viability were assessed using the trypan blue exclusion method. Accordingly, samples with viability above 80-90% were deemed suitable for any downstream single-cell sequencing workflow.

For CNS tissues, three anatomical regions, *(i)* telencephalon, *(ii)* diencephalon and mesencephalon, and *(iii)* rhombencephalon and spinal cord, were pooled from 20 animals into three sample tubes. Each pooled sample was then split into four aliquots and loaded onto the same 10X Genomics chip for cell capture using the 10X v3.1 HT workflow. After GEM generation, all 12 CNS samples (A1–A12) underwent library preparation simultaneously. The resulting libraries were multiplexed and sequenced on the same NovaSeq S4 lane. A smaller-scale 10X run consisting of three samples (samples 1–3), with one sample representing each of the same CNS regions, was also performed using the standard 10x v3.1 workflow. In addition, two telencephalon samples were processed separately using the 10x v3.1 HT workflow as an independent replicate set (G1–G2); these samples were sequenced together with telencephalon regeneration samples. A similar strategy was applied to non-neural tissues. Samples B1–B16 were processed together on the same 10x v3.1 HT chip and included multiple tissue types. For some tissues, pooled samples were split into multiple aliquots using the same pooling and aliquoting strategy described above, whereas other tissues (i. e., fin samples) were processed individually. In a separate experiment, internal organ samples were processed as an independent replicate set (H1–H4), again using the same pooling and aliquoting strategy. These (H1–H4) libraries were sequenced separately from the B-series samples, with all four libraries multiplexed on the same sequencing lane.

### 10X single-cell mRNA sequencing

Cells were processed following Chromium Next GEM Single Cell 3’ HT Reagent Kit v3.1 Dual Index (with Gel Bead Kit PN-1000372; Chip M; Dual Index Kit TT Set A-PN-1000215) and 10X Genomics Chromium Controller, following standard manufacturer’s guidelines. Samples were sequenced on an Illumina NovaSeq 6000 (150 paired-end reads) across all lines. A scaffold FASTA file and GFF3 file for the *Danionella cerebrum* genome were generated by the Judkewitz Lab (*30, 36*). After converting the GFF3 to a GTF file via gffread v0.12.7 (*133*), sequencing reads were mapped using the 10x Genomics Cell Ranger v7.2.0 pipeline (*134*). Cell Ranger outputs were then passed to CellBender 0.3.0 (*135*) for ambient RNA correction via the remove-background command. CellBender parameter adjustments included: --fpr set to 0.3 for internal organ samples, --expected-cells set to the exact number of cells predicted by Cell Ranger for each individual sample, --total-droplets-included set to a manually estimated value by inspection of Cell Ranger’s barcode rank plot for each individual sample, and --learning-rate used the default value of 1e-4 for each individual sample but was halved continuously as long as the CellBender report suggested re-running with a reduced learning rate. Using Seurat v5.0.3 (*136*), quality control and downstream analyses were performed separately for the atlas dataset and for the regeneration timecourse dataset. Droplets without counts were removed from unfiltered CellBender outputs, followed by doublet identification via scDblFinder v1.12.0 (*137*) (dbr=0.08). Any cells with nFeature_RNA < 200, nFeature_RNA > 99.5% percentile of nFeaure_RNA, nCount_RNA < 400, or nCount_RNA > 10000 were filtered from each dataset prior to doublet removal. Remaining cells were assessed for nUMI, nGene, percent mitochondrial, and percent ribosomal transcript content, which were represented in violin plots. Using gene annotations, percent mitochondrial content was based on genes with descriptions including the term “mitochondrial”, and percent ribosomal content was based on genes with names matching a ribosomal gene regular expression (^RP[LS][1-9]+[0-9]*[A-Z0-9]+). Gene expression was log normalized per cell via NormalizeData and scaled via ScaleData. Cells were visualized using the uniform manifold approximation and projection (UMAP) algorithm. For each dataset, the number of dimensions used with RunPCA, RunUMAP, and FindNeighbors was determined using JackStraw with a p value cutoff of 0.05. Clusters were determined via FindClusters using the leiden algorithm. UMAP plots of gene expression were created using Seurat’s FeaturePlot function (order=T). Marker genes were identified via Seurat’s FindAllMarkers function (only.pos=T) and filtered by those with p_val_adj < 0.05. Average expression values for each cluster were obtained via Seurat’s PseudobulkExpression, scaled per gene across clusters via R’s base scale function (center=F), and visualized as heatmaps via ComplexHeatmap v2.14.0 (*138*). For every cluster, cells that came from independent library preparations were treated as replicates. Libraries that were sequenced together were treated as batches in DESeq2. DESeq2 was used to compare each specific cluster against all other cells combined, controlling for variation between sequencing batches. DESeq2 results were compared with the original Seurat (FindAllMarkers) results, retaining only genes that showed a strong, statistically significant difference in both tests (adjusted P-value < 0.05 and log2 fold-change > 0.5). Finally, the top 20 marker genes were taken for each cluster from the Seurat analysis (ranked by adjusted P value), and the percentage of marker genes successfully confirmed by DESeq2 for each specific cluster was calculated. This verification was performed on both the internal organs subset and the CNS subset. Accordingly, 71.13% of all filtered Seurat markers and 95.7% of the top 20 Seurat markers per cluster were validated for internal organs across all internal organ clusters. Additionally, 62.4% of all filtered Seurat markers and 85.4% of the top 20 Seurat markers per cluster were validated for the CNS dataset across all CNS clusters. We report the results of this analysis in supplemental table S1 (DESeq2 analysis tabs). Violin plots and dot plots were created using Seurat’s VlnPlot and DotPlot functions, respectively. Alluvial plots were created using ggalluvial v0.12.5 (*139*). To determine the optimal clustering resolution, we used clustree (*140*) to find the most stable clustering resolution, where there was minimal new assignment of cells to new clusters at increasing clustering resolutions. Data processing utilized dplyr v1.1.4 (*141*), tidyr v1.3.1 (*142*), and tibble v3.2.1(*143*).

### Comparative paedomorphic comparison analysis

Three categories of single-cell RNA-sequencing data were compared for paedomorphic state analyses: *i)* a query atlas of adult *Danionella*, *ii)* a larval zebrafish reference (*80*), and *iii)* an adult zebrafish reference assembled from published datasets (*45, 46, 81–89*). Larval zebrafish were represented by the ZMAP atlas spanning 0–5 days post-fertilization. The adult zebrafish reference was a panel of eleven independently generated and independently annotated datasets, each processed separately. Three of these adult datasets include notes relevant to the interpretation of the data: the whole-organism microwell-seq data (*89*) span embryonic and adult stages; from this resource, only the adult-annotated cells were used. The brain dataset (*46*) profiles the adult telencephalon specifically, rather than the whole adult brain. The skin dataset (*86*) single-nucleus combinatorial indexing (sci)-RNA-seq from zebrafish at the adult transition state (during squamation) rather than fully mature adults, and was supplied as a Monocle3 ‘cell_data_set’ that was converted to a Seurat object by extracting the raw counts matrix, re-labeling features from the object’s ‘gene_short_name’ field (with ‘make.unique’ applied to ties), transferring cell metadata and the existing UMAP, and log-normalizing. Four tissue-class objects (neural, connective, epithelial, and muscle) were used for comparisons. Orthologous zebrafish gene symbols were previously produced by one-to-one orthology mapping of the native identifiers to zebrafish symbols. When a comparison required gene symbols, feature names were matched by lowercasing and removing a trailing one-to-many orthology suffix. For the *Danionella* atlas, the marker dot plot was read primarily in native *Danionella cerebrum* (DC) transcript ID-identifier space, with a curated symbol-to-DC-identifier table. Larval and adult expression were represented by gene symbol throughout. All cells across all data sources carried a “harmonized” ‘canonical_celltype’ label, reconciled across datasets into a common vocabulary. For the adult panel, cell labels were matched to each Seurat object per dataset, scoped by dataset of origin, so that cell barcodes shared between independent 10X experiments could never be cross-assigned. Each ‘canonical_celltype’ was mapped to one of four tissue classes (neural, connective, epithelial, muscle) by a fixed lookup table for the “developmental-position” analysis. For the expression dotplots, cell-type labels were additionally collapsed into a smaller set of coarse cell-type buckets by a separate lookup table (i.e. all fibroblast subtypes were grouped into “fibroblasts”; amacrine cells and photoreceptors were grouped into “retinal neurons”), and each bucket was assigned to its tissue class.

### Developmental-position scoring (per-cell-type pseudobulks) and selection of stage-discriminative genes and per-cell correlation

For each cell type present in all three sources, a pseudobulk expression vector was computed per source by aggregating counts across that cell type’s cells. Atlas and larval pseudobulks were expressed as log2 counts-per-million, log2(CPM + 1). The adult pseudobulk was a rank-based consensus across the adult datasets contributing that cell type (the per-dataset gene-rank vectors were averaged), to prevent any single platform or library-depth regime in the adult panel from dominating. Because all subsequent cell-level comparisons use Spearman (rank) correlation, the scoring was not affected by the difference in scale between the log2-CPM atlas/larval pseudobulks and the rank-based adult pseudobulk. For each cell type, the genes most informative about larval-versus-adult identity were selected: the atlas, larval, and adult pseudobulks were each rank-transformed across genes. For every gene the absolute difference between its adult rank and its larval rank was computed and the up to 1000 genes with the largest such difference were kept. All correlation scorings for a given cell type was restricted to their discriminative gene set. Each cell was scored by two Spearman correlations that were computed over its cell type’s discriminative genes (the correlation of the cell’s expression vector with the larval pseudobulk and with the adult pseudobulk). Spearman correlation was used because it is robust to per-cell normalization and the pseudobulk scale differences. At most 500 cells per cell type per source were sampled (fixed random seed (‘set.seed(42)’) set immediately before each sampling step), and all *Danionella* cells of the relevant cell types were scored for the panels. For each cell type, two anchors were defined in the two-dimensional space (larval correlation, ⍴-larval; adult correlation, ⍴-adult): the larval anchor L = (L_x, L_y) as the median coordinate of the ZMAP reference cells, and the adult anchor A = (A_x, A_y) as the median coordinate of the adult reference cells. For a cell at P = (cor_larval, cor_adult), the “developmental position” was the normalized scalar projection of P onto the L to A axis, dev_position = [(P − L) · (A − L)] / ǁA − Lǁ², so that dev_position = 0 at the larval anchor, and is 1 at the adult anchor; values outside [0, 1] denoted cells that lie beyond the larval or the adult anchor along the axis. The orthogonal residual to the axis (perp_dist) and the axis length (LA_len = ǁA − Lǁ) were kept as quality metrics. Developmental position was therefore defined as a within-cell-type relative coordinate, in which each cell type uses its own discriminative genes and anchors.

### Cross-source expression dot plots (curated markers)

For each source, cells were grouped by coarse cell-type bucket within tissue class, and for every (bucket, gene) pair two statistics were computed from the counts matrix: the percentage of cells with non-zero counts (percent expressing), and the mean of log1p(counts-per-10,000) across cells (mean expression, where counts-per-10000 is the per-cell count divided by that cell’s total counts and multiplied by 10^4^). For Seurat v5 objects, count layers were joined before extraction. The adult value for each (bucket, gene) was pooled across the eleven adult datasets as a cell-count-weighted mean of the per-dataset percent-expressing and per-dataset mean-expression values, with per-bucket cell counts summed. Datasets were never concatenated, so a gene’s pooled value uses only the datasets that contain both that cell type and that gene. Because percent expressing is a scale-free fraction, it is robust to per-dataset normalization and platform differences and serves as the primary (dot size) encoding/metric for comparisons. Mean expression, which is only approximately comparable across datasets, is used for color encoding. Dot size encodes percent expression on a fixed-area scale from 0 to 100%. Dot color encodes mean expression z-scored within each (source, gene) –for each gene, expression values were centered and scaled across the buckets of a single source and set to [−2.5, 2.5], so that the relative across-cell-type pattern is comparable between sources despite absolute-scale differences; genes with fewer than two non-missing values or zero variance within a source were set to zero. For the marker panel, a gene was displayed only if it was detected in all three sources for that tissue class, and genes were ordered along the vertical axis by a fixed, coarsely developmental-stage rank (larval-state and progenitor markers at the top, adult-state markers towards the bottom). The full per-tissue/source/gene tables underlying dot plots, including the cell counts behind every dot, were recorded (table S6). Analyses were performed in R using Seurat, matrix, dplyr, tibble, ggplot2, and ggrepel packages.

### CNS Region and neurotransmitter class comparison analyses

Single-cell RNA-seq datasets used in this analysis were generated across three experiments: 10X v3.1 HT (samples A1-A12), where we pooled three regions (telencephalon; diencephalon + mesencephalon; and rhombencephalon + spinal cord) were split into four aliquots. A second 10X v3.1 HT run comprised two telencephalon samples (G1, G2); and a standard chemistry 10X run of one sample per region (sample 1, 2, 3; fig. S1) for the three regions was performed. Ambient RNA was removed using CellBender during initial quality control steps. Cells were retained, for this analysis, if they belonged to a neuronal cluster, mapped to one of the three CNS regions, and assigned a neurotransmitter (NT) label, ultimately yielding a dataset of ∼269,000 CNS neurons. Each cell in the dataset was assigned a “region” label from its sample of origin and an experiment label defining the three separate runs. NT class was assigned in two different ways: *i)* a cluster-derived label (glutamatergic, GABAergic and other-NT) was taken from the cluster NT identity labels (originally assigned during manual cluster annotations); *ii)* a marker-derived NT identity was calculated from canonical neurotransmitter synthesis/transporter genes (vesicular glutamate transporters for glutamatergic neurons; glutamate decarboxylase/vesicular GABA transporter for GABAergic neurons; glycinergic, cholinergic and monoaminergic marker sets for other-NT) using module scores. For analyses that required a definitive/discrete NT label, a confidence-based thresholded assignment was used.

To study the arrangement of neuronal “types” rather than individual cells, we followed a “metacell” strategy adapted from F. M. Krienen et. al, 2023 (*47*). Neurons were clustered within each region or NT compartment, and each resulting cluster was aggregated (summed counts and normalized) into a pseudobulk metacell, producing a total of 206 metacells, each with a defined NT class and region (e.g., telencephalon X glutamatergic). Metacells with <20 cells were excluded from the dendrogram-based analyses. A metacell was assigned to its NT class only if that confidence score exceeded the of the next-highest-scoring class by at least 0.5 standard deviations to ensure the assigned class was clearly separated from the next-most likely class. Metacells not meeting these criteria were classified as ambiguous and excluded. Highly variable genes (HVG; n = 3000) were selected across the metacells, top 100 PCs were computed, and metacells were hierarchically clustered (average linkage on Euclidean PC distance). The Pearson correlation matrix using HVGs was visualized, annotating the region, NT class and experiment. To determine clade composition, the dendrogram was cut at k=2-30 clades, and at each level/depth the mean clade purity for region and for the NT class was recorded (i.e., fraction of each clade that belong to its dominant NT-label or region). To test whether NT usage involves a broad co-expression program, for every gene, we computed its Spearman correlation across metacells to the vesicular glutamate transporters (*slc17a7a*, *slc17a6a*/*b*) and to a cholinergic marker (*chata*), and compared these to a background distribution of random gene-pair correlations. We also tested whether the two vesicular glutamate transporters share an expression program by testing whether the 120 genes most strongly correlated with *slc17a7a* (vGLUT1) were also correlated with *slc17a6a/b* (vGLUT2). Dendrogram robustness was assessed by recomputing the tree with a correlation-based distance and measuring its topological agreement with the PC-distance tree (cophenetic correlation and Baker’s gamma), with significance evaluated against a null distribution of 100 trees built from permuted PC scores.

### Variance partitioning (PERMANOVA) test

To quantify how much of the transcriptomic variation among metacells is attributable to region versus neurotransmitter, we used permutational multivariate analysis of variance. This analysis (‘vegan::adonis2’, 999 permutations, marginal terms) partitioned the metacell distance matrix (1 - Pearson on HVGs) into region, NT, and experiment components, to report each factor’s (region, NT, and experiment) unique R². NT was assigned in three ways as a sensitivity metric: the original cluster NT label, the marker-thresholded discrete label, and continuous *slc17a6/slc17a7* (vGLUT) and *gad1/gad2, slc32a1* (GAD/VGAT) module scores. To assess whether neurotransmitter class separated neuronal types more strongly in certain brain regions than others, we repeated the PERMANOVA within each region. A complementary per-gene variance decomposition was computed with ‘variancePartition’ over HVGs, using region, glutamatergic and GABAergic marker scores, and experiment as variables.

Since the region and experiment (chemistry) are partially correlated, cell-level analyses were performed on both uncorrected (no integration) PCA embedding and an embedding integrated across the three experiments with Harmony (batch key = experiment, never aliquot). For each embedding (uncorrected PCA and Harmony-integrated), we calculated a Louvain resolution sweep (0.05-3) scoring agreement between the de novo clusters (without reference to metadata) and region/NT/experiment labels by adjusted Rand index (ARI), to measure how well two partitions of the same cells agree (1 = identical groupings, 0 = chance-level agreement), correcting for the agreement expected by chance.

### Region versus NT-class pseudobulk correlation analyses

Neurons were aggregated into pseudobulk profiles by intersecting brain region and neurotransmitter class (one profile per region-NT combination, AggregateExpression), and each profile was log-normalized. Pairwise Pearson correlations were computed across profiles over the top 2000 HVGs. The resulting correlation matrix was hierarchically clustered and visualized. Separately, cells were partitioned by NT class and re-clustered de novo within each class (GABAergic, glutamatergic, other; Seurat FindClusters, resolution 2.0, on a within-subset experiment-integrated Harmony embedding); each cluster was scored for region specificity as the fraction of its cells in the dominant region. The reverse analysis partitioned cells by region and re-clustered within each (GABAergic and glutamatergic only), scoring NT specificity analogously. Score distributions per cluster are plotted as box plots.

To test for cross-experiment reproducibility, neurons were aggregated into pseudobulk profiles per experiment × region (minimum of 50 cells per profile; no integration applied), and the profiles were correlated (Pearson, over the top 2,000 highly variable genes). For the two experiments spanning all three regions (HT v3.1, samples A1–A12; standard v3.1, samples 1–3), each region’s profile in one chemistry was matched to its most-correlated profile in the other chemistry, and mean same-region correlations were compared with cross-region correlations. Profiles were also hierarchically clustered and annotated by region and experiment to assess whether they grouped by region or by experiment.

### Single-cell sex-prediction confidence analysis

To assign a predicted sex (female or male) to each cell, we trained a supervised Random Forest classifier on our integrated whole-body *Danionella* scRNA-seq dataset using the caret framework, following the strategy developed for the African turquoise killifish single-cell atlas by B. B. Teefy et. al., 2023 (*144*). Because *Danionella* samples were pooled prior to droplet capture, ground-truth sex labels were not available at the cell or library level and were instead derived computationally from the expression of conserved teleost germline markers. Experimentally, we pooled 20 fish (18 males, 2 females) for CNS samples and 5 fish (3 males, 2 females) for non-neural samples (the rest of the body); therefore, we expected a bias towards “male-score” in the CNS samples. To identify “anchor” cells, for each cell, per-cell module scores were computed with Seurat::AddModuleScore (nbin = 24, ctrl = 100) using two non-overlapping marker panels. The female panel comprised oocyte (*buc*, *gdf9*, *zar1*), egg-envelope (*zp2.3*, *zp3b*, *zp3a.2*, *zpcx*), and vitellogenin (*vtg1*, *vtg2*, *vtg3*) transcripts; the male panel comprised meiotic (*sycp3*, *sycp2*), piRNA-pathway (*piwil1*, *piwil2*), and testicular somatic (*amh*, *dmrt1*) transcripts. We deliberately excluded pan-germline genes (*ddx4*, *dazl*, *dnd1*) and estrogen receptors (*esr1*, *esr2a*, *esr2b*). Cells with a female module score above the 97th percentile of the score distribution and a negative male module score were assigned as “female anchors”. The same criterion was used to identify “male anchors”. Anchor localization on the integrated UMAP was visually confirmed to coincide with annotated gonadal cell populations. To construct the training set, anchor cells were downsampled to equalize female and male class sizes, then partitioned 80/20 into training and held-out test sets using caret::createDataPartition with class stratification. For feature selection, the top 5000 highly variable genes were computed from the training anchor subset using Seurat::FindVariableFeatures. From these, only genes detected in more than 5% of cells in at least 90% of non-gonadal clusters were kept, to ensure that the features could generalize to somatic cells in the atlas. Pan-germline genes and estrogen receptors were removed from the feature set to prevent the classifier from re-learning gonad-specific signatures. A Random Forest classifier was trained with caret::train using the ranger implementation (500 trees, impurity-based feature importance, tuneLength = 3), with 5-fold cross-validation (trainControl(method = “cv”, number = 5, classProbs = TRUE, summaryFunction = twoClassSummary)) and ROC AUC as the optimization metric. Model performance was evaluated on the held-out anchor test set. The trained model was applied to log-normalized expression of the feature genes for all cells in the atlas dataset, yielding a per-cell female probability (p_female). Cells were assigned a binary predicted_sex (female if p_female > 0.5, otherwise male) and a five-tier sex_confidence label (male_high: p_female < 0.2; male_low: 0.2–0.4; ambiguous: 0.4–0.6; female_low: 0.6–0.8; female_high: ≥0.8). Feature importance rankings were extracted using caret::varImp to identify the transcripts that most strongly contribute to somatic-sex discrimination in *Danionella*. All analyses were performed in R using Seurat, caret, ranger, and dplyr.

### Cross-species analysis

For our mouse cross-species analysis, we used all brain cell samples from our *Danionella cerebrum* atlas and cells from the mouse whole-brain transcriptomic cell type atlas (Allen Brain Cell Atlas). Using the 20250531-release manifest for the mouse brain atlas, all raw 10Xv3 samples were subsampled to include 25% of cells from each cluster. Using the GENCODE mouse vM23 GTF and GRCm38 (v98) FASTA, AGAT (*145*) was used to keep the longest isoform of each gene via agat_sp_keep_longest_isoform.pl and to extract the CDS from each longest isoform via agat_sp_extract_sequences.pl (“-t cds”). The same technique was applied to the *Danionella cerebrum* genome to obtain a CDS for each longest isoform. These CDS files and scRNA-seq datasets were used with SAMap v2.0.2 (*90*) (DEFAULT_N_PCS set to 50) and eggNOG-mapper v2.1.12 (*146*) (-m=diamond (*147*), --itype=CDS, and --target-orthologs=one2one) to find one to one orthologs. Species orthologs were obtained via SAMap’s convert_eggnog_to_homologs function (og_key= ‘eggNOG_OGs’, taxon=33213). The utilized taxon code is specified in the SAMap vignette as a bilaterian taxon code for obtaining orthologs. Features for all datasets were restricted to one-to-one orthologs and log-normalized per cell via Seurat’s NormalizeData function. Using average gene expression for each cluster obtained via Seurat’s PseudobulkExpression, values were normalized per gene across clusters by dividing by mean expression, and correlation tests were performed on clusters between species via Psych’s corr.test function (method=“sp”, adjust=“fdr”, alpha=0.01, ci=F). Correlation plots were created via ComplexHeatmap. UMAP plots (fig. S33A-C) were constructed using SAMap’s UMAP coordinates.

For the zebrafish cross-species analysis, we used all high-throughput telencephalon brain samples (A1, A2, A7, A8, G1, G2) from our *Danionella cerebrum* CNS dataset and cells from an adult zebrafish telencephalon dataset L. Anneser et. al., 2024 (*42*). To match the number of cells from the zebrafish dataset, our telencephalon samples were subsampled to include approximately 12% of cells from each cluster, keeping at least 20 cells from each cluster when possible. Using v4.3.2 of the Lawson Lab zebrafish transcriptome annotation (*148*) GTF and their GRCz11 zebrafish genome FASTA, AGAT was used as previously described to keep the longest isoform of each gene, and “--mrna” was used with agat_sp_extract_sequences.pl to extract the mRNA sequence from each isoform because the GTF originally used with the data contains few CDS attributes. The previously described methods were used to extract CDS regions for *Danionella cerebrum*, run SAMap (DEFAULT_N_PCS set to the default value, 100), subset features to orthologs, normalize data, perform correlation tests, and create plots.

### Monocle3 pseudotime analysis

Pseudotime values were calculated via Monocle3 v1.4.26 (*149*). Pseudotime gene expression plots used normalized gene expression values scaled via R’s base scale function (center=F), smoothed across pseudotime via mgcv’s gam function, rescaled per gene from 0 to 1, and then plotted using ggplot2 v3.5.0 (*150*). Pseudotime values were binned using the same number of bins as the number of existing Seurat clusters. For heatmaps of the top 10 markers per pseudotime bin, marker genes were identified for each pseudotime bin via Seurat’s FindAllMarkers function (only.pos=T), filtered to unique markers only, and sorted by p_val. Average expression values for each pseudotime bin were obtained via Seurat’s PseudobulkExpression, scaled per gene across pseudotime bins via R’s base scale function (center=F), and visualized as heatmaps via ComplexHeatmap.

### Brain injury experiments and regeneration timecourse analysis

Brain injuries were performed either using dissection pins (Living Systems Instrumentation, SKU: PIN-0.1MM and PIN-0.2MM) or a vacuum-guided tissue removal method. Specifically, animals were anesthetized with Tricaine-S (MS-222; 200 mg/L; Syndel USA) and transferred onto a wet pad under a dissection microscope. A 0.1 mm-diameter dissection pin was used to make a unilateral telencephalon injury starting at a position posterior to the olfactory bulb and extending posteriorly towards the habenula, reaching as deep as the ventral pallial area (∼200-250 µm). For tissue removal, a mouth pipette, or a vacuum manifold attached to a rubber tubing with a filter, and a pulled glass capillary, was used to apply suction unilaterally, removing parts of the dorsolateral and dorsomedial pallium, posteriorly. Animals were allowed to recover in a tank with system water for ∼1 hour following surgery, then returned to their home tank for further recovery. They were later euthanized and fixed at different time points for regeneration/single-cell RNA sequencing experiments.

To quantify differences in sample composition and percentages across clusters, we organized cell counts by cluster (seurat_cluster) and sample (orig.ident) and converted these values to within-cluster percentages. Each cluster was represented by its sample percentages, and clusters were ranked by the distinctiveness of their composition using Jensen-Shannon divergence (JSD) (base 2) between each cluster’s sample distribution and the global sample distribution across cells. Accordingly, high divergence indicated enrichment or reduction in specific samples in clusters. Statistical significance was evaluated by a permutation test in which sample-of-origin labels were randomly shuffled across all cells while holding cluster sizes fixed (2,000 permutations), recomputing each cluster’s JSD to the reference under each permutation to build a null distribution. One-sided empirical p-values were calculated as (number of permutations with JSD ≥ observed + 1) / (number of permutations + 1) and corrected for multiple testing across clusters using the Benjamini–Hochberg procedure. Because the large number of cells per cluster yields a narrow null distribution, the resulting p-values are floor-limited and were used only to confirm non-random structure, not to rank clusters. To quantify the precision of each JSD estimate, 95% confidence intervals were obtained by bootstrap resampling of cells within each cluster with replacement (2000 iterations), with cluster ranking based on JSD magnitude and these intervals. Cluster factor levels were reordered based on this ranking, where high JSD scores indicated the most divergent clusters (marked by asterisks) (Fig. 6E).

For telencephalon regeneration analyses (Fig. 6, fig. S43), to visualize transcriptional dynamics, we generated a heatmap of genes identified as positively differentially expressed markers across timepoints (FindAllMarkers, p_adj_<0.05). For each gene, average expression was calculated per timepoint and scaled across timepoints (SD-scaling, without centering); all timepoints and conditions are compared to one another. Based on the heatmap, we manually filtered genes with scaled average expression ≥ 2 scaled units in at least one timepoint to select the genes that drive the temporal dynamics. A curated subset of these genes based on functional significance was included in the heatmaps shown in the main figure (Fig. 6H, I), and a complete list of differentially expressed genes by timepoint is presented in table S9.

### EdU labeling

F-ara-EdU (Click Chemistry Tools) was delivered through a soaking method in system water at a 1.25 mg/mL concentration for 24 hours. Animals were then transferred to tanks containing system water, and daily water changes (20-30%) were performed throughout the experiment. Animals were then euthanized and fixed at different time points in 4% PFA, overnight, at 4 °C, with gentle shaking. Following three 15-minute-long washes in PBST, EdU was developed through Click-IT reaction using Alexa Flour Azide (Invitrogen).

### HCR-FISH

Animals were fixed for 30-45 minutes in 4% PFA, at room temperature, gently shaking. Specimens were stored in methanol at −20 °C. Fixed animals were processed for whole-mount HCR FISH through two separate protocols. In the first approach, for fin, organ, and CNS labeling, specimens were equilibrated to room temperature, rehydrated through PBST:methanol to PBST, and transferred to in situ baskets. 500 µL buffer was used for washes, and 250 µL for pre-hybridization, hybridization, and amplification steps. The samples were permeabilized with Proteinase K for 10 minutes at room temperature, washed in PBST, and optically cleared in pre-warmed Deep Clear solution 1 (*151*) at 37 °C for 1 hour, followed by three additional PBST washes. Samples were then pre-hybridized in warm hybridization buffer for 30 minutes at 37 °C, and incubated >20 hours at 37 °C in hybridization solution (Molecular Instruments), containing probe oligos (1 pmol of each probe per 100 µL total). In a second approach, primarily used to process brain samples, animals were fixed in 4% PFA for about 24 hours, washed in PBST, and permeabilized in 1X PBS with 1% Triton-X-100 with 0.2% SDS for about 48 hours at 25 °C, in Eppendorf tubes. Samples were later transferred into 2X SSCT washes, hybridized at 37 °C for about 20 hours with the probe mix (1 µL probe stock per 250 µL hybridization solution per tube). Following hybridization, both protocols used 5 washes with pre-warmed wash buffer (Molecular Instruments) at 37 °C, followed by SSCT washes at controlled room temperature (∼23-30 °C). For amplification, hairpins (h1-h5 for each amplifier) were prepared by incubation at 95 °C for ∼90 seconds, then allowed to cool in the dark for ∼10-30 minutes. They were then combined with fresh amplification buffer (4-5 µL per hairpin per sample volume) and incubated overnight in the dark at controlled room temperature. Lastly, samples were washed extensively in 5X SSCT and counterstained with DAPI for 1 hour at room temperature. Samples were either mounted immediately after a brief PBST rinse in ProLong Gold (Invitrogen, Product no: P36930) or were optically cleared to RIMS (EasyIndex, Life Canvas Technologies) for imaging.

### Microscopy

Stereo images were acquired using a Zeiss Discovery V8 stereomicroscope with an AxioCam HRC camera. Fluorescence images were acquired using a Leica TCS SP8 confocal microscope. ImageJ (Fiji), Pixelmator, AxioVision, and ZEN (Zeiss) digital imaging software were used for processing and quantification of all HCR-FISH images.

### MERFISH

Adult individuals were fixed as described above. After fixation, samples were washed twice with 1x PBS prior to embedding in O.C.T. (Sakura Tissue-Tek) using a 10×10mm plastic cryomold (Sakura Tissue-Tek) and frozen using dry ice. Tissue blocks were then acclimated to −20 °C in a cryostat (Leica CM3050S) for at least 30 minutes prior to sectioning at a thickness of 16µm. Sections were collected on MERSCOPE slides (Vizgen) and utilized immediately for downstream experiments.

Sections were prepared according to the MERSCOPE sample preparation guidelines. In brief, samples were fixed in 4% paraformaldehyde for 15 minutes, washed three times in 1x PBS for 5 minutes each, and then placed overnight in 70% ethanol (v/v) at 4 °C. The following day, samples were incubated in Decrosslinking Buffer (Vizgen) for 5 minutes at 90 °C and cooled on the benchtop for 5 minutes. Samples were then washed twice with 5mL Conditioning Buffer (Vizgen) for 1 minute each, followed by incubation with Conditioning Buffer (Vizgen) for 30 minutes at 37 °C. Conditioning Buffer was then aspirated and replaced with 100uL of Pre-Anchoring Reaction Buffer (100uL Conditioning Buffer (Vizgen), 5uL Pre-Anchoring Activator (Vizgen), 5uL RNase Inhibitor Murine (NEB) covered by a 2×2 cm piece of parafilm. Samples were sealed with parafilm and placed in a humidified incubator at 37 °C for 2 hours. Samples were then washed with 5mL of Sample Prep Wash Buffer (Vizgen), followed by incubation in 5mL of Formamide Wash Buffer (Vizgen) for 30 minutes at 37 °C. The Formamide Wash Buffer was then aspirated from the sample and replaced with 100µL of Anchoring Buffer (Vizgen), and the sample was covered with a 2×2 cm piece of parafilm. Samples were sealed with parafilm and incubated overnight in a 37 °C humidified incubator. Following incubation, samples were washed in 5mL of Formamide Wash Buffer for 15 minutes at 47 °C, and then washed once with 5mL of Sample Prep Wash Buffer for 2 minutes at room temperature. Samples were then gel-embedded using the following protocol. A Gel Coverslip (Vizgen) was prepared by cleaning with RNaseZap (Invitrogen) and 70% ethanol (v/v). 100µL of Gel Slick (Lonza Bioscience) was then added for 10 minutes at room temperature. Samples were then incubated in a gel embedding solution (5mL Gel Embedding Premix (Vizgen), 25µL of 10% ammonium persulfate (w/v; Thermo Fisher), 2.5µL of N,N,N’,N’-tetramethylethylenediamine (Sigma Aldrich)) for 1 minute, followed by embedding with 50µL of gel embedding solution for 1.5 hours at room temperature. The Gel Coverslip was placed onto the sample to enable polymerization. Following gel embedding, the Gel Coverslip was carefully removed, and samples were cleared using the following protocol. Samples were digested for 6 hours in 200µL, covered with parafilm, at 37 °C using a Digestion Mix (200µL Digestion Premix (Vizgen) and 5µL RNase Inhibitor Murine (NEB)). Samples were then cleared in a solution of 1.25mL Clearing Premix (Vizgen) and 120µL Proteinase K (NEB) for 4 hours at 47 °C in a humidified incubator. 3.75mL of Clearing Premix was added to each sample and incubated overnight at 47 °C in a humidified incubator, keeping the total incubation time below 24 hours. Samples were then transferred to a 37 °C incubator for a maximum of 3 days.

Following clearing, samples were quenched for autofluorescence using a MERSCOPE Photobleacher (Vizgen) for a minimum of 3 hours. Following quenching, samples were washed 3x with 5mL of Sample Prep Wash Buffer for 5 minutes each at room temperature. Samples were then incubated in 5mL of Formamide Wash Buffer at 37 °C for 30 minutes. Following incubation, samples were carefully aspirated to remove the Formamide Wash Buffer and then incubated with 100µL of the corresponding MERSCOPE gene panel (Vizgen) covered in parafilm at 37 °C for 42 hours in a humidified incubator. Following hybridization, samples were washed twice with 5mL of Formamide Wash Buffer at 47 °C for 30 minutes each, and were then utilized immediately for MERSCOPE imaging, following standard protocols (Vizgen). Raw MERFISH data were processed using the MERSCOPE platform. Images were processed using MERSCOPE Visualizer.

## Acknowledgments

We thank the Reddien Lab for lively discussions and for critical input on the data. Specifically, we thank M. Lucila Scimone and Conor McMann for their constructive comments and discussions on the manuscript. We thank the Adam Douglass Lab for providing *Danionella cerebrum* larvae, which enabled the initial establishment of a *Danionella* colony in the laboratory. We thank Stephen J. Fleming for crucial discussions on the use of CellBender, and Ralf Britz and Kevin W. Conway for valuable discussions and their insights into the biology of *Danionella cerebrum.* We are grateful to George Bell and Troy Whitfield of the Whitehead Institute Bioinformatics and Research Computing Center for computational assistance; to the Whitehead Institute Genome Technology Core for sequencing-related expertise; and to the Whitehead Institute Information Technology Department for technical assistance. We also thank Andy Nutter-Upham and Scott McCallum from the Whitehead Institute Information Technology Department for their expertise in building the interactive *Danionella cerebrum* atlas website. Finally, we thank the *Danionella* community for key input and open discussions on newly developing protocols and ideas that enable the study of this emerging experimental model system.

## Author Contributions

KDA and PWR conceptualized the study, performed experiments, and wrote the manuscript with input from PA and BJ. KDA performed all surgical procedures with help from PWR, developed cell dissociation protocols, performed cell dissociations, ran 10X Chromium experiments, and prepared 10X single-cell sequencing libraries. PA and KDA implemented the computational quality-control pipeline for data analysis. KDA and PA performed all computational and statistical analyses. CP and KDA performed MERFISH experiments and imaging. MK and KDA optimized and performed all HCR experiments and microscopy. BJ and MK generated and provided WT and *NeuroD1*:*GCaMP6f Danionella cerebrum* strains, and an updated *Danionella cerebrum* genome resource along with transcript/gene annotations. OP and KDA optimized and refined husbandry and housing conditions for *Danionella cerebrum*.

## Data availability

The *Danionella cerebrum* single-cell RNA-seq data generated in this study have been deposited in the Sequence Read Archive (SRA) under accession number PRJNA1402358. The code used for the analysis of single-cell RNA-seq data and the processed data are available on Zenodo: https://zenodo.org/records/18236172. The single-cell RNA-seq and MERFISH datasets generated during this study can be interactively explored through the companion website at: digifish.wi.mit.edu/atlas.

---

**Table 1.** Markers for all datasets

**Table 2.** MERFISH gene list

**Table 3.** Cluster annotations for all datasets

**Table 4.** Average expression values and z-scores for heatmaps shown in fig S18

**Table 5.** Differentially expressed genes for anteroposterior body positions and fins

**Table 6.** Paedomorphic state comparison analysis

**Table 7.** Differentially expressed genes across CNS regions

**Table 8.** Cross-species analyses

**Table 9.** Telencephalon regeneration time course markers

**Table 10.** Quality control analyses

## Supplementary Materials

Supplementary Text

Figs. S1 to S44

Tables S1 to S10

