## Supplemental Figures for "Whole-body single-cell atlas of an adult vertebrate in homeostasis and regeneration"

### Supplemental Text

#### Molecular analysis of paedomorphic features

We explored paedomorphic gene expression signatures across connective, epithelial, and muscle tissues. For connective tissue, including blood, our analyses revealed expression of embryonic hemoglobin variants *hbbe2* and *hbae1.3* in adult *Danionella* (Fig. 3M). Similarly, we observed strong expression of *matn4* and *col2a1b*, genes typically expressed during development in chondrocytic components of the connective tissue (*152, 153*) (Fig. 3M; fig. S21), indicating a predominant cartilage-formation program. *col10a1a*, a gene required for osteogenesis (*154*), was expressed prominently (cluster 8), whereas *bglap*, marking late-stage osteoblasts and bone mineralization (*155*), was expressed relatively lower (Fig. 3M, N; fig. S21).

To assess the presence of potential paedomorphic gene expression signatures in epithelial tissues, we first generated a panel of genes with expression in basal, stratified, simple, periderm-like, ion-transport, and secretory epithelial states. To test for the persistence of juvenile-like simple epithelia and/or mature, fully differentiated adult basal/stratified epithelia, we selected a panel of markers that are candidates to indicate these states (Fig. 3M, fig. S21). We reasoned that widespread distribution of juvenile-like gene expression patterns could indicate a paedomorphic state for a given tissue class, whereas a fully differentiated gene expression program would suggest an adult-like state. We selected *epcam*, *cldn7b*, and *cdh* as core epithelial identity markers; these encode a pan-epithelial marker, a protein responsible for epidermal integrity by forming tight junctions, and lastly, a protein responsible for cadherin-based integrity of epithelial tissues, respectively (*156-158*). We included other basal keratinocyte markers, *tp63*, *krt5*, and *krtt1c19e* (*159-161*), as well as genes responsible for specialized functions such as formation of hemidesmosomal anchorage (a process that anchors epithelial cells to the underlying basement membrane through specialized junctions), like *itga4b*, *itga6b*, and *col17a1a* (*162*). As indicators of other specialized cell states, such as a mucus-secreting program (e.g., mature goblet cells), we included *agr2*, and, for ion transport, we added *slc4a4a* and *atp1a1.3* (*163, 164*). Because larval and juvenile epithelia in many teleosts exhibit simple-epithelium cytokeratin programs, these genes might provide a readout for a paedomorphic state in the adult. To identify potential larval/juvenile-like broad expression patterns, we examined expression of *krt4*, *krt8*, and *krt18a.1*, which encode simple cytokeratins (*159*). We observed broad co-expression of *krt4* and *krt8*, with *krt18a.1* expressed in multiple clusters (Fig. 3M; fig. S21). Specifically, *krt8* and *krt18a.1*, which are broadly expressed in larval and juvenile stages as well as in simple epithelia in the adult state (*165, 166*), were expressed in the oral mucosa, in cranial epithelial tissue, and in the gills of adult *Danionella*. Gill lamellae expressed *tp63*, a marker for basal epithelial progenitors, along with *epcam*, *krt8*, consistent with the simple squamous architecture of the lamellar respiratory epithelium (Fig. 3M) (*167*). Additionally, the lack of scales and significant pigmentation patterns suggest paedomorphic features in epithelial cells. By contrast, the brain vasculature displayed an adult-like gene expression signature with marked expression of *mfsd2aa*, *cldn5*, *slc2a1a*, *abcb4*, and *slc7a5*, indicating a mature-state blood-brain barrier program for cells of cluster 3 (fig. S21) (*168, 169*). Cluster 45 had strong *dll4* (arterial fate marker), *cxcr4*, and *cav1* expression, along with some *krt8* expression (Fig. 3M, fig. S21), but not expression of *krt4* or *krt18a.1*, demonstrating a mature large-vessel/arterial program (*170*).

Internal organs showed that the exocrine pancreas (marked by *ptf1a* and *cpb1*), liver (*apoba*, *abcb11b*, and *fabp10a*), intestine (*fabp6* and *slc20a1a*), and kidney (*wt1b* and *pax2a*) generally displayed previously established adult-like gene expression signatures (*171-174*), (Fig. 3M, fig. S21). A muscle gene panel showed lower expression for genes that are primarily developmentally active or enriched during larval/juvenile states (*mhyz2* and *mhyz1.2*), indicating an adult-like signature for this tissue (Fig. 3M, O; fig. S21).

Thyroid hormone (TH) can act as a systemic tissue-maturation or metamorphosis cue, coordinating maturation or tissue specialization transitions with tissue-selective effects, since a collection of factors (e.g., deiodinases, transporters, receptors) linked to thyroid signaling can be differentially expressed across cell types (*175-177*). It was previously shown that *Danionella* possesses functional thyroid follicles and intact T4 production, and that paedomorphosis in *Danionella* is not because of impaired thyroid hormone synthesis (*178*). In parallel, brain states associated with social dynamics, such as collective behavior, were also linked to thyroid signaling-related gene programs (*26, 30*). Consistent with these findings, our analysis with key TH signaling-related genes showed broad expression of thyroid hormone receptors, *thraa* and *thrb*; a CNS-biased expression pattern for T4 entry regulated by *slco1c1*, a T4 transporter; but potentially limited local T4 to T3 activation (regulated by *dio2*) and weak expression of *klf9*, a direct target of thyroid signaling (*179*) (with telencephalon showing slightly higher expression compared to other brain samples), and relatively low expression of metabolic effectors (*cpt1aa* and *pdk4*) of TH in neural tissues. Non-neural tissues showed relatively stronger expression of *dio1*, *klf9,* and *ucp2*, suggesting a tissue-type-biased effect that could be investigated in the future to assess their potential contribution to systems-level heterochronic effects (fig. S22).

**Telencephalon regeneration**

scRNA-seq at time points during regeneration revealed 53 clusters of cells (127,168 total cells), which generally represented the overall cellular complexity of the telencephalon, including radial glia, intermediate progenitors, different neuronal and non-neuronal cell classes, as well as injury-specific cell clusters/states (Fig. 6C, fig. S42G, table S9). DGE analysis indicated that the majority of global gene expression changes in response to injury occurred between 12 hours and 7 days post-injury (Fig. 6D). Beyond the inherent asymmetrical gene expression differences between the two sides of the telencephalon, there also existed extensive transcriptional changes on the uninjured side of the telencephalon. Analysis of cluster composition showed that clusters 39, 47, 49, 50, and 53 were enriched for cells from specific regeneration time points (Fig. 6E). These clusters showed cellular states associated with an inflammatory response and myeloid cell recruitment, as well as radial glia proliferation and neurogenesis, and neuronal maturation (Fig. 6E, F; fig. 42H, I). Examining gene expression signatures of the most highly expressed genes across regeneration-specific clusters further revealed a wound-response program. Specifically, clusters 47 and 50 contained acute inflammation-related genes that control leukocyte recruitment and microglia activation, including *ptprc*, *ccr9a*, *csf1ra*, *lect2l (180, 181)*; a separate cluster, 39, expressed radial glia proliferation and microglia-related genes (*fabp7a*, *ptgdsb.1*, *apoc1*).

A detailed timepoint-specific analysis confirmed transcriptional signatures of a wound response, stem cell proliferation, neurogenesis, and neuronal maturation cascade during regeneration (Fig. 6H-K, fig. S42J, table S9). We examined gene expression signatures enriched for each time point in both the injured and contralateral, uninjured telencephalon. We observed similarities between the injured and uninjured sides and signatures unique to injury, as well as signatures unique to the uninjured side, when compared with control animals with no injury (i.e., no injury on either side). Damage-associated chemokines and cytokines (*cxcl18b* and *il11a*) and immune modulators linked to monocyte and macrophage recruitment and inflammation (*tnfrsf11b*, *cd9b*, *s100a10b*, and *rnasekb*) indicated a wound response program (*182-185*). Factors linked with endoplasmic reticulum proteostasis, secretory and endolysosomal trafficking (*p4hb*, *SMCO3*, *cst14a.2*, *atp6v1g1*, and *apoeb*) (*186-189*); genes associated with protein synthesis (*rps19*/*10a*, *rpl19*, *rpc3*) and metabolic activation (*spag7*, *aldoaa*, and *mt-nd5*) (*190-192*); and genes linked to activity-dependent neuronal remodeling (*cdk5r2a*) (*193*) suggested neuronal remodeling program. The contralateral, uninjured telencephalon lacked the myeloid cell recruitment signature (i.e., no enrichment for *cxcl18b*, *il11a*, *s100a10b*, *apoeb*, *cd9b*), but showed a systemic response linked to metabolic activation and stress response-related genes (*p4hb*, *calr3a/b*, *cst14a.2*, *mt-nd5*, *skg2b*) (*192, 194*), along with *tnfrsf11b*, a gene linked to cytokine regulation (*195*) (Fig. 6H, I).

Day 3 injured telencephalon-enriched gene expression signatures suggested a state shift from an acute immune response to regulated immune modulation, with microglia- and macrophage-related genes *cxcl11.1*, *Fcer1g*, *cd84*, *cd74a*, and *Csf1r* expressed along with *lect2l*, indicating a leukocyte-modulation response (*182, 196, 197*) (Fig. 6H). We also observed upregulation of genes related to lipid-clearing dynamics and efferocytosis, such as *fabp11a*, *apoc1*, and *anxa1a* (*198-200*), and genes linked to barrier formation and regenerative response, *Cldnb* (claudin b) and *epd*, expressed by ependymal and meningeal cells (*201-204*). Concurrently, expression of *fabp7a*, a key radial glia marker, indicated a neural progenitor proliferation/neurogenesis response in the injured telencephalon (*12, 46, 106, 114*). On the contralateral, uninjured side, expression of stress-modulatory genes *mxi1* and *DUSP8* suggested the existence of a systemic compensatory state (*205-207*). The upregulation of *abca1*, *hyal4*, and *napgb*, genes together previously linked with membrane lipid flux, synaptic vesicle turnover, and extracellular matrix stability, suggested a metabolic state that differed from the inflammatory or neurogenic mode observed on the injured side at this time point (*208-210*). We also observed upregulation of *gsx2*, a gene implicated in the developmental patterning of the zebrafish forebrain (*211*) and in an injury response following damage in the mouse neocortex (*212*). We detected upregulation of *bmf2*, a BH3-only protein and pro-apoptotic member of the Bcl-2 family, uniquely expressed in the uninjured, contralateral telencephalon on day 3 following injury. Bmf was previously shown to be activated in response to ECM-detachment stress in neurons (*213, 214*). At Days 7 and 14 post-injury, both the injured and the uninjured contralateral telencephalon showed broad neuronal structural remodeling and an early neuronal maturation program. Specifically, the Day 7 expression profile revealed genes associated with the Notch and FGF pathways (*dla*, *notch1b*, *fgfr2*, *spry2* and *eya2*) that have been previously implicated in promotion of radial glial identity (*215*); progenitor and glial cell type markers (*msi1*, *gsx2*, *slc1a3a*, and *cx43*) (*211, 212, 216-219*); and genes linked with metabolic regulation (*aldob*, *bhmt*, *hpdb*, and *abca1b*). Both *cx43* and *bhmt* have previously been implicated in the regeneration of the zebrafish fin and liver (*220-222*). Day 30 after unilateral telencephalic injury showed upregulated circuit maturation and neuromodulatory signatures for both sides of the telencephalon. We detected *gng8* and *kiss1* expression on the injured side at this time point, which may suggest the re-establishment of modulatory coupling across the long-range and habenular axes, respectively, and *chata* expression might indicate the regeneration and strengthening of cholinergic transmission as neural networks functionally integrate (Fig. 6H).

Finally, we generated pseudotime trajectories of regenerating neurons and focused on glutamatergic lineages (Fig. 6J). To elucidate the dynamics, we first built correlation matrices between regenerating and steady-state radial glia and progenitors, and then between regenerating and steady-state glutamatergic neurons (fig. S44). Using steady-state radial glia and intermediate progenitor cells, and glutamatergic neuron clusters that showed similarity to their regenerating counterparts, we constructed pseudotime trajectories that predominantly predicted dorsolateral pallium (*rgs14a*+) and olfactory glutamatergic projection neuron (*cntn5*+) gene expression signatures (Fig. 6J, fig. S44).

**Comparative analyses between *Danionella*, mouse and adult zebrafish telencephalon**
Comparing *Danionella cerebrum* and mouse neural clusters or classes primarily captured conserved regional and/or developmental transcriptional programs, and neurotransmitter identity was not the principal driver of cross-species cell-type similarity. In the glutamatergic compartment, telencephalic glutamatergic clusters from *Danionella* showed broad similarity to thalamic-habenular (diencephalic relay; 18-TH and 17-MH-LH) classes; to neuroendocrine-associated glutamatergic classes (16-HY and 15- HY-Gnrh1), along with a pineal class (25-Pineal), and to the cerebellar granule-like excitatory neuron class (29-CB). These similarities were primarily driven by the telencephalic glutamatergic clusters from the *Danionella cerebrum* CNS. In the GABAergic compartment, subsets of *Danionella* telencephalon neural clusters showed similarity primarily with subpallial and basal ganglia GABAergic classes (09-CNU-LGE, 07 CTX-MGE, and 08-CNU-MGE), as well as hindbrain/cerebellar GABAergic classes (28-CB) (Fig. 4K). The GABAergic clusters that showed similarity to mouse cell classes and supertypes spanned the telencephalon, diencephalon/mesencephalon, and rhombencephalon, consistent with a broader Medial Ganglionic Eminence/Lateral Ganglionic Eminence-like program deployed across multiple regions. Additionally, *Danionella* radial glia clusters showed strong similarity to mouse glial classes (30 Astro-Epen and 32-OEC) and were broadly anticorrelated with neuronal classes, indicating an ependymoglial/radial glial identity. *Danionella* radial glia clusters from multiple brain origins (25, 7, 14, 4, 1, and 28) showed broad correlation across mouse Astro-TE/OLF and astroependymal, tanycyte, and ependymal-like states. Region-specific clusters (i.e., 15, 17, 18) showed selective similarity to astroependymal/tanycyte and Bergmann glia programs (fig. S35). Separately, a comparison between different glutamatergic and GABAergic neurons across different regions of the *Danionella cerebrum* CNS revealed a stronger similarity between two different neurotransmitter classes from the same brain region, when compared to the same neurotransmitter class from different regions (Fig. 4J, fig. S32E). This trend was also reflected when mouse and *Danionella* neurons were compared using SAMap, with similarity mainly driven by regional or developmental origin (fig. S33E, table S8, assortativity scores

Fig S1.

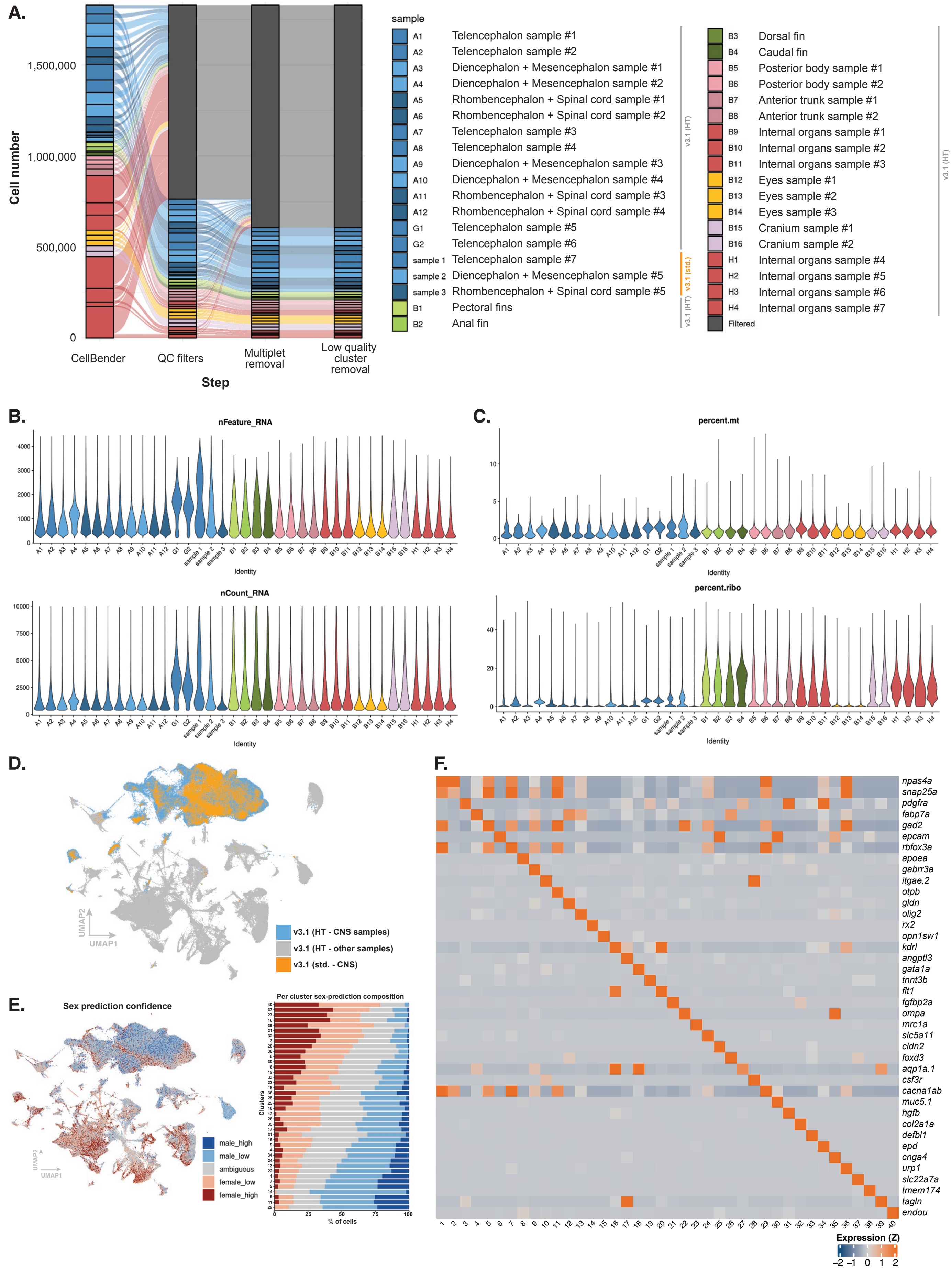

A.
Cell number
1,500,000
1,000,000
500,000
0
CellBender
QC filters
Multiplet removal
Low quality cluster removal
Step
sample
A1 Telencephalon sample #1
A2 Telencephalon sample #2
A3 Diencephalon + Mesencephalon sample #1
A4 Diencephalon + Mesencephalon sample #2
A5 Rhombencephalon + Spinal cord sample #1
A6 Rhombencephalon + Spinal cord sample #2
A7 Telencephalon sample #3
A8 Telencephalon sample #4
A9 Diencephalon + Mesencephalon sample #3
A10 Diencephalon + Mesencephalon sample #4
A11 Rhombencephalon + Spinal cord sample #3
A12 Rhombencephalon + Spinal cord sample #4
G1 Telencephalon sample #5
G2 Telencephalon sample #6
sample 1 Telencephalon sample #7
sample 2 Diencephalon + Mesencephalon sample #5
sample 3 Rhombencephalon + Spinal cord sample #5
B1 Pectoral fins
B2 Anal fin
v3.1 (HT)
v3.1 (std.)
v3.1 (HT)
B3 Dorsal fin
B4 Caudal fin
B5 Posterior body sample #1
B6 Posterior body sample #2
B7 Anterior trunk sample #1
B8 Anterior trunk sample #2
B9 Internal organs sample #1
B10 Internal organs sample #2
B11 Internal organs sample #3
B12 Eyes sample #1
B13 Eyes sample #2
B14 Eyes sample #3
B15 Cranium sample #1
B16 Cranium sample #2
H1 Internal organs sample #4
H2 Internal organs sample #5
H3 Internal organs sample #6
H4 Internal organs sample #7
Filtered
v3.1 (HT)
B.
nFeature_RNA
nCount_RNA
Identity
C.
percent.mt
percent.ribo
Identity
D.
UMAP2
UMAP1
v3.1 (HT - CNS samples)
v3.1 (HT - other samples)
v3.1 (std. - CNS)
E.
Sex prediction confidence
UMAP2
UMAP1
male_high
male_low
ambiguous
female_low
female_high
Per cluster sex-prediction composition
Clusters
% of cells
F.
npas4a
snap25a
pdgfra
fabp7a
gad2
epcam
rbfox3a
apoea
gabrr3a
itgae.2
otpb
gldn
olig2
rx2
opn1sw1
kdrl
angptl3
gata1a
tnnt3b
flt1
fgfbp2a
ompa
mrc1a
slc5a11
cldn2
foxd3
aqp1a.1
csf3r
cacna1ab
muc5.1
hgfb
col2a1a
defbl1
epd
cnga4
urp1
slc22a7a
tmem174
tagln
endou
Expression (Z)
−2 −1 0 1 2

**fig S1. General overview and quality control of the single-cell RNA-seq data (A)** Sankey plot showing computational quality control steps taken, the list of experimental sample replicates, and the 10X single-cell sequencing chemistry used to process the samples (table S10). **(B)** Violin plots showing the percentage of the number of uniquely expressed genes (nFeatures) and the percentage of the number of RNA molecules (UMIs) detected across all genes (nCounts) across all experimental samples and replicates. **(C)** Violin plots indicating the percentage of mitochondrial and ribosomal genes detected across all experimental samples and replicates. **(D)** UMAP plot of single-cells labeled by 10X chemistry, central nervous system samples showing homogenous mixing across chemistries. **(E)** Predicted sex composition across the Danionella whole-body single-cell atlas. CNS samples were generated by pooling 20 fish (18 males, 2 females) and body samples by pooling 5 fish (3 males, 2 females); the resulting sex skew is reflected in the predictions. A Random Forest classifier trained on transcriptionally unambiguous gonadal cells (following the strategy from B.B. Teefy et. al, 2023 (144)) was used to assign each cell a continuous female vs male probability score. UMAP colored by predicted sex for each cell and per-samples stacked bar plot of confidence tiers. **(F)** Selected top markers used to indicate the molecular identities of the main clusters of the whole-body atlas.

**Figure S2.**

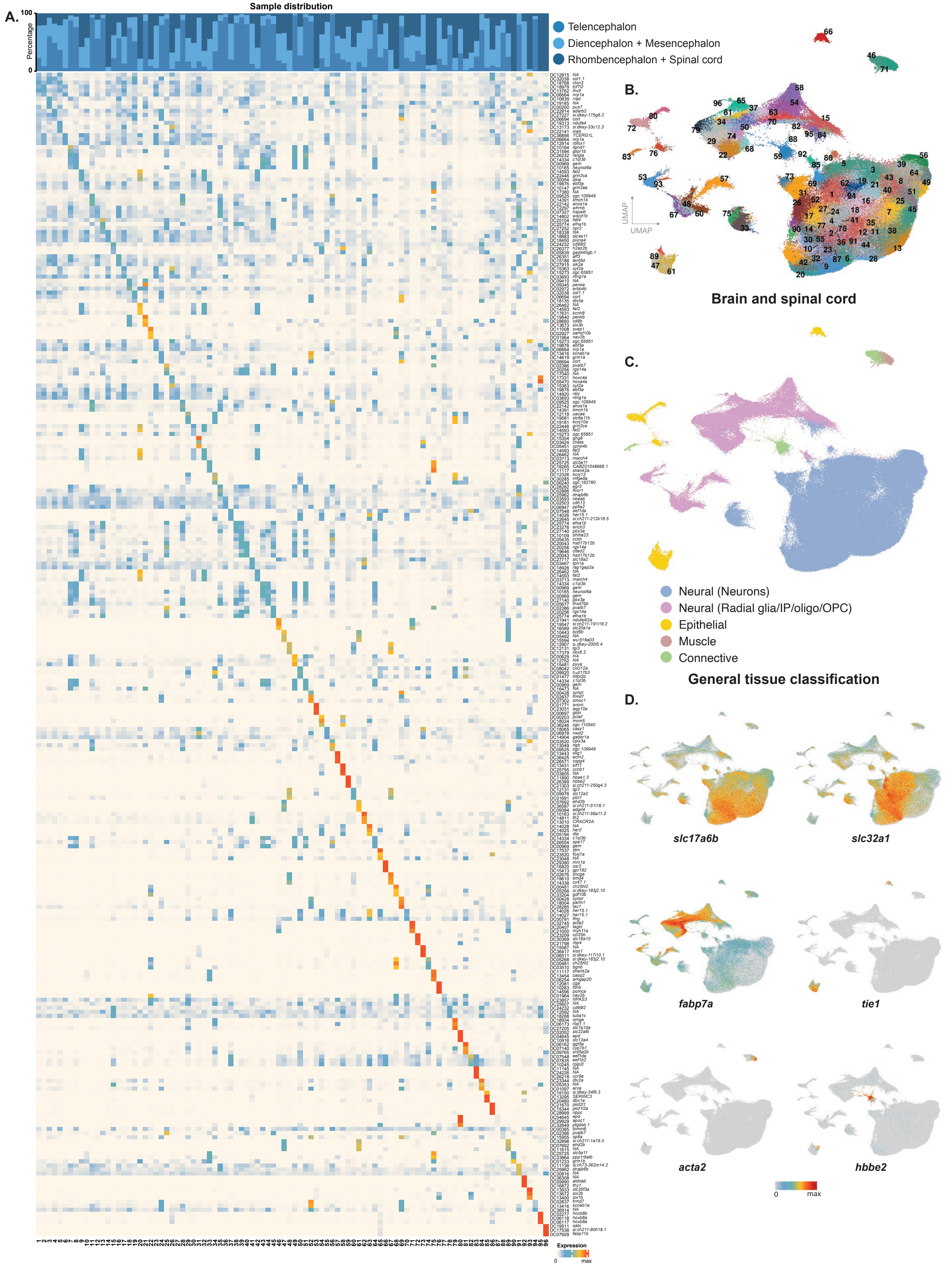

A.
Sample distribution
Percentage
100
0
Telencephalon
Diencephalon + Mesencephalon
Rhombencephalon + Spinal cord
Expression
0
max
B.
UMAP
UMAP
Brain and spinal cord
C.
Neural (Neurons)
Neural (Radial glia/IP/oligo/OPC)
Epithelial
Muscle
Connective
General tissue classification
D.
slc17a6b
slc32a1
fabp7a
tie1
acta2
hbbe2
0
max

**fig. S2. Characterization of brain and spinal cord single-cell RNA-seq data (A)** Top three markers and sample distributions for the central nervous system (CNS) clusters, including brain and spinal cord, but excluding eye samples, are shown. **(B)** UMAP representation of all CNS clusters. **(C)** UMAP plot showing distribution of neural, epithelial, connective, and muscle cells across the CNS samples. **(D)** Feature plots showing expression of marker genes for glutamatergic neurons (*slc17a6a*), GABAergic neurons (*slc32a1*), radial glia (*fabp7a*), endothelial cells (*tie1*), smooth muscle cells (*acta2*), and erythrocytes (*hbbe2*) across the CNS dataset.

**Figure S3.**

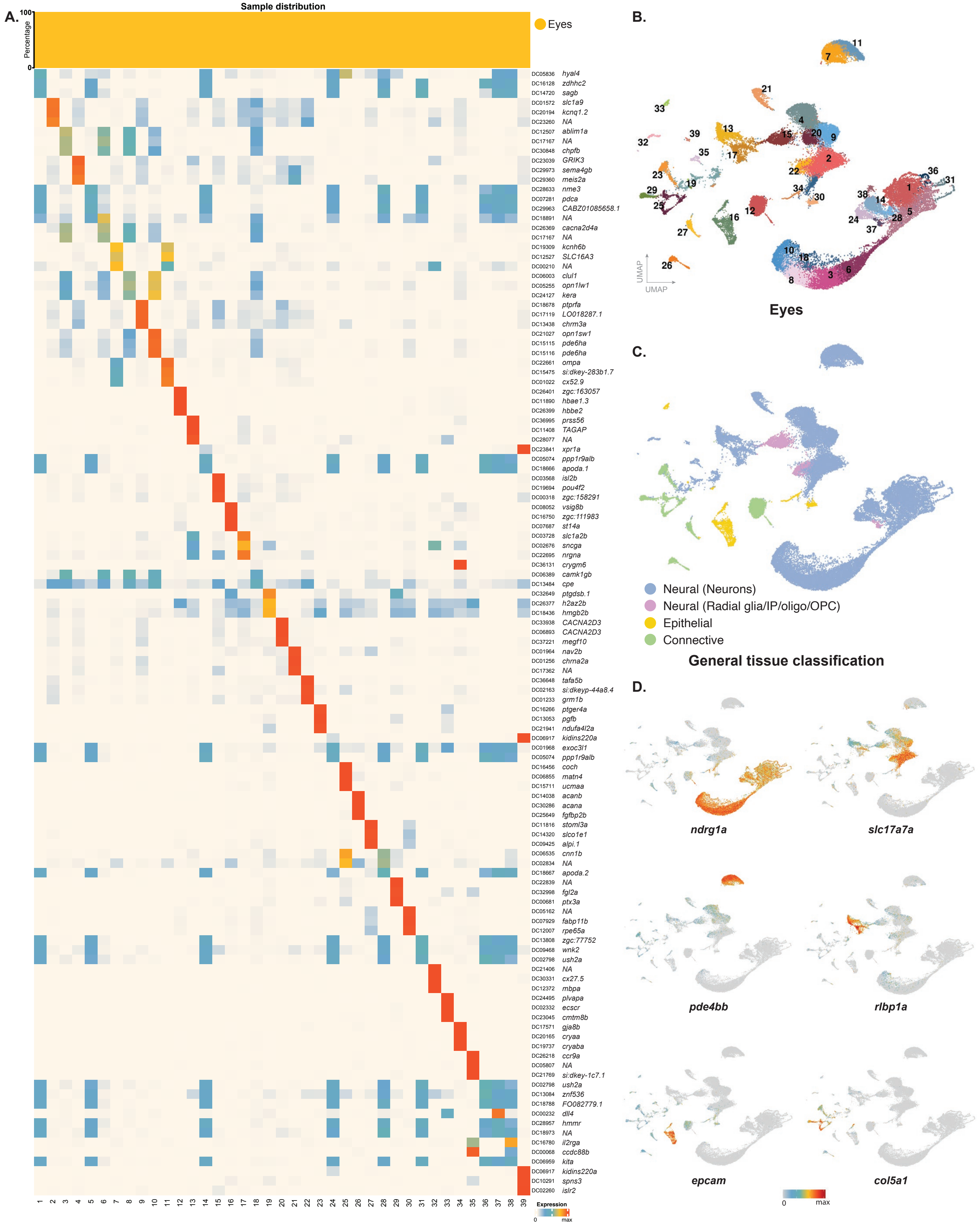
A.
Sample distribution
Percentage
Eyes
Expression
0
max
B.
UMAP
Eyes
C.
Neural (Neurons)
Neural (Radial glia/IP/oligo/OPC)
Epithelial
Connective
General tissue classification
D.
ndrg1a
slc17a7a
pde4bb
rlbp1a
epcam
col5a1
0
max

**fig. S3. Characterization of eye single-cell RNA-seq data (A)** Top three markers and sample distributions for eye clusters are shown. **(B)** UMAP representation of all eye clusters. **(C)** UMAP plot showing distribution of neural, epithelial, connective, and muscle cells across the eye samples. **(D)** Feature plots showing expression of marker genes for photoreceptor neurons (*ndrg1a*), retinal bipolar cells. (*slc17a7a*), retinal horizontal cells (*pde4bb*), simple epithelium (*epcam*), and fibroblasts (*col5a1*) across the eye dataset.

**Figure S4.**

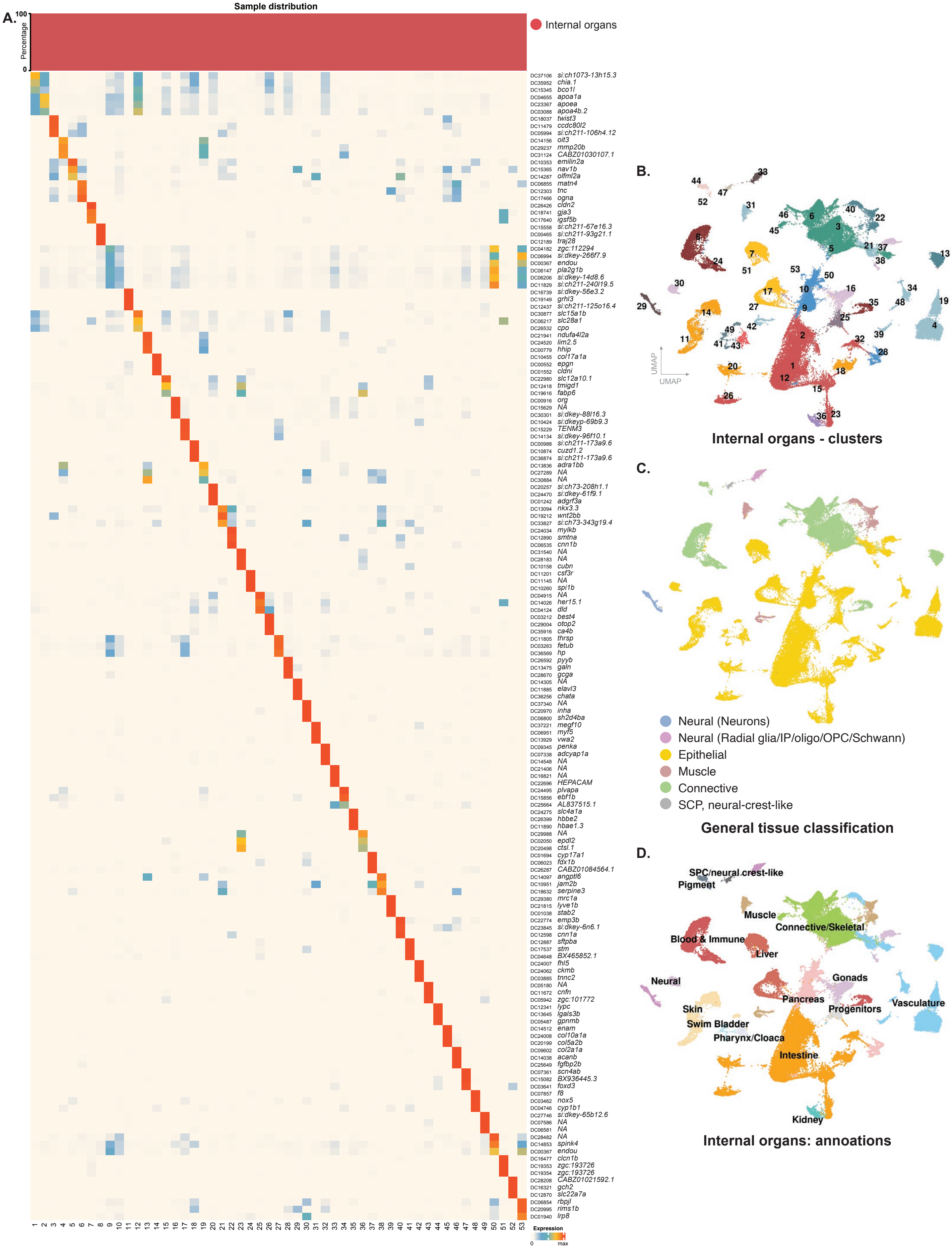
A.
Sample distribution
Percentage
100
0
Internal organs
Expression
0
max
B.
UMAP
UMAP
Internal organs - clusters
C.
Neural (Neurons)
Neural (Radial glia/IP/oligo/OPC/Schwann)
Epithelial
Muscle
Connective
SCP, neural-crest-like
General tissue classification
D.
SPC/neural crest-like
Pigment
Muscle
Connective/Skeletal
Blood & Immune
Liver
Neural
Gonads
Pancreas
Skin
Progenitors
Vasculature
Swim Bladder
Pharynx/Cloaca
Intestine
Kidney
Internal organs: annoations

**fig. S4. Characterization of internal organs single-cell RNA-seq data (A)** Top three markers and sample distributions for internal organs clusters are shown. **(B)** UMAP representation of all internal organ clusters. **(C)** UMAP plot showing distribution of neural, epithelial, connective, and muscle cells across the internal organ samples. **(D)** UMAP plot showing tissue/organ-level annotations within the internal organs dataset.

**Figure S5.**

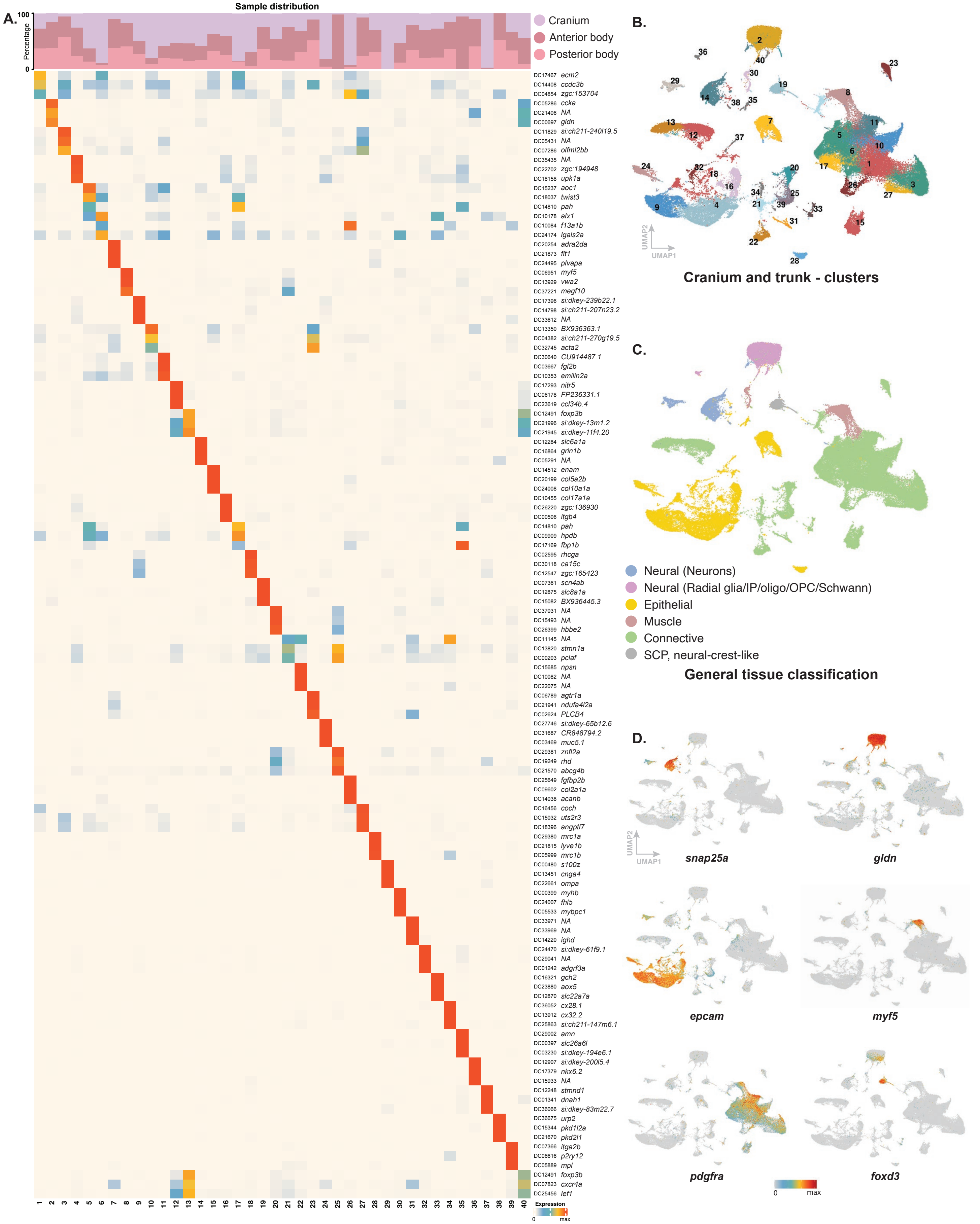

A.
Sample distribution
Percentage
100
0
Cranium
Anterior body
Posterior body
DC17467 ecm2
DC14408 ccdc3b
DC04854 zgc:153704
DC05286 ccka
DC21406 NA
DC00697 gldn
DC11829 si:ch211-240l19.5
DC05431 NA
DC07286 olfml2bb
DC35435 NA
DC22702 zgc:194948
DC18158 upk1a
DC15237 aoc1
DC18037 twist3
DC14810 pah
DC10178 alx1
DC10084 f13a1b
DC24174 lgals2a
DC20254 adra2da
DC21873 flt1
DC24495 plvapa
DC06951 myf5
DC13929 vwa2
DC37221 megf10
DC17396 si:dkey-239b22.1
DC14798 si:ch211-207n23.2
DC33612 NA
DC13350 BX936363.1
DC04382 si:ch211-270g19.5
DC32745 acta2
DC30640 CU914487.1
DC03667 fgl2b
DC10353 emilin2a
DC17293 nitr5
DC06178 FP236331.1
DC23619 ccl34b.4
DC12491 foxp3b
DC21996 si:dkey-13m1.2
DC21945 si:dkey-11f4.20
DC12284 slc6a1a
DC16864 grin1b
DC05291 NA
DC14512 enam
DC20199 col5a2b
DC24008 col10a1a
DC10455 col17a1a
DC26220 zgc:136930
DC00506 itgb4
DC14810 pah
DC09909 hpdb
DC17169 fbp1b
DC02595 rhcga
DC30118 ca15c
DC12547 zgc:165423
DC07361 scn4ab
DC12875 slc8a1a
DC15082 BX936445.3
DC37031 NA
DC15493 NA
DC26399 hbbe2
DC11145 NA
DC13820 stmn1a
DC00203 pclaf
DC15685 npsn
DC10082 NA
DC22075 NA
DC06789 agtr1a
DC21941 ndufa4l2a
DC02624 PLCB4
DC27746 si:dkey-65b12.6
DC31687 CR848794.2
DC03469 muc5.1
DC29381 znfl2a
DC19249 rhd
DC21570 abcg4b
DC25649 fgfbp2b
DC09602 col2a1a
DC14038 acanb
DC16456 coch
DC15032 uts2r3
DC18396 angptl7
DC29380 mrc1a
DC21815 lyve1b
DC05999 mrc1b
DC00480 s100z
DC13451 cnga4
DC22661 ompa
DC00399 myhb
DC24007 fhl5
DC05533 mybpc1
DC33971 NA
DC33969 NA
DC14220 ighd
DC24470 si:dkey-61f9.1
DC29041 NA
DC01242 adgrf3a
DC16321 gch2
DC23880 aox5
DC12870 slc22a7a
DC36052 cx28.1
DC13912 cx32.2
DC25863 si:ch211-147m6.1
DC29002 amn
DC00397 slc26a6l
DC03230 si:dkey-194e6.1
DC12907 si:dkey-200l5.4
DC17379 nkx6.2
DC15933 NA
DC12248 stmnd1
DC01341 dnah1
DC36066 si:dkey-83m22.7
DC36675 urp2
DC15344 pkd1l2a
DC21670 pkd2l1
DC07366 itga2b
DC06616 p2ry12
DC05889 mpl
DC12491 foxp3b
DC07823 cxcr4a
DC25456 lef1
1 2 3 4 5 6 7 8 9 10 11 12 13 14 15 16 17 18 19 20 21 22 23 24 25 26 27 28 29 30 31 32 33 34 35 36 37 38 39 40
Expression
0
max
B.
UMAP2
UMAP1
Cranium and trunk - clusters
C.
Neural (Neurons)
Neural (Radial glia/IP/oligo/OPC/Schwann)
Epithelial
Muscle
Connective
SCP, neural-crest-like
General tissue classification
D.
snap25a
gldn
epcam
myf5
pdgfra
foxd3
0
max

**fig. S5. Characterization of cranium and trunk single-cell RNA-seq data (A)** Top three markers and sample distributions for cranium and trunk clusters are shown. **(B)** UMAP representation of all cranium and trunk clusters. **(C)** UMAP plot showing distribution of neural, epithelial, connective, and muscle cells across cranium and trunk samples. **(D)** Feature plots showing expression of marker genes for mature neurons (*snap25a*), Schwann cells (gldn), simple epithelial cells (*epcam*), myoblasts (*myf5*), fibroblasts (*pdgfra*), and Schwann cell precursors and neural-crest-like cells (*foxd3*) across the cranium and trunk dataset.

**Figure S6.**

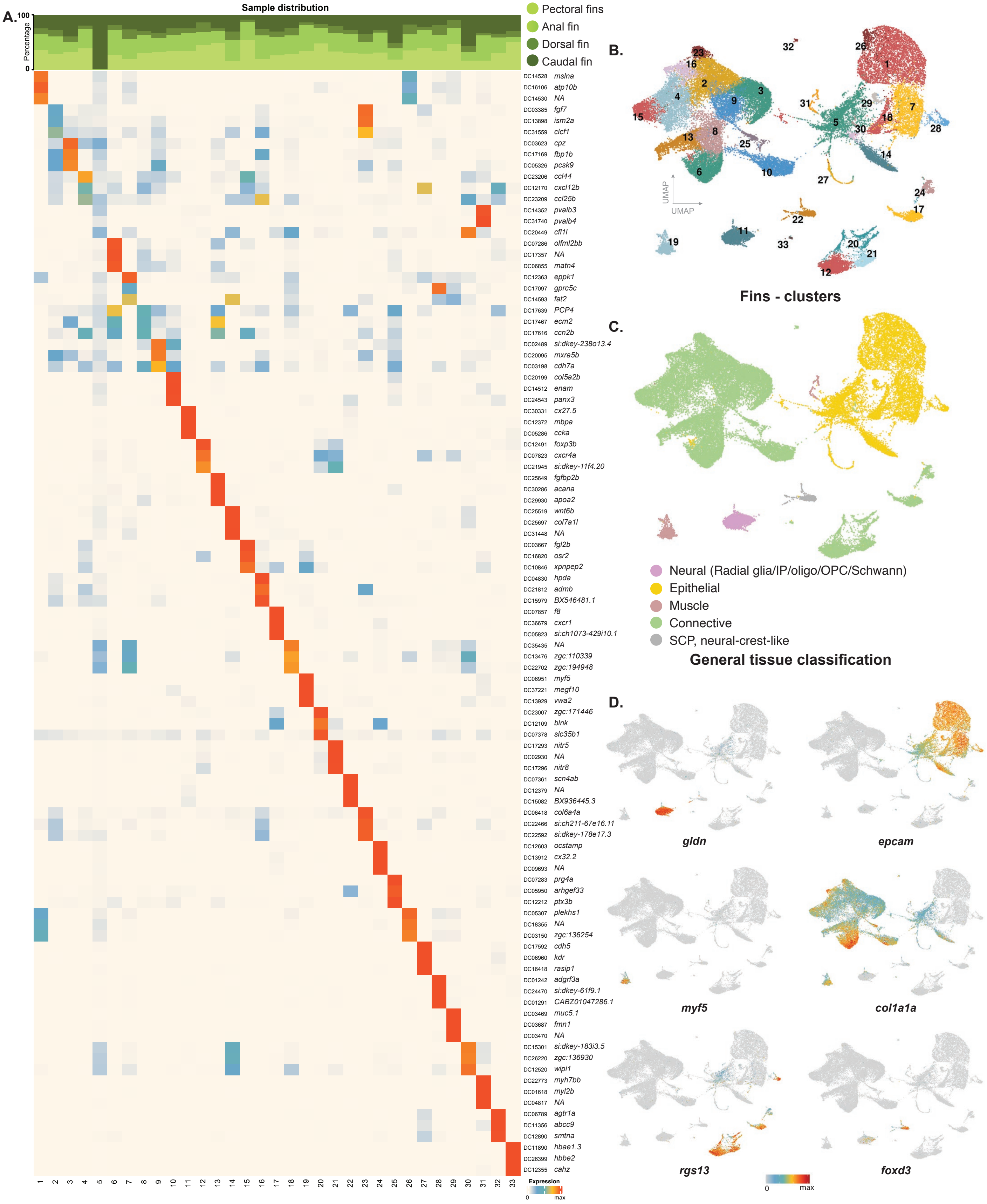

A.
Sample distribution
Percentage
100
0
Pectoral fins
Anal fin
Dorsal fin
Caudal fin
mslna
atp10b
NA
fgf7
ism2a
clcf1
cpz
fbp1b
pcsk9
ccl44
cxcl12b
ccl25b
pvalb3
pvalb4
cfl1l
olfml2bb
NA
matn4
eppk1
gprc5c
fat2
PCP4
ecm2
ccn2b
si:dkey-238o13.4
mxra5b
cdh7a
col5a2b
enam
panx3
cx27.5
mbpa
ccka
foxp3b
cxcr4a
si:dkey-11f4.20
fgfbp2b
acana
apoa2
wnt6b
col7a1l
NA
fgl2b
osr2
xpnpep2
hpda
admb
BX546481.1
f8
cxcr1
si:ch1073-429i10.1
NA
zgc:110339
zgc:194948
myf5
megf10
vwa2
zgc:171446
blnk
slc35b1
nitr5
NA
nitr8
scn4ab
NA
BX936445.3
col6a4a
si:ch211-67e16.11
si:dkey-178e17.3
ocstamp
cx32.2
NA
prg4a
arhgef33
ptx3b
plekhs1
NA
zgc:136254
cdh5
kdr
rasip1
adgrf3a
si:dkey-61f9.1
CABZ01047286.1
muc5.1
fmn1
NA
si:dkey-183i3.5
zgc:136930
wipi1
myh7bb
myl2b
NA
agtr1a
abcc9
smtna
hbae1.3
hbbe2
cahz
1 2 3 4 5 6 7 8 9 10 11 12 13 14 15 16 17 18 19 20 21 22 23 24 25 26 27 28 29 30 31 32 33
Expression
0
max
B.
UMAP
UMAP
Fins - clusters
C.
Neural (Radial glia/IP/oligo/OPC/Schwann)
Epithelial
Muscle
Connective
SCP, neural-crest-like
General tissue classification
D.
gldn
epcam
myf5
col1a1a
rgs13
foxd3
0
max

**fig. S6. Characterization of fin single-cell RNA-seq data (A)** Top three markers and sample distributions for fin clusters are shown. **(B)** UMAP representation of all fin clusters. **(C)** UMAP plot showing distribution of neural, epithelial, connective, and muscle cells across fin samples. **(D)** Feature plots showing expression of marker genes for Schwann cells (*gldn*), simple epithelial cells (epcam), myoblasts (*myf5*), connective tissue cells, including fibroblasts and cartilage cells (*col1a1a*), immune cells (*rgs13*), and Schwann cell precursors and neural-crest-like cells (*foxd3*) across the fin dataset.

**Figure S7.**

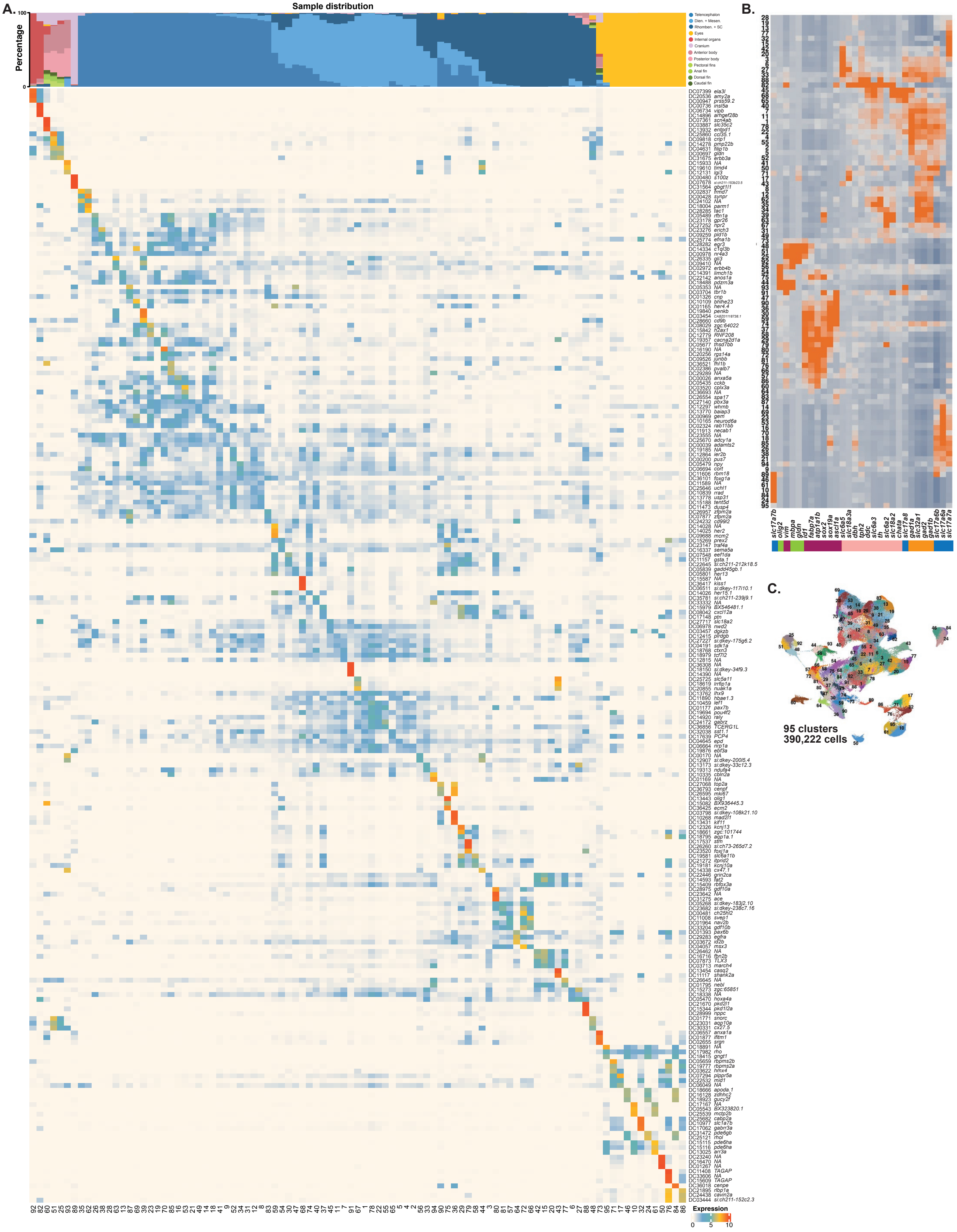
A.
Sample distribution
Percentage
Telencephalon
Dien. + Mesen.
Rhomben. + SC
Eyes
Internal organs
Cranium
Anterior body
Posterior body
Pectoral fins
Anal fin
Dorsal fin
Caudal fin
Expression
B.
C.
95 clusters
390,222 cells

**fig. S7. Characterization of neural (CNS and PNS) single-cell RNA-seq data (A)** Top three markers and sample distributions for all neural clusters are shown. **(B)** Heatmap showing the expression of a curated marker gene set used to identify cell classes shown in Fig 4 A-C. **(C)** UMAP plot showing 95 molecularly distinct neural cell clusters representing neural cell types from across the whole body.

**Figure S8.**

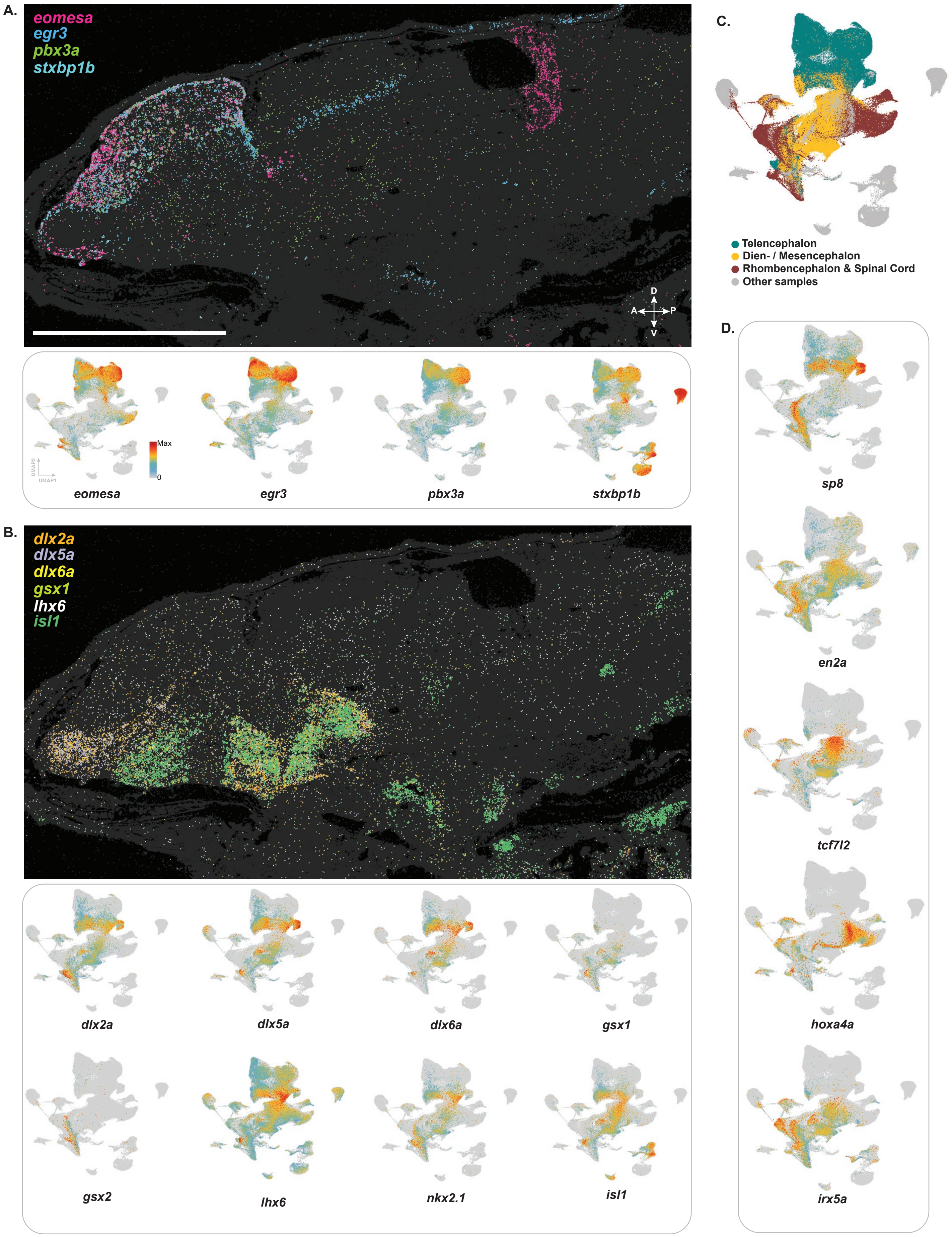
A.
eomesa
egr3
pbx3a
stxbp1b
D
A
P
V
Max
0
UMAP2
UMAP1
eomesa
egr3
pbx3a
stxbp1b
B.
dlx2a
dlx5a
dlx6a
gsx1
lhx6
isl1
dlx2a
dlx5a
dlx6a
gsx1
gsx2
lhx6
nkx2.1
isl1
C.
Telencephalon
Dien- / Mesencephalon
Rhombencephalon & Spinal Cord
Other samples
D.
sp8
en2a
tcf7l2
hoxa4a
irx5a

**fig. S8. MERFISH analysis of regionally expressed genes in the brain (A)** MERFISH and Feature plots showing expression of genes primarily expressed in the dorsal pallium. **(B)** MERFISH and Feature plots showing expression of genes primarily expressed in the ventral pallium. **(C)** UMAP showing the distribution of cells from the telencephalon, diencephalon, mesencephalon, rhombencephalon, and spinal cord across the all neural cells dataset, including the central and peripheral nervous systems. **(D)** Feature plots showing gene expression patterns enriched across distinct anatomical zones.

Figure S9.

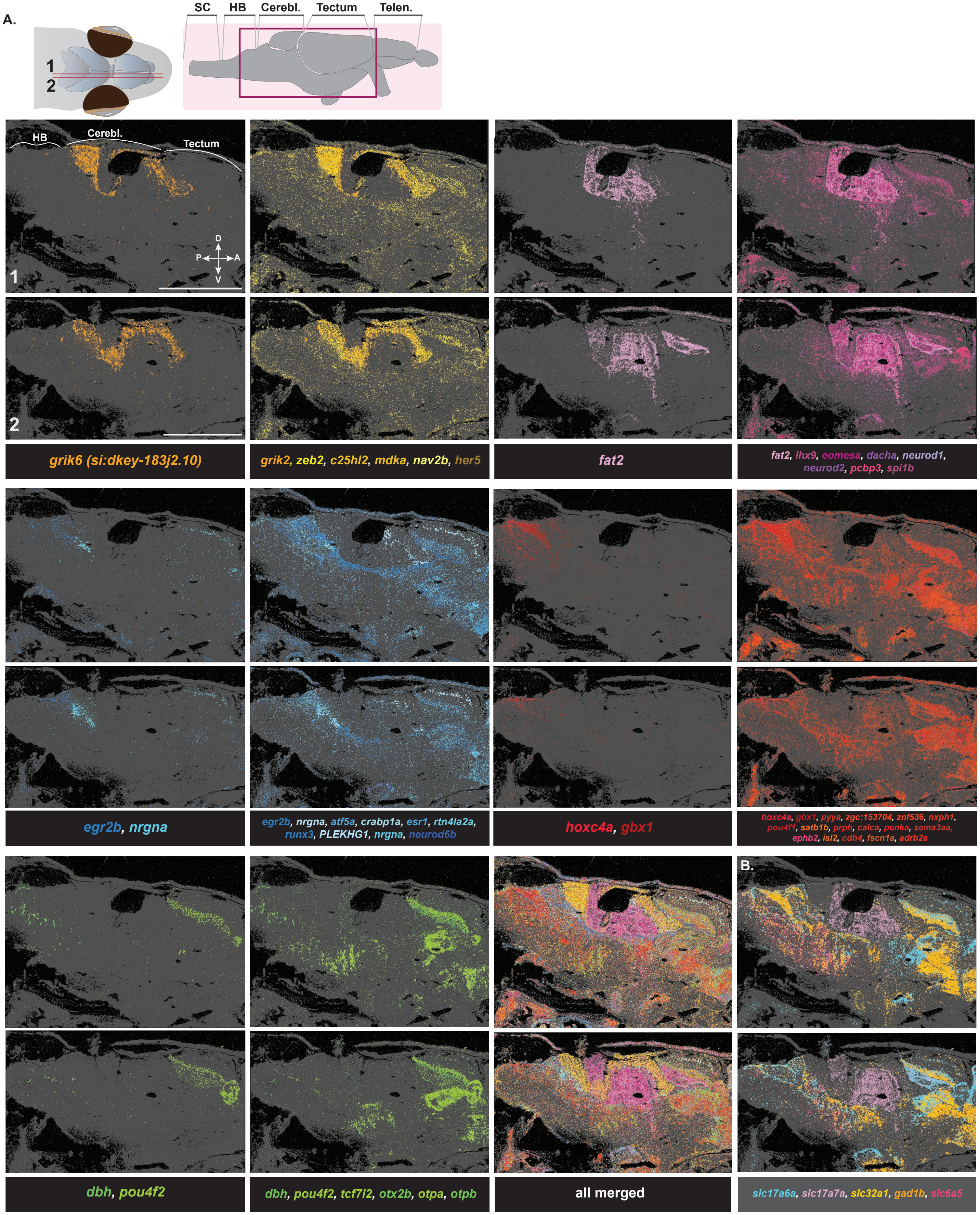
A.
1
2
SC
HB
Cerebl.
Tectum
Telen.
HB
Cerebl.
Tectum
D
P
A
V
1
2
grik6 (si:dkey-183j2.10)
grik2, zeb2, c25hl2, mdka, nav2b, her5
fat2
fat2, lhx9, eomesa, dacha, neurod1, neurod2, pcbp3, spi1b
egr2b, nrgna
egr2b, nrgna, atf5a, crabp1a, esr1, rtn4la2a, runx3, PLEKHG1, nrgna, neurod6b
hoxc4a, gbx1
hoxc4a, gbx1, pyya, zgc:153704, znf536, nxph1, pou4f1, satb1b, prph, calca, penka, sema3aa, ephb2, isl2, cdh4, fscn1a, adrb2a
B.
dbh, pou4f2
dbh, pou4f2, tcf7l2, otx2b, otpa, otpb
all merged
slc17a6a, slc17a7a, slc32a1, gad1b, slc6a5

**fig. S9. MERFISH analysis of regionally expressed genes in the mid and hindbrain regions (A)** MERFISH panels from two separate mediolateral sections showing expression of genes with regional expression patterns in cerebellum, optic tectum, and hindbrain. **(B)** MERFISH panel showing expression of glutamatergic (*slc17a6a, scl17a7a*), GABAergic (*slc32a1a, gad1b*) and histaminergic (*slc6a5a*) neuron markers. Scale bar: 1mm; SC: spinal cord, HB: hindbrain, Cerebl: cerebellum.

Figure S10.

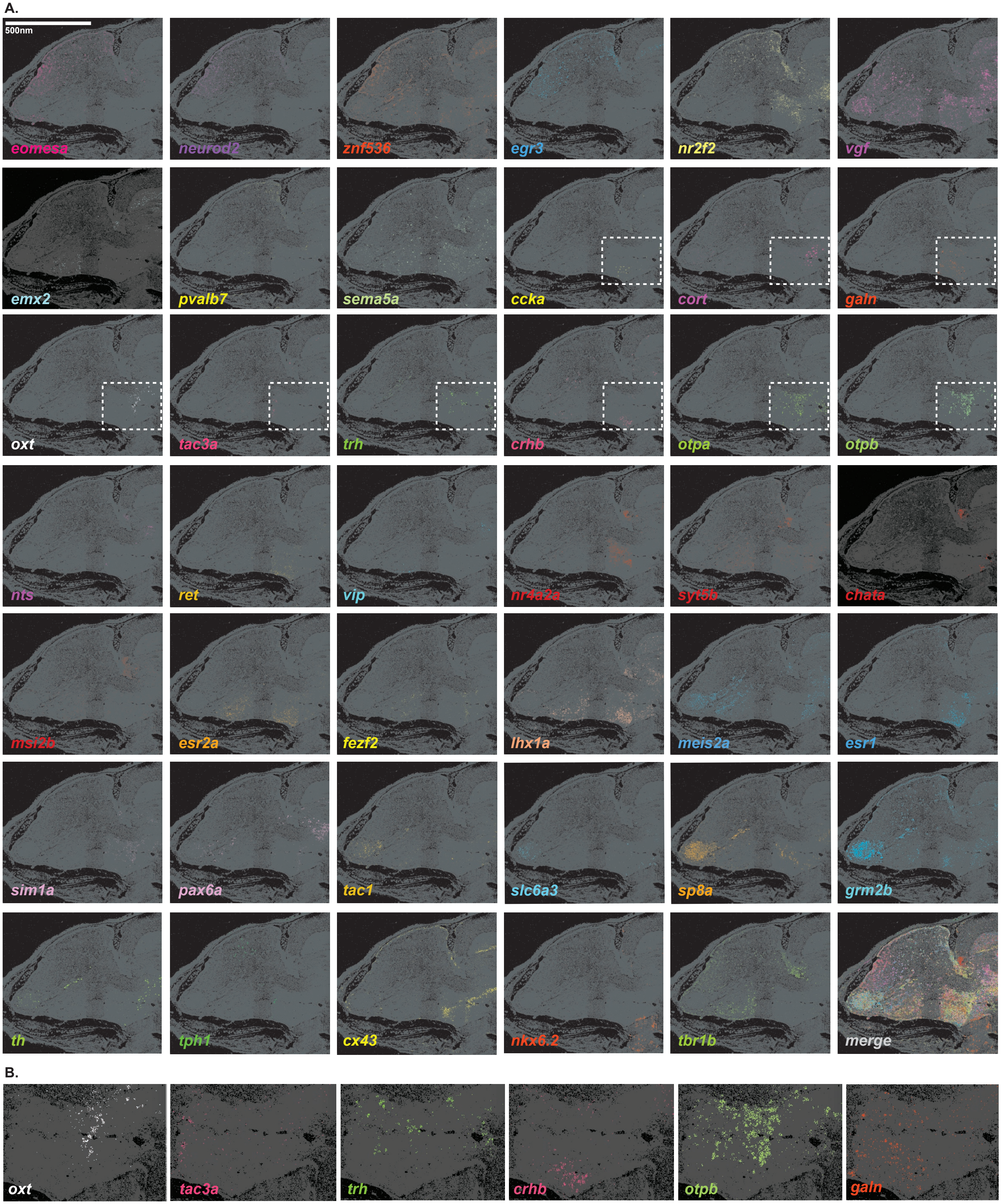
A.
500nm
eomesa
neurod2
znf536
egr3
nr2f2
vgf
emx2
pvalb7
sema5a
ccka
cort
galn
oxt
tac3a
trh
crhb
otpa
otpb
nts
ret
vip
nr4a2a
syt5b
chata
msi2b
esr2a
fezf2
lhx1a
meis2a
esr1
sim1a
pax6a
tac1
slc6a3
sp8a
grm2b
th
tph1
cx43
nkx6.2
tbr1b
merge
B.
oxt
tac3a
trh
crhb
otpb
galn

**fig. S10. MERFISH analysis of genes expressed in the adult *Danionella cerebrum* telencephalon (A)** MERFISH panels showing expression of genes with distinct expression patterns in the telencephalon **(B)** Expanded view of the boxes from the third row on panel A, showing localization of rare neuropeptidergic neurons in the pre-optic area. Scale bar: 500nm.

Figure S11.

A.

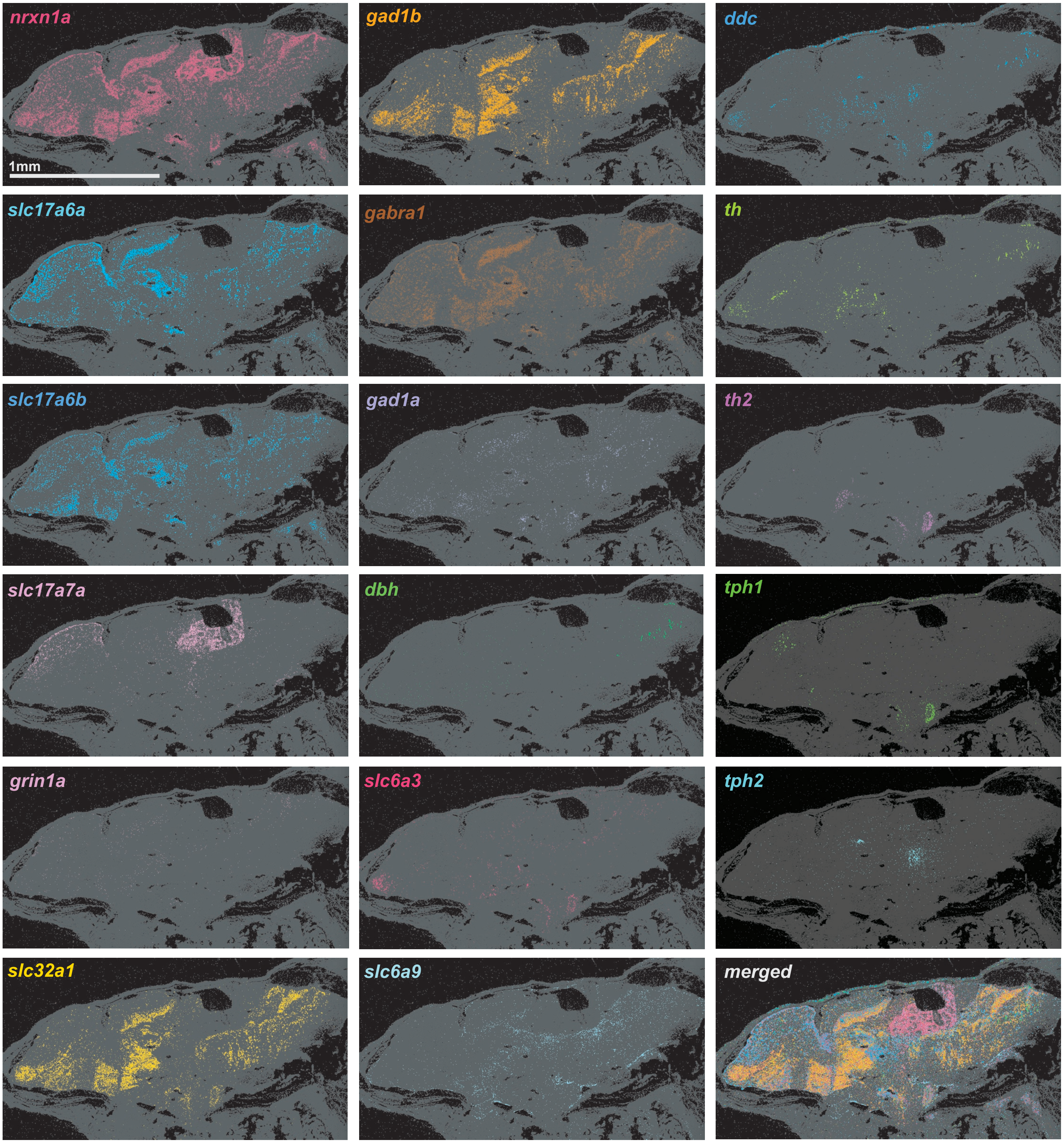

nrxn1a
1mm
gad1b
ddc
slc17a6a
gabra1
th
slc17a6b
gad1a
th2
slc17a7a
dbh
tph1
grin1a
slc6a3
tph2
slc32a1
slc6a9
merged

**fig. S11**. **MERFISH analysis of cell adhesion and neurotransmitter function genes (A)** MERFISH panels showing distribution of GABAergic, glutamatergic and other neurotransmitter class neurons across the brain.

### Figure S12.

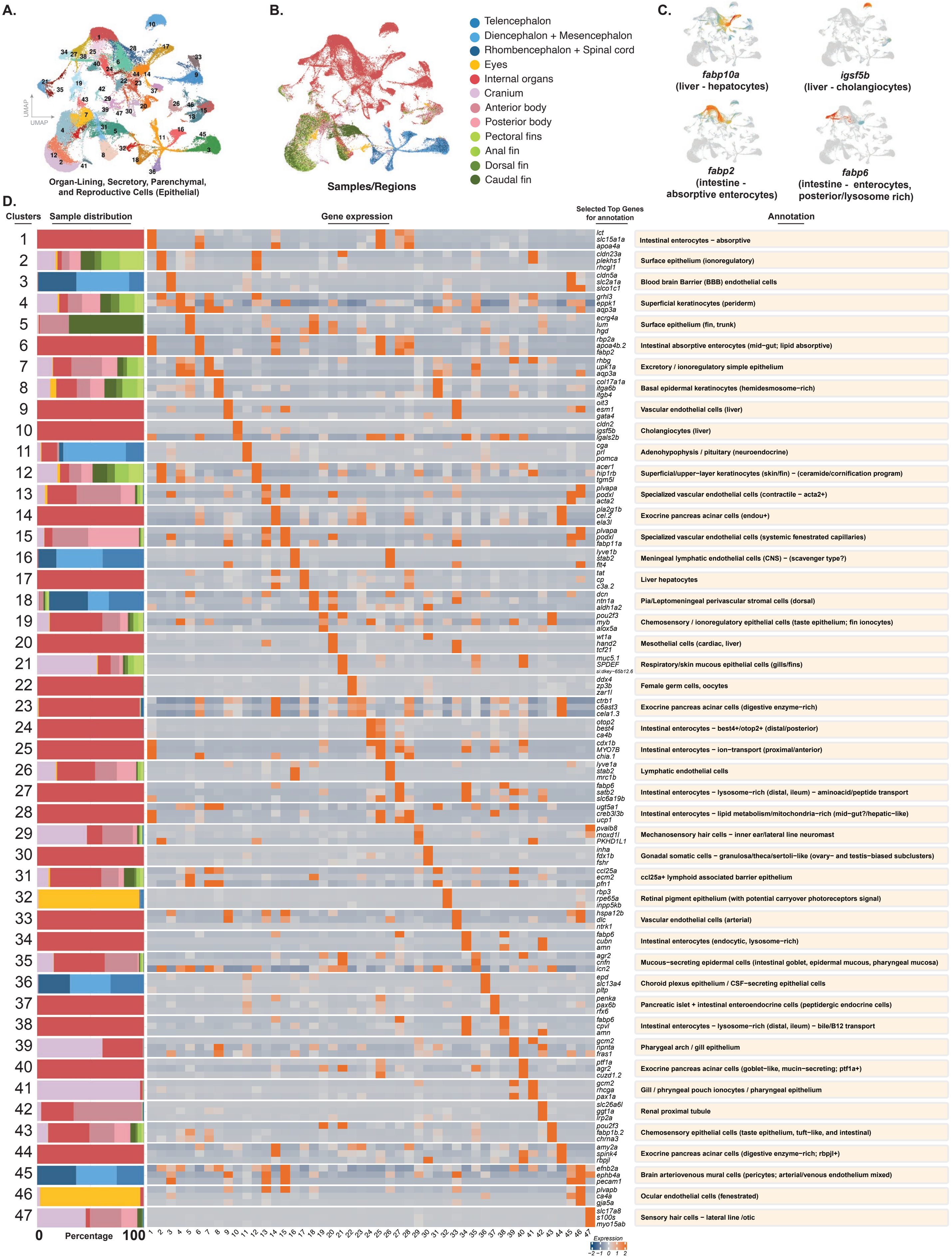

A.
UMAP
Organ-Lining, Secretory, Parenchymal, and Reproductive Cells (Epithelial)
B.
Samples/Regions
Telencephalon
Diencephalon + Mesencephalon
Rhombencephalon + Spinal cord
Eyes
Internal organs
Cranium
Anterior body
Posterior body
Pectoral fins
Anal fin
Dorsal fin
Caudal fin
C.
fabp10a
(liver - hepatocytes)
igsf5b
(liver - cholangiocytes)
fabp2
(intestine - absorptive enterocytes)
fabp6
(intestine - enterocytes, posterior/lysosome rich)
D.
Clusters
Sample distribution
Gene expression
Selected Top Genes for annotation
Annotation
lct slc15a1a apoa4a Intestinal enterocytes – absorptive
cldn23a plekhs1 rhcgl1 Surface epithelium (ionoregulatory)
cldn5a slc2a1a slco1c1 Blood brain Barrier (BBB) endothelial cells
grhl3 eppk1 aqp3a Superficial keratinocytes (periderm)
ecrg4a lum hgd Surface epithelium (fin, trunk)
rbp2a apoa4b.2 fabp2 Intestinal absorptive enterocytes (mid–gut; lipid absorptive) rhbg upk1a aqp3a Excretory / ionoregulatory simple epithelium
col17a1a itga6b itgb4 Basal epidermal keratinocytes (hemidesmosome–rich) oit3 esm1 gata4 Vascular endothelial cells (liver)
cldn2 igsf5b lgals2b Cholangiocytes (liver)
cga prl pomca Adenohypophysis / pituitary (neuroendocrine)
acer1 hip1rb tgm5l Superficial/upper–layer keratinocytes (skin/fin) – (ceramide/cornification program) plvapa podxl acta2 Specialized vascular endothelial cells (contractile – acta2+) pla2g1b cel.2 ela3l Exocrine pancreas acinar cells (endou+)
plvapa podxl fabp11a Specialized vascular endothelial cells (systemic fenestrated capillaries) lyve1b stab2 flt4 Meningeal lymphatic endothelial cells (CNS) – (scavenger type?) tat cp c3a.2 Liver hepatocytes
dcn ntn1a aldh1a2 Pia/Leptomeningeal perivascular stromal cells (dorsal) pou2f3 myb alox5a Chemosensory / ionoregulatory epithelial cells (taste epithelium; fin ionocytes) wt1a hand2 tcf21 Mesothelial cells (cardiac, liver)
muc5.1 SPDEF si:dkey–65b12.6 Respiratory/skin mucous epithelial cells (gills/fins) ddx4 zp3b zar1l Female germ cells, oocytes
ctrb1 c6ast3 cela1.3 Exocrine pancreas acinar cells (digestive enzyme–rich) otop2 best4 ca4b Intestinal enterocytes – best4+/otop2+ (distal/posterior) cdx1b MYO7B chia.1 Intestinal enterocytes – ion–transport (proximal/anterior) lyve1a stab2 mrc1b Lymphatic endothelial cells
fabp6 satb2 slc6a19b Intestinal enterocytes – lysosome–rich (distal, ileum) – aminoacid/peptide transport ugt5a1 creb3l3b ucp1 Intestinal enterocytes – lipid metabolism/mitochondria–rich (mid–gut?/hepatic–like) pvalb8 moxd1l PKHD1L1 Mechanosensory hair cells – inner ear/lateral line neuromast inha fdx1b fshr Gonadal somatic cells – granulosa/theca/sertoli–like (ovary– and testis–biased subclusters) ccl25a ecm2 pfn1 ccl25a+ lymphoid associated barrier epithelium
rbp3 rpe65a inpp5kb Retinal pigment epithelium (with potential carryover photoreceptors signal) hspa12b dlc ntrk1 Vascular endothelial cells (arterial)
fabp6 cubn amn Intestinal enterocytes (endocytic, lysosome–rich)
agr2 cnfn icn2 Mucous–secreting epidermal cells (intestinal goblet, epidermal mucous, pharyngeal mucosa) epd slc13a4 pltp Choroid plexus epithelium / CSF–secreting epithelial cells penka pax6b rfx6 Pancreatic islet + intestinal enteroendocrine cells (peptidergic endocrine cells) fabp6 cpvl amn Intestinal enterocytes – lysosome–rich (distal, ileum) – bile/B12 transport gcm2 npnta fras1 Pharygeal arch / gill epithelium
ptf1a agr2 cuzd1.2 Exocrine pancreas acinar cells (goblet–like, mucin–secreting; ptf1a+) gcm2 rhcga pax1a Gill / phryngeal pouch ionocytes / pharyngeal epithelium slc26a6l ggt1a lrp2a Renal proximal tubule
pou2f3 fabp1b.2 chrna3 Chemosensory epithelial cells (taste epithelium, tuft–like, and intestinal) amy2a spink4 rbpjl Exocrine pancreas acinar cells (digestive enzyme–rich; rbpjl+) efnb2a ephb4a pecam1 Brain arteriovenous mural cells (pericytes; arterial/venous endothelium mixed) plvapb ca4a gja5a Ocular endothelial cells (fenestrated)
slc17a8 s100s myo15ab Sensory hair cells – lateral line /otic
0 Percentage 100
Expression
−2 −1 0 1 2

**fig. S12. Characterization of epithelial (organ-lining, secretory, parenchymal, and reproductive) cell classes of the body (A)** UMAP representation of all organ-lining, secretory, parenchymal, and reproductive, epithelial, clusters. **(B)** UMAP plot showing distribution of experimental samples across the dataset. **(C)** Feature plot examples indicating the expression of key genes marking specific cell types. **(D)** Heatmap showing expression of selected genes, including top-enriched genes per cluster, used to annotate each cluster, and histograms showing distribution of experimental samples for each epithelial cluster.

**Figure S13.**

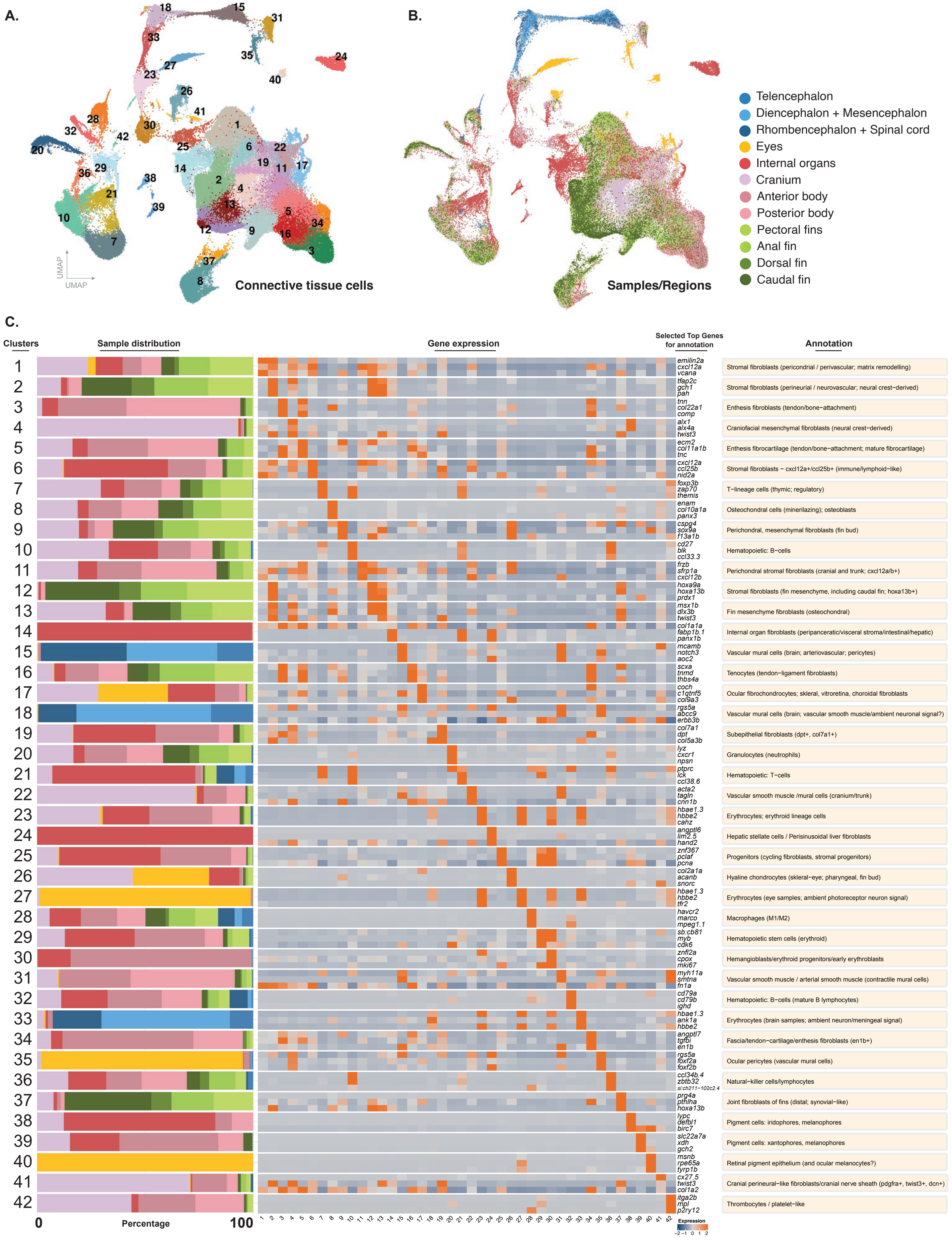
A.
B.
C.
Connective tissue cells
UMAP
UMAP
Samples/Regions
Telencephalon
Diencephalon + Mesencephalon
Rhombencephalon + Spinal cord
Eyes
Internal organs
Cranium
Anterior body
Posterior body
Pectoral fins
Anal fin
Dorsal fin
Caudal fin
Clusters
Sample distribution
Gene expression
Selected Top Genes for annotation
Annotation
emilin2a cxcl12a vcana — Stromal fibroblasts (pericondrial / perivascular; matrix remodelling)
tfap2c gch1 pah — Stromal fibroblasts (perineurial / neurovascular; neural crest−derived)
tnn col22a1 comp — Enthesis fibroblasts (tendon/bone−attachment)
alx1 alx4a twist3 — Craniofacial mesenchymal fibroblasts (neural crest−derived)
ecm2 col11a1b tnc — Enthesis fibrocartilage (tendon/bone−attachment; mature fibrocartilage)
cxcl12a ccl25b nid2a — Stromal fibroblasts − cxcl12a+/ccl25b+ (immune/lymphoid−like)
foxp3b zap70 themis — T−lineage cells (thymic; regulatory)
enam col10a1a panx3 — Osteochondral cells (minerilazing); osteoblasts
cspg4 sox9a f13a1b — Perichondral, mesenchymal fibroblasts (fin bud)
cd27 blk ccl33.3 — Hematopoietic: B−cells
frzb sfrp1a cxcl12b — Perichondral stromal fibroblasts (cranial and trunk; cxcl12a/b+)
hoxa9a hoxa13b prdx1 — Stromal fibroblasts (fin mesenchyme, including caudal fin; hoxa13b+)
msx1b dlx3b twist3 — Fin mesenchyme fibroblasts (osteochondral)
col1a1a fabp1b.1 panx1b — Internal organ fibroblasts (peripanceratic/visceral stroma/intestinal/hepatic)
mcamb notch3 aoc2 — Vascular mural cells (brain; arteriovascular; pericytes)
scxa tnmd thbs4a — Tenocytes (tendon−ligament fibroblasts)
coch c1qtnf5 col9a3 — Ocular fibrochondrocytes; skleral, vitroretina, choroidal fibroblasts
rgs5a abcc9 erbb3b — Vascular mural cells (brain; vascular smooth muscle/ambient neuronal signal?)
col7a1 dpt col5a3b — Subepithelial fibroblasts (dpt+, col7a1+)
lyz cxcr1 npsn — Granulocytes (neutrophils)
ptprc lck ccl38.6 — Hematopoietic: T−cells
acta2 tagln cnn1b — Vascular smooth muscle /mural cells (cranium/trunk)
hbae1.3 hbbe2 cahz — Erythrocytes; erythroid lineage cells
angptl6 lim2.5 hand2 — Hepatic stellate cells / Perisinusoidal liver fibroblasts
znf367 pclaf pcna — Progenitors (cycling fibroblasts, stromal progenitors)
col2a1a acanb snorc — Hyaline chondrocytes (skleral−eye; pharyngeal, fin bud)
hbae1.3 hbbe2 tfr2 — Erythrocytes (eye samples; ambient photoreceptor neuron signal)
havcr2 marco mpeg1.1 — Macrophages (M1/M2)
sb:cb81 myb cdk6 — Hematopoietic stem cells (erythroid)
znfl2a cpox mki67 — Hemangioblasts/erythroid progenitors/early erythroblasts
myh11a smtna fn1a — Vascular smooth muscle / arterial smooth muscle (contractile mural cells)
cd79a cd79b ighd — Hematopoietic: B−cells (mature B lymphocytes)
hbae1.3 ank1a hbbe2 — Erythrocytes (brain samples; ambient neuron/meningeal signal)
angptl7 tgfbi en1b — Fascia/tendon−cartilage/enthesis fibroblasts (en1b+)
rgs5a foxf2a foxf2b — Ocular pericytes (vascular mural cells)
ccl34b.4 zbtb32 si:ch211−102c2.4 — Natural−killer cells/lymphocytes
prg4a pthlha hoxa13b — Joint fibroblasts of fins (distal; synovial−like)
lypc defbl1 birc7 — Pigment cells: iridophores, melanophores
slc22a7a xdh gch2 — Pigment cells: xantophores, melanophores
msnb rpe65a tyrp1b — Retinal pigment epithelium (and ocular melanocytes?)
cx27.5 twist3 col1a2 — Cranial perineural−like fibroblasts/cranial nerve sheath (pdgfra+, twist3+, dcn+)
itga2b mpl p2ry12 — Thrombocytes / platelet−like
0
Percentage
100
Expression
−2 −1 0 1 2

**fig. S13. Characterization of connective cell classes of the body (A)** UMAP representation of all connective cell clusters. **(B)** UMAP plot showing distribution of experimental samples across the dataset. **(C)** Heatmap showing expression of selected genes, including top-enriched genes per cluster, used to annotate each cluster, and histograms showing distribution of experimental samples for each connective cell cluster.

Figure S14.

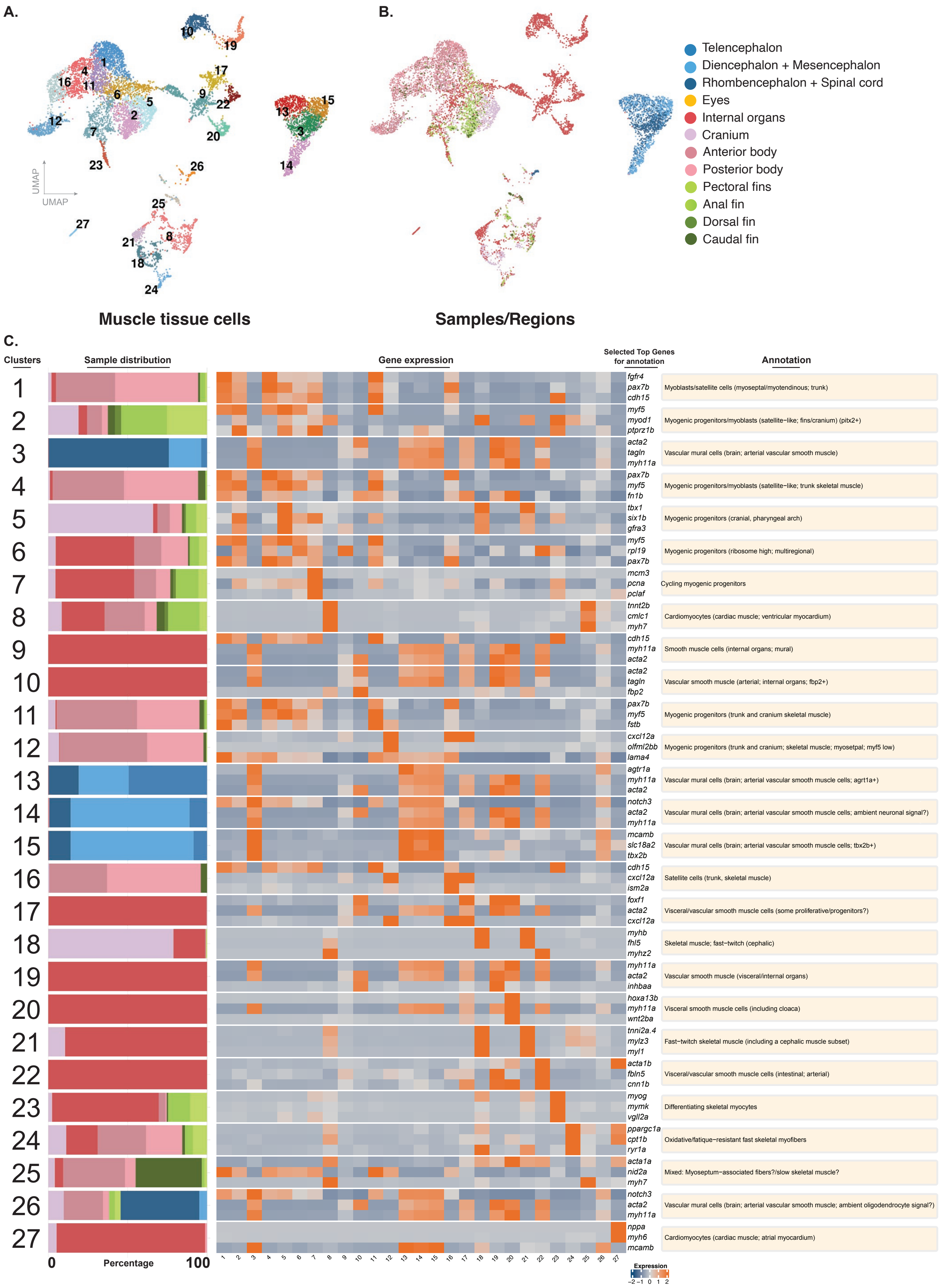
A.
B.
C.
UMAP
UMAP
Telencephalon
Diencephalon + Mesencephalon
Rhombencephalon + Spinal cord
Eyes
Internal organs
Cranium
Anterior body
Posterior body
Pectoral fins
Anal fin
Dorsal fin
Caudal fin
Muscle tissue cells
Samples/Regions
Clusters
Sample distribution
Gene expression
Selected Top Genes for annotation
Annotation
fgfr4 pax7b cdh15 Myoblasts/satellite cells (myoseptal/myotendinous; trunk) myf5 myod1 ptprz1b Myogenic progenitors/myoblasts (satellite−like; fins/cranium) (pitx2+) acta2 tagln myh11a Vascular mural cells (brain; arterial vascular smooth muscle) pax7b myf5 fn1b Myogenic progenitors/myoblasts (satellite−like; trunk skeletal muscle) tbx1 six1b gfra3 Myogenic progenitors (cranial, pharyngeal arch)
myf5 rpl19 pax7b Myogenic progenitors (ribosome high; multiregional)
mcm3 pcna pclaf Cycling myogenic progenitors
tnnt2b cmlc1 myh7 Cardiomyocytes (cardiac muscle; ventricular myocardium) cdh15 myh11a acta2 Smooth muscle cells (internal organs; mural)
acta2 tagln fbp2 Vascular smooth muscle (arterial; internal organs; fbp2+) pax7b myf5 fstb Myogenic progenitors (trunk and cranium skeletal muscle) cxcl12a olfml2bb lama4 Myogenic progenitors (trunk and cranium; skeletal muscle; myosetpal; myf5 low) agtr1a myh11a acta2 Vascular mural cells (brain; arterial vascular smooth muscle cells; agrt1a+) notch3 acta2 myh11a Vascular mural cells (brain; arterial vascular smooth muscle cells; ambient neuronal signal?) mcamb slc18a2 tbx2b Vascular mural cells (brain; arterial vascular smooth muscle cells; tbx2b+) cdh15 cxcl12a ism2a Satellite cells (trunk, skeletal muscle)
foxf1 acta2 cxcl12a Visceral/vascular smooth muscle cells (some proliferative/progenitors?) myhb fhl5 myhz2 Skeletal muscle; fast−twitch (cephalic)
myh11a acta2 inhbaa Vascular smooth muscle (visceral/internal organs)
hoxa13b myh11a wnt2ba Visceral smooth muscle cells (including cloaca)
tnni2a.4 mylz3 myl1 Fast−twitch skeletal muscle (including a cephalic muscle subset) acta1b fbln5 cnn1b Visceral/vascular smooth muscle cells (intestinal; arterial) myog mymk vgll2a Differentiating skeletal myocytes
ppargc1a cpt1b ryr1a Oxidative/fatique−resistant fast skeletal myofibers acta1a nid2a myh7 Mixed: Myoseptum−associated fibers?/slow skeletal muscle? notch3 acta2 myh11a Vascular mural cells (brain; arterial vascular smooth muscle; ambient oligodendrocyte signal?) nppa myh6 mcamb Cardiomyocytes (cardiac muscle; atrial myocardium)
0
Percentage
100
Expression
−2 −1 0 1 2

**fig. S14. Characterization of muscle cell classes of the body (A)** UMAP representation of all muscle cell clusters. **(B)** UMAP plot showing distribution of experimental samples across the dataset. **(C)** Heatmap showing expression of selected genes, including top-enriched genes per cluster, used to annotate each cluster, and histograms showing distribution of experimental samples for each muscle cell cluster.

Figure S15.

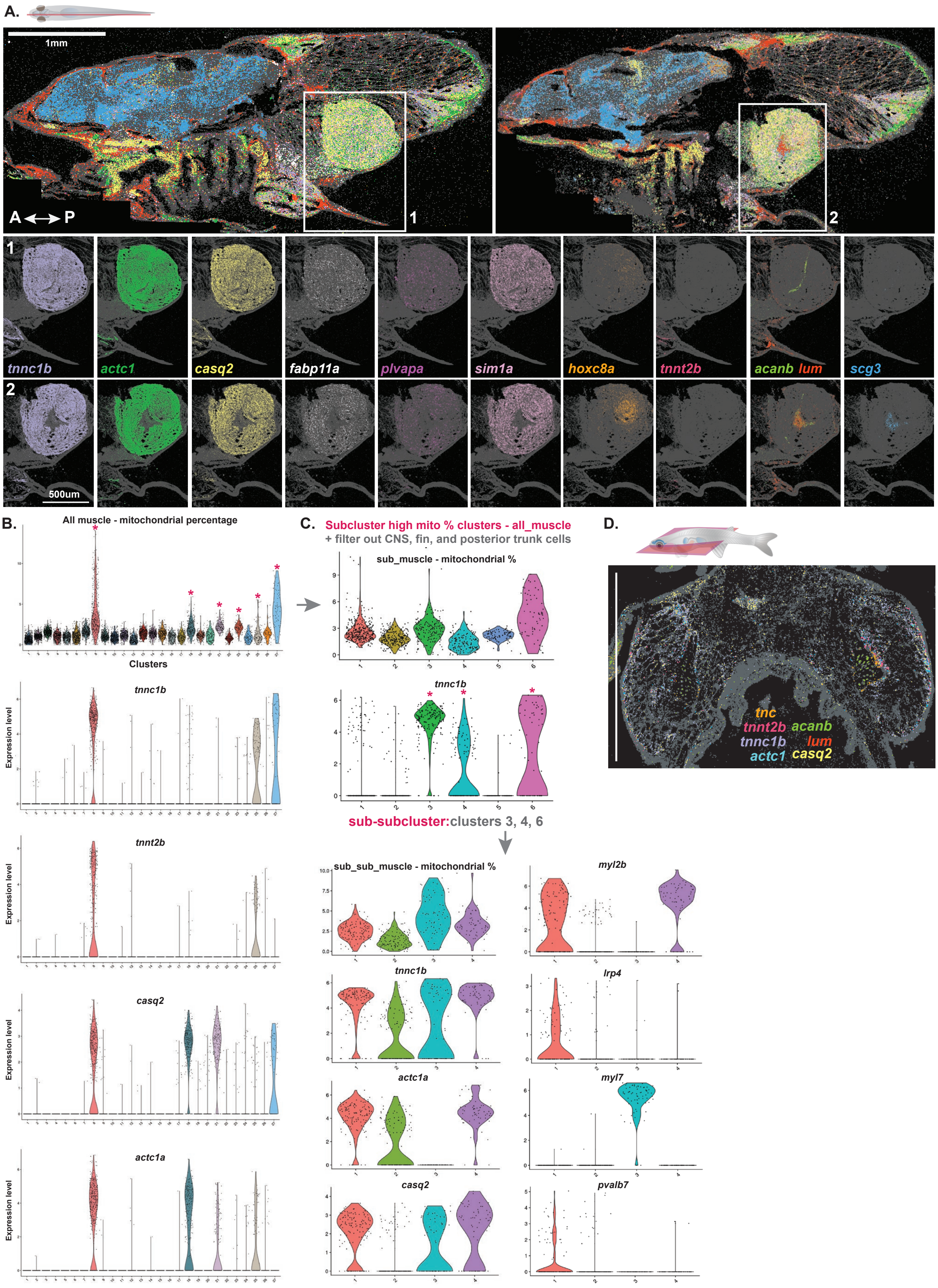
A.
1mm
A ↔ P
1
2
tnnc1b
actc1
casq2
fabp11a
plvapa
sim1a
hoxc8a
tnnt2b
acanb lum
scg3
500um
B.
All muscle - mitochondrial percentage
Clusters
tnnc1b
tnnt2b
casq2
actc1a
Expression level
C.
Subcluster high mito % clusters - all_muscle
+ filter out CNS, fin, and posterior trunk cells
sub_muscle - mitochondrial %
tnnc1b
sub-subcluster:clusters 3, 4, 6
sub_sub_muscle - mitochondrial %
tnnc1b
actc1a
casq2
myl2b
lrp4
myl7
pvalb7
D.
tnc
tnnt2b
acanb
tnnc1b
lum
actc1
casq2

**fig. S15. In-depth characterization of drumming muscle cells (A)** MERFISH results showing expression patterns of a select set of genes in the drumming muscle for two separate mediolateral positions. **(B)** Violin plots indicating percentage of mitochondrial gene expression across muscle clusters, along with the expression of a select subset of genes observed to be expressed in the drumming muscle. **(C)** Sub-clustering mitochondria-rich clusters from the global muscle dataset, and later sub-sub-clustering *tnnc1b*+ clusters from the resulting dataset yield potential drumming muscle clusters. Clusters 1 and 4 in the final dataset show expression of *tnnc1b*, *actc1a*, *casq2*, and *myl2b.* Cluster 1 showed selective expression of *lrp4*, a neuromuscular junction marker, and *pvalb7*, a calcium-binding protein.

**Figure S16.**

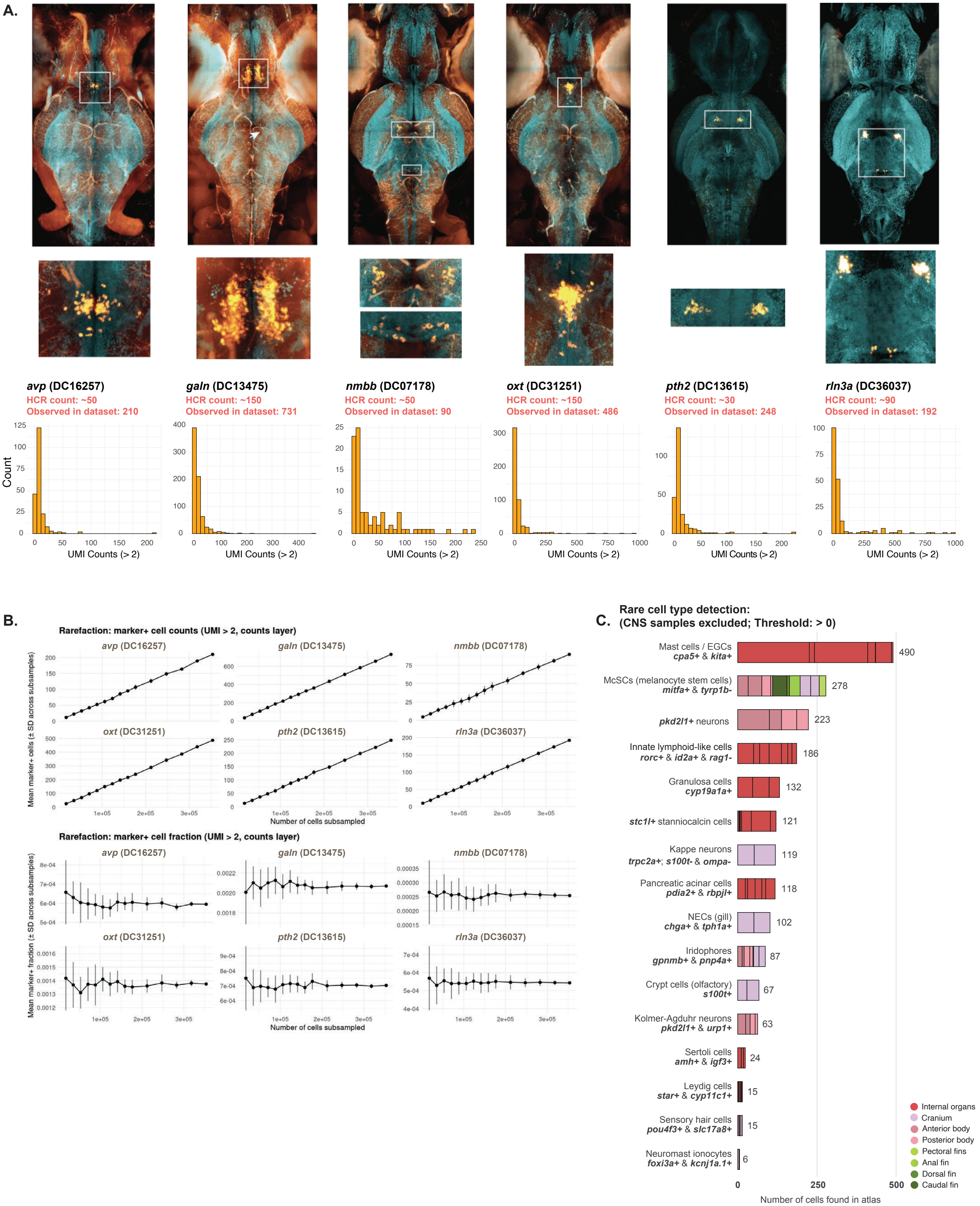
A.
avp (DC16257)
HCR count: ~50
Observed in dataset: 210
galn (DC13475)
HCR count: ~150
Observed in dataset: 731
nmbb (DC07178)
HCR count: ~50
Observed in dataset: 90
oxt (DC31251)
HCR count: ~150
Observed in dataset: 486
pth2 (DC13615)
HCR count: ~30
Observed in dataset: 248
rln3a (DC36037)
HCR count: ~90
Observed in dataset: 192
Count
UMI Counts (> 2)
B.
Rarefaction: marker+ cell counts (UMI > 2, counts layer)
Mean marker+ cells (± SD across subsamples)
Number of cells subsampled
Rarefaction: marker+ cell fraction (UMI > 2, counts layer)
Mean marker+ fraction (± SD across subsamples)
C.
Rare cell type detection:
(CNS samples excluded; Threshold: > 0)
Mast cells / EGCs
cpa5+ & kita+
490
McSCs (melanocyte stem cells)
mitfa+ & tyrp1b-
278
pkd2l1+ neurons
223
Innate lymphoid-like cells
rorc+ & id2a+ & rag1-
186
Granulosa cells
cyp19a1a+
132
stc1l+ stanniocalcin cells
121
Kappe neurons
trpc2a+; s100t- & ompa-
119
Pancreatic acinar cells
pdia2+ & rbpjl+
118
NECs (gill)
chga+ & tph1a+
102
Iridophores
gpnmb+ & pnp4a+
87
Crypt cells (olfactory)
s100t+
67
Kolmer-Agduhr neurons
pkd2l1+ & urp1+
63
Sertoli cells
amh+ & igf3+
24
Leydig cells
star+ & cyp11c1+
15
Sensory hair cells
pou4f3+ & slc17a8+
15
Neuromast ionocytes
foxi3a+ & kcnj1a.1+
6
Number of cells found in atlas
Internal organs
Cranium
Anterior body
Posterior body
Pectoral fins
Anal fin
Dorsal fin
Caudal fin

**fig. S16. Saturation analyses using rare cells (A)** HCR-FISH of rare neuropeptidergic neuron markers and cell counts, with histograms showing the total number of neuropeptidergic cells detected in the whole-body single-cell transcriptome atlas: *avp*+ neuropeptidergic neurons, HCR-FISH baseline count: ~50 cells, observed in atlas: 210 cells; *galn*+ neuropeptidergic neurons, HCR-FISH baseline count: ~150 cells, observed in atlas: 731 cells; *nmbb*+ neuropeptidergic neurons, HCR-FISH baseline count: ~50 cells, observed in atlas: 90 cells; *oxt*+ neuropeptidergic neurons, HCR-FISH baseline count: ~150 cells, observed in atlas: 486 cells; *pth2*+ neuropeptidergic neurons, HCR-FISH baseline count: ~30 cells, observed in atlas: 248 cells; *rln3a*+ neuropeptidergic neurons, HCR-FISH baseline count: ~90 cells, observed in atlas: 192 cells **(B)** Subsampling (rarefaction) analysis showing how reliably a specific marker is detected as the dataset size increases (top), and through random subsampling at multiple increasing fractions (bottom), indicating that saturation for these rare markers was reached. **(C)** Detection and counts of previously noted, known, rare cell types across different tissues in the *Danionella* whole-body single-cell transcriptome atlas.

Figure S17.

A.

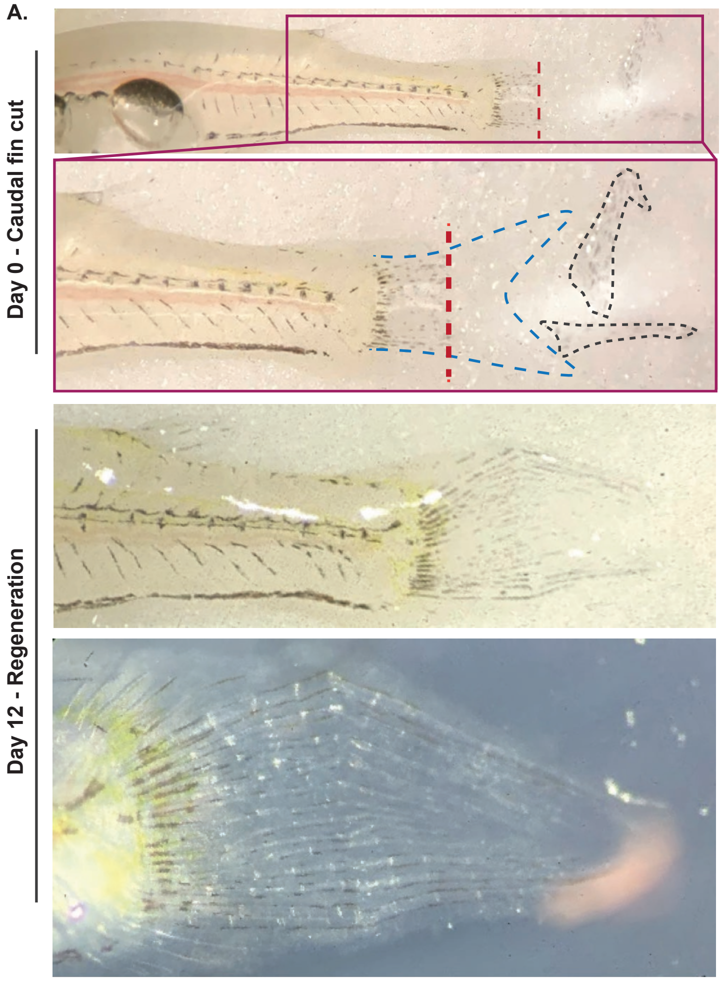

**fig. S17. Caudal fin regeneration in adult *Danionella cerebrum* (A)** Caudal fin regeneration in adult *Danionella cerebrum*, represented by Day 0 caudal fin tip cut and Day 12 regeneration timepoints. Black dotted outlines: cut caudal fin pieces.

Figure S18.

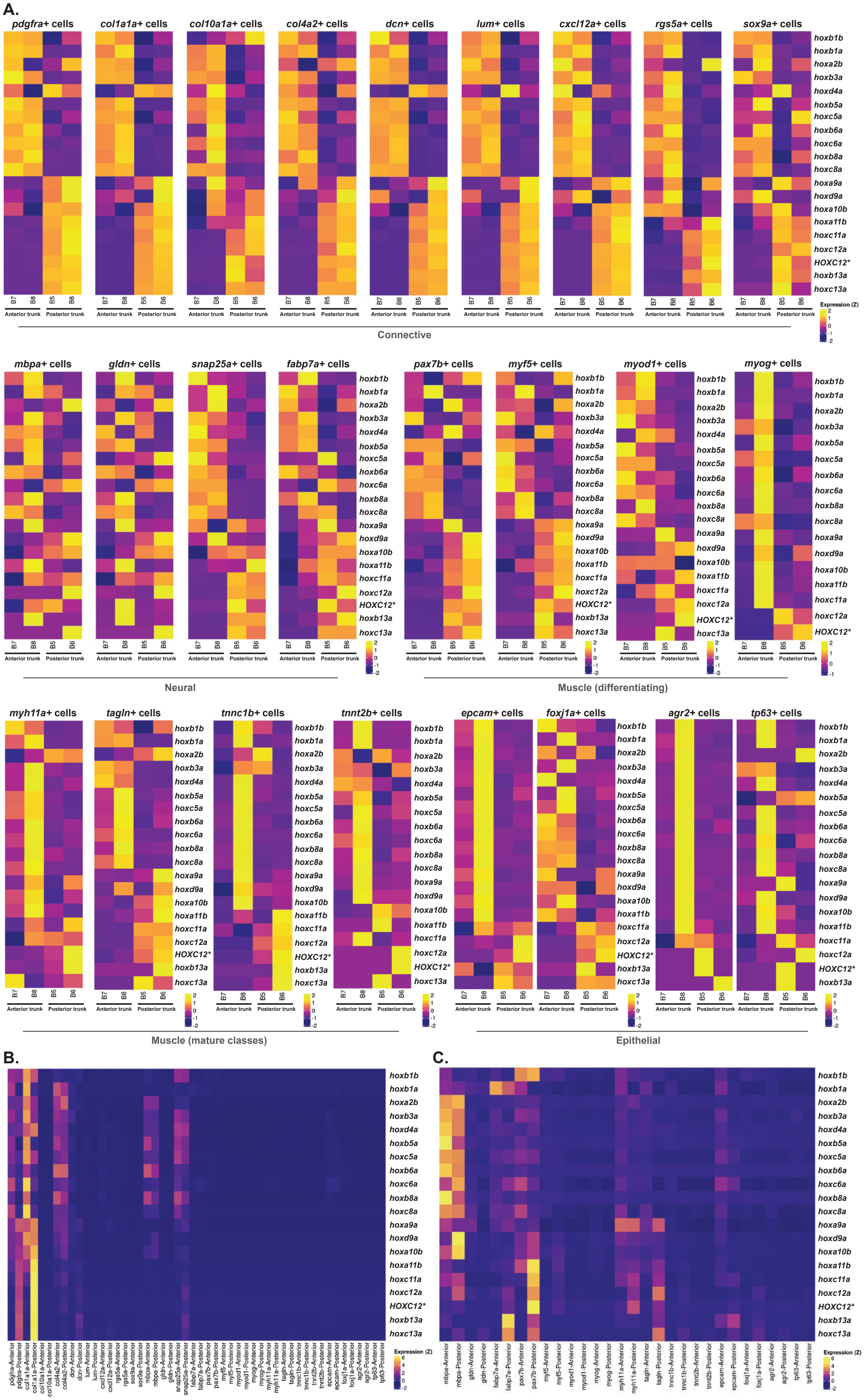
A.
pdgfra+ cells
col1a1a+ cells
col10a1a+ cells
col4a2+ cells
dcn+ cells
lum+ cells
cxcl12a+ cells
rgs5a+ cells
sox9a+ cells
hoxb1b
hoxb1a
hoxa2b
hoxb3a
hoxd4a
hoxb5a
hoxc5a
hoxb6a
hoxc6a
hoxb8a
hoxc8a
hoxa9a
hoxd9a
hoxa10b
hoxa11b
hoxc11a
hoxc12a
HOXC12*
hoxb13a
hoxc13a
B7
B8
B5
B6
Anterior trunk
Posterior trunk
Expression (Z)
Connective
mbpa+ cells
gldn+ cells
snap25a+ cells
fabp7a+ cells
pax7b+ cells
myf5+ cells
myod1+ cells
myog+ cells
Neural
Muscle (differentiating)
myh11a+ cells
tagln+ cells
tnnc1b+ cells
tnnt2b+ cells
epcam+ cells
foxj1a+ cells
agr2+ cells
tp63+ cells
Muscle (mature classes)
Epithelial
B.
C.

**fig. S18. Comparative differential expression analysis of positional information across adult *Danionella cerebrum* tissue and cell classes (A)** Comparisons of expression patterns of Hox genes in anterior and posterior trunk cell types. **(B)** Heatmap indicating higher expression levels of Hox genes in connective tissue cell types relative to neural, muscle, and epithelial cell types found in anterior and posterior trunk samples. **(C)** Heatmap indicating higher expression levels of Hox genes in connective and neural cell types found in anterior and posterior trunk samples when connective tissue cell types were removed from the analysis.

Figure S19.

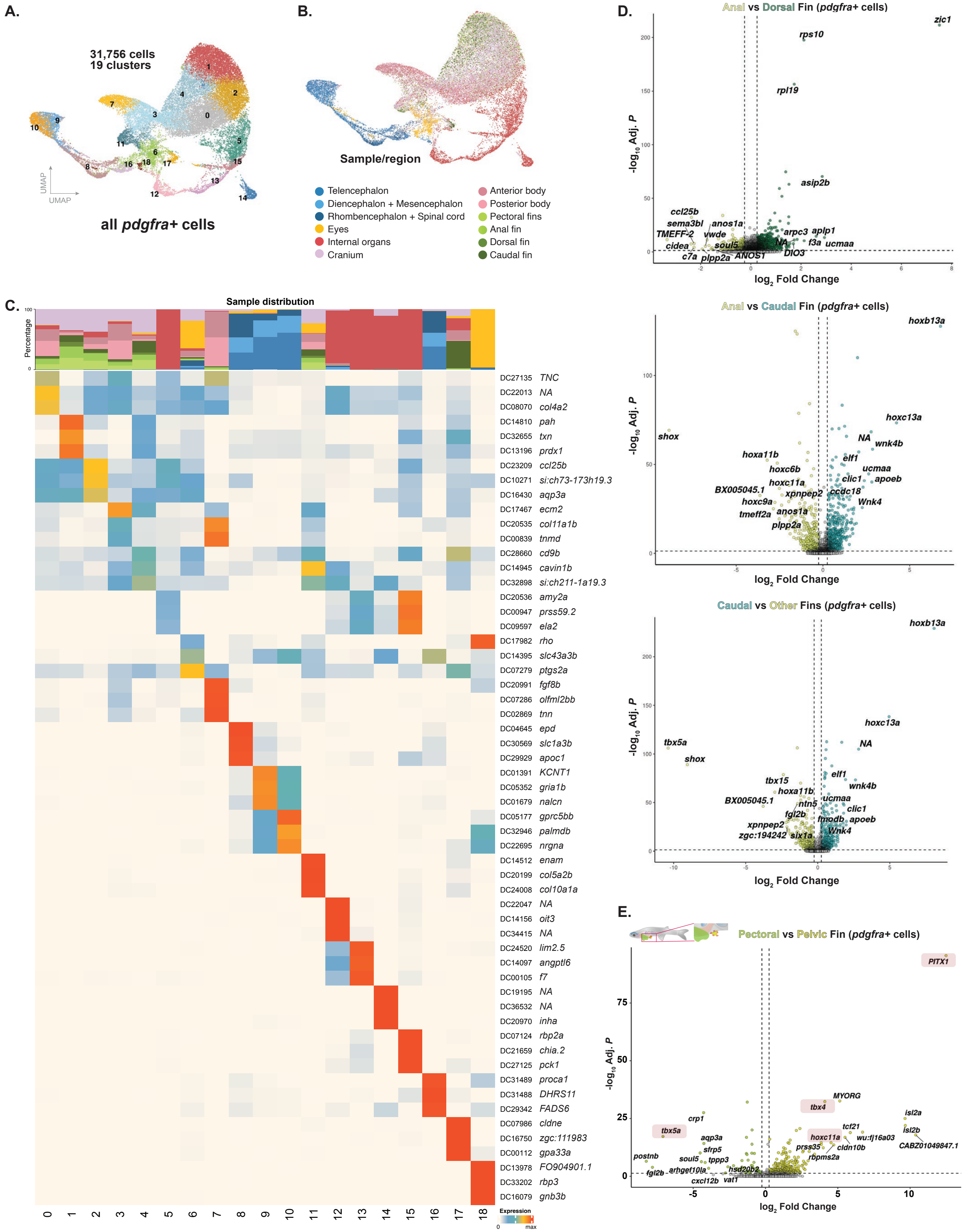
A.
31,756 cells
19 clusters
UMAP
UMAP
all pdgfra+ cells
B.
Sample/region
Telencephalon
Diencephalon + Mesencephalon
Rhombencephalon + Spinal cord
Eyes
Internal organs
Cranium
Anterior body
Posterior body
Pectoral fins
Anal fin
Dorsal fin
Caudal fin
C.
Sample distribution
Percentage
Expression
D.
Anal vs Dorsal Fin (pdgfra+ cells)
Anal vs Caudal Fin (pdgfra+ cells)
Caudal vs Other Fins (pdgfra+ cells)
-log10 Adj. P
log2 Fold Change
E.
Pectoral vs Pelvic Fin (pdgfra+ cells)

**fig. S19. Analysis of positional information in fibroblasts across different tissues (A)** UMAP representation of all *pdgfra*+ fibroblasts clusters in the adult body. **(B)** UMAP plot showing distribution of experimental samples across the fibroblasts dataset, indicating the diversity of fibroblasts across different tissue classes. **(C)** Top three markers and sample distributions for fibroblast clusters are shown. **(D)** Volcano plots showing the top genes with enriched expression in fibroblasts when anal-dorsal, anal-caudal, and caudal-all other fin fibroblasts were compared. **(E)** Volcano plots demonstrating the top differentially expressed genes between pdgfra+ pectoral fibroblasts and *pdgfra*+/*tbx4*+ putative pelvic fin fibroblasts.

Figure S20.

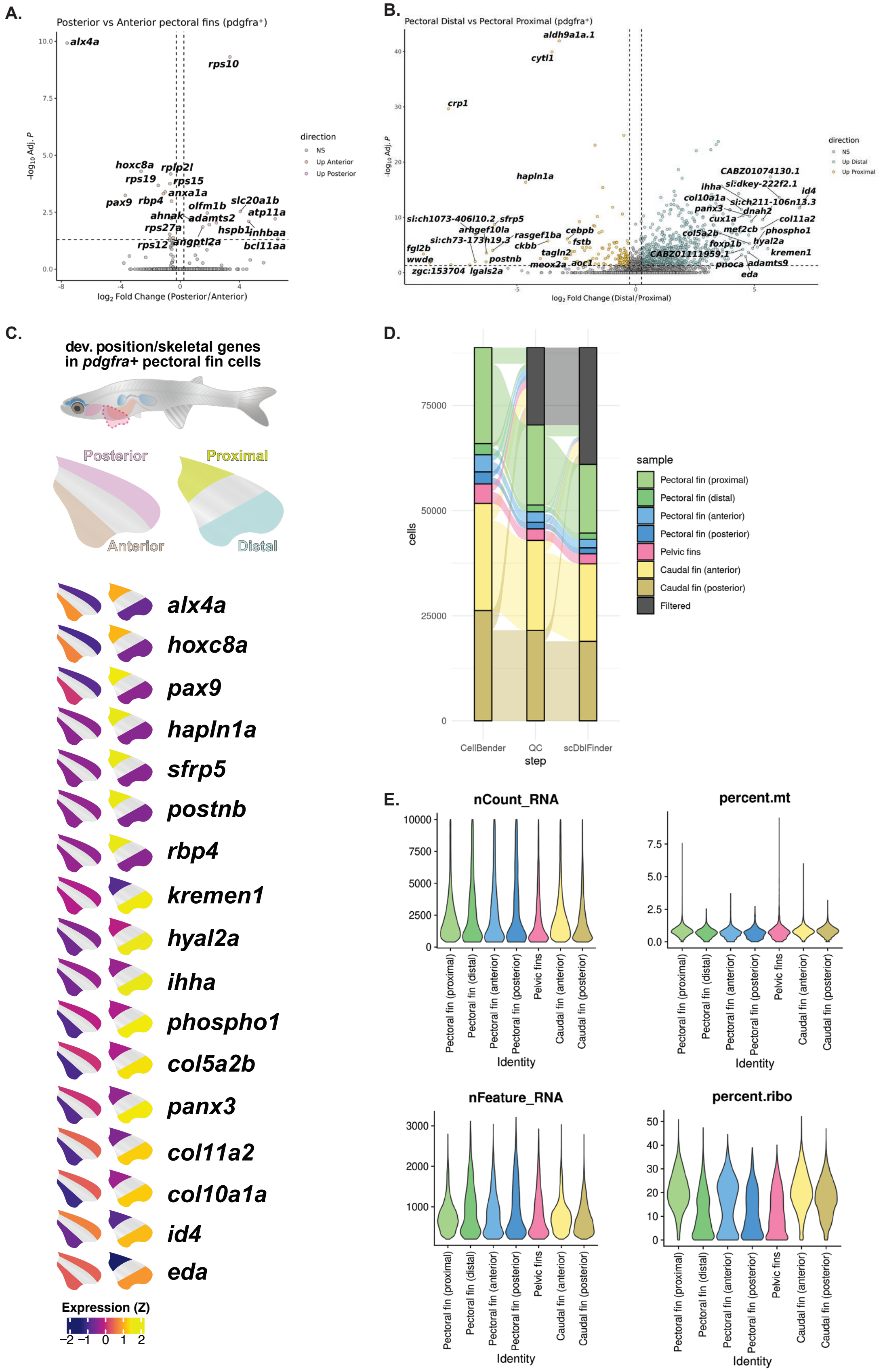

A.
Posterior vs Anterior pectoral fins (pdgfra⁺)
direction
NS
Up Anterior
Up Posterior
log2 Fold Change (Posterior/Anterior)
B.
Pectoral Distal vs Pectoral Proximal (pdgfra⁺)
direction
NS
Up Distal
Up Proximal
log2 Fold Change (Distal/Proximal)
C.
dev. position/skeletal genes in pdgfra+ pectoral fin cells
Posterior
Proximal
Anterior
Distal
alx4a
hoxc8a
pax9
hapln1a
sfrp5
postnb
rbp4
kremen1
hyal2a
ihha
phospho1
col5a2b
panx3
col11a2
col10a1a
id4
eda
Expression (Z)
D.
sample
Pectoral fin (proximal)
Pectoral fin (distal)
Pectoral fin (anterior)
Pectoral fin (posterior)
Pelvic fins
Caudal fin (anterior)
Caudal fin (posterior)
Filtered
cells
CellBender
QC
scDblFinder
step
E.
nCount_RNA
percent.mt
nFeature_RNA
percent.ribo
Identity

**fig. S20. Differential gene expression across anteroposterior and proximodistal pectoral fin axes (A)** Volcano plot showing top differentially expressed genes across anteroposterior axis of the pectoral fin. **(B)** Volcano plot showing top differentially expressed genes across proximodistal axis of the pectoral fin. **(C)** Expanded summary heatmap of top differentially expressed genes across different pectoral fin axes. Summary version of this heatmap is presented in Fig. 3L. **(D)** Sankey plot showing computational quality control steps taken, the list of experimental fin samples: pectoral fin positions, caudal fin positions and the pelvic fins **(E)** Violin plots showing the percentage of the number of uniquely expressed genes (nFeatures) and the percentage of the number of RNA molecules (UMIs) detected across all genes (nCounts) and violin plots indicating the percentage of mitochondrial and ribosomal genes detected across these experimental fin samples.

### Figure S21.

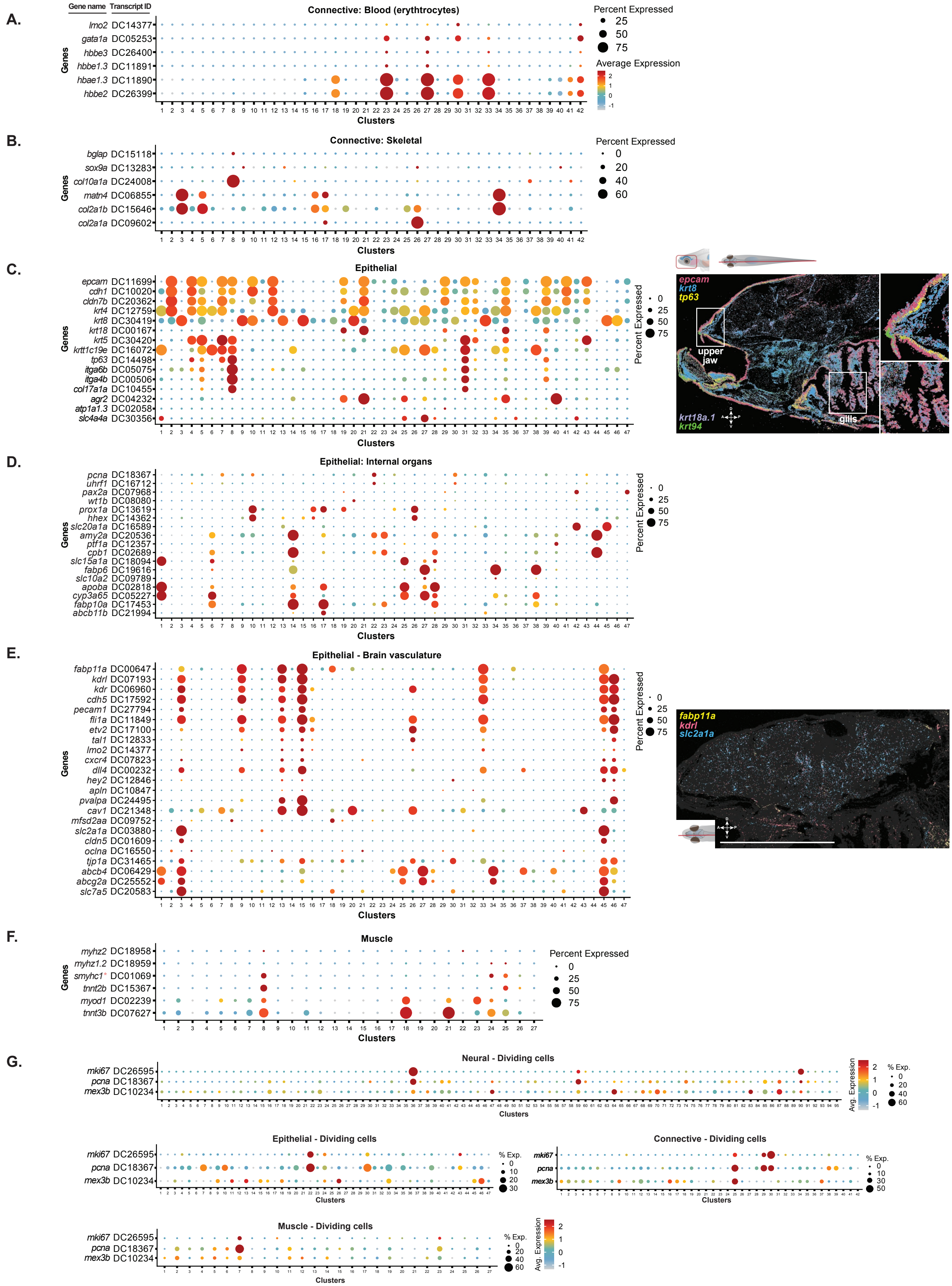

A.
Gene name
Transcript ID
Connective: Blood (erythrocytes)
lmo2 DC14377
gata1a DC05253
hbbe3 DC26400
hbbe1.3 DC11891
hbae1.3 DC11890
hbbe2 DC26399
Genes
Clusters
Percent Expressed
Average Expression
B.
Connective: Skeletal
bglap DC15118
sox9a DC13283
col10a1a DC24008
matn4 DC06855
col2a1b DC15646
col2a1a DC09602
C.
Epithelial
epcam DC11699
cdh1 DC10020
cldn7b DC20362
krt4 DC12759
krt8 DC30419
krt18 DC00167
krt5 DC30420
krtt1c19e DC16072
tp63 DC14498
itga6b DC05075
itga4b DC00506
col17a1a DC10455
agr2 DC04232
atp1a1.3 DC02058
slc4a4a DC30356
epcam
krt8
tp63
upper jaw
gills
krt18a.1
krt94
D.
Epithelial: Internal organs
pcna DC18367
uhrf1 DC16712
pax2a DC07968
wt1b DC08080
prox1a DC13619
hhex DC14362
slc20a1a DC16589
amy2a DC20536
ptf1a DC12357
cpb1 DC02689
slc15a1a DC18094
fabp6 DC19616
slc10a2 DC09789
apoba DC02818
cyp3a65 DC05227
fabp10a DC17453
abcb11b DC21994
E.
Epithelial - Brain vasculature
fabp11a DC00647
kdrl DC07193
kdr DC06960
cdh5 DC17592
pecam1 DC27794
fli1a DC11849
etv2 DC17100
tal1 DC12833
lmo2 DC14377
cxcr4 DC07823
dll4 DC00232
hey2 DC12846
apln DC10847
pvalpa DC24495
cav1 DC21348
mfsd2aa DC09752
slc2a1a DC03880
cldn5 DC01609
oclna DC16550
tjp1a DC31465
abcb4 DC06429
abcg2a DC25552
slc7a5 DC20583
fabp11a
kdrl
slc2a1a
F.
Muscle
myhz2 DC18958
myhz1.2 DC18959
smyhc1 DC01069
tnnt2b DC15367
myod1 DC02239
tnnt3b DC07627
G.
Neural - Dividing cells
mki67 DC26595
pcna DC18367
mex3b DC10234
Avg. Expression
% Exp.
Epithelial - Dividing cells
Connective - Dividing cells
Muscle - Dividing cells

**fig. S21. Dot plots of curated gene sets to assess potential paedomorphic features (A-F)** Dot plots showing expression of a select set of genes marking developmental and mature cell states across the clusters of connective, epithelial, and muscle datasets, respectively. (G) Dot plots showing expression of cell division markers (*mki67*, *pcna*, and *mex3b*) in neural, epithelial, connective and muscle datasets.

**Figure S22.**

**A.**

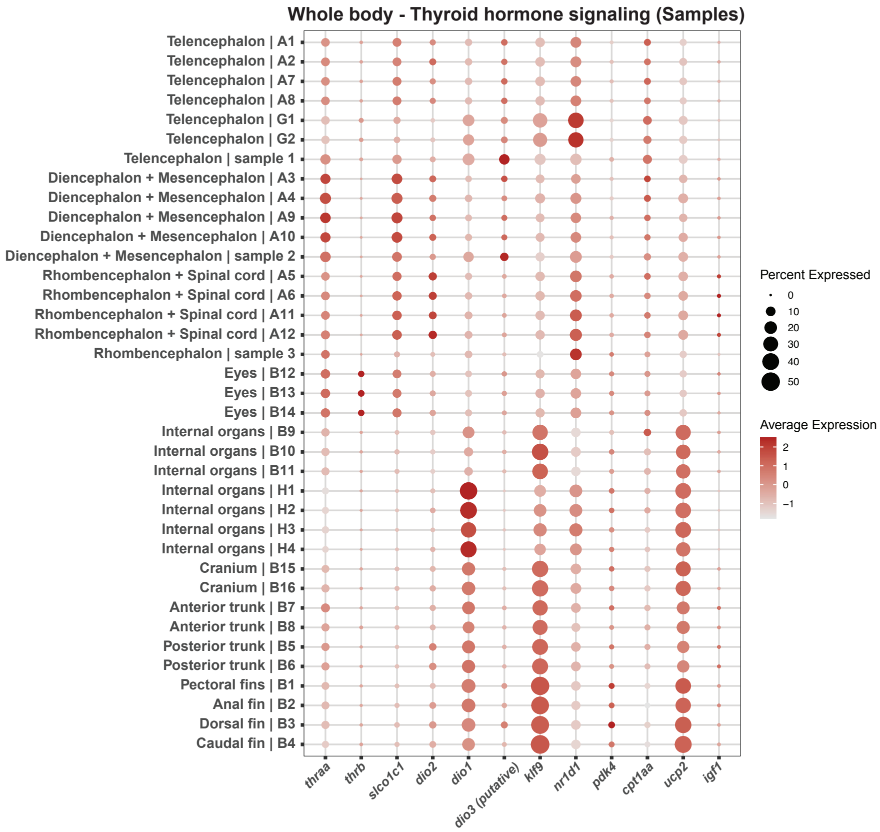

**B.**

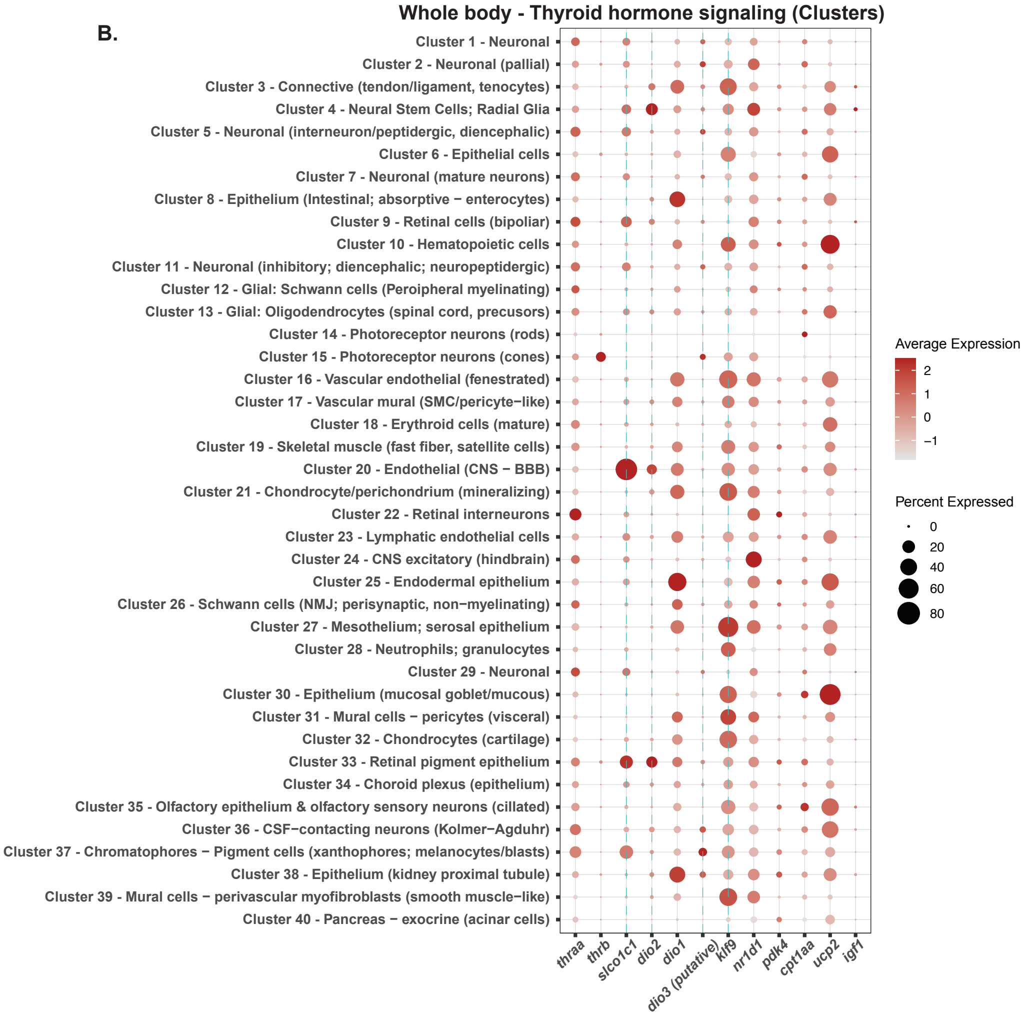

**fig. S22. Thyroid hormone signaling across adult tissues (A)** Expression of thyroid signaling-related genes, including thyroid receptors, delivery-, activation-, and direct output-related genes and metabolic effector genes across all experimental samples, and **(B)** clusters of the whole-body single-cell transcriptome atlas of the adult *Danionella cerebrum*.

Figure S23.

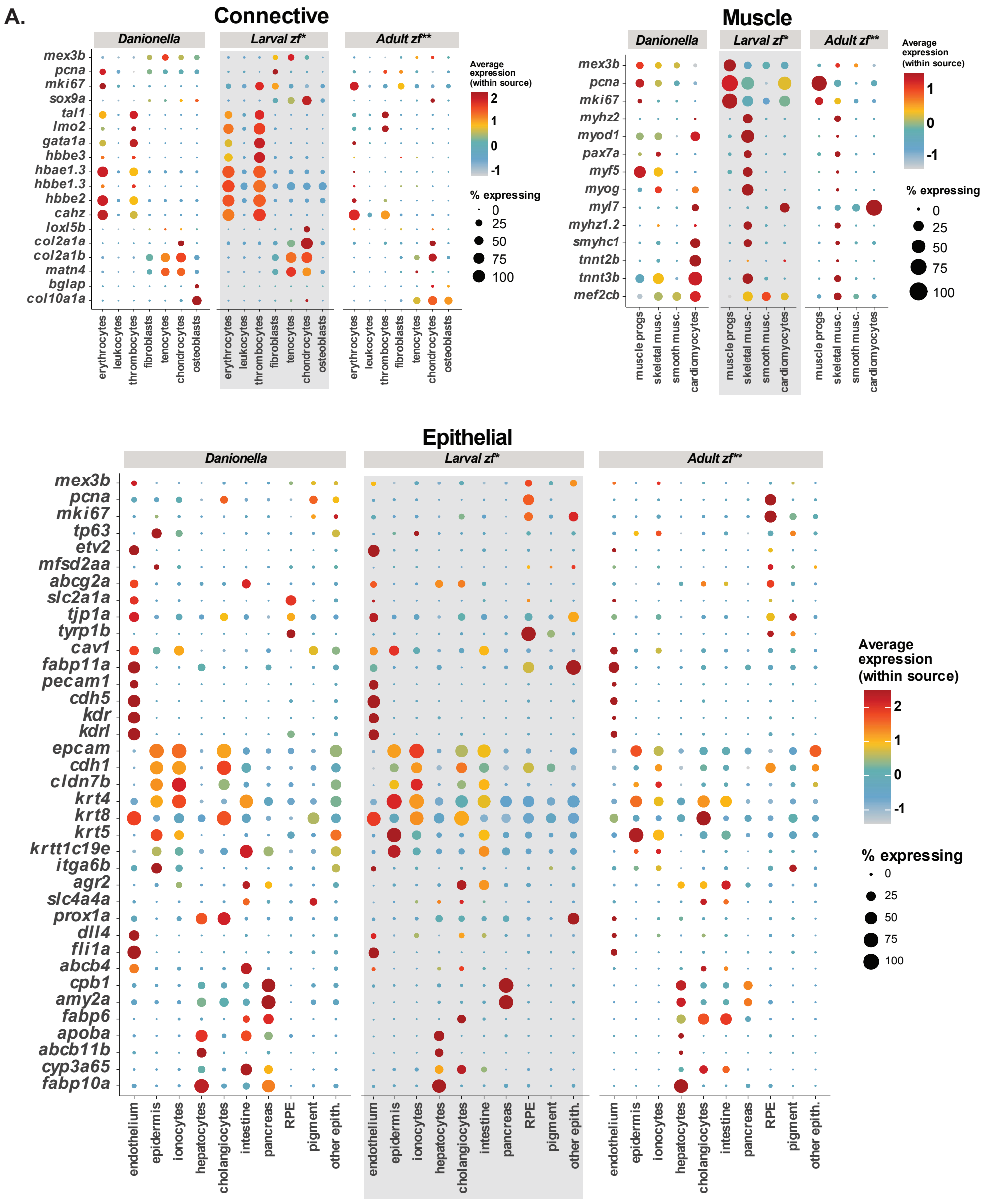
A.
Connective
Danionella
Larval zf*
Adult zf**
Average expression (within source)
% expressing
0
25
50
75
100
mex3b
pcna
mki67
sox9a
tal1
lmo2
gata1a
hbbe3
hbae1.3
hbbe1.3
hbbe2
cahz
loxl5b
col2a1a
col2a1b
matn4
bglap
col10a1a
erythrocytes
leukocytes
thrombocytes
fibroblasts
tenocytes
chondrocytes
osteoblasts
Muscle
myhz2
myod1
pax7a
myf5
myog
myl7
myhz1.2
smyhc1
tnnt2b
tnnt3b
mef2cb
muscle progs
skeletal musc.
smooth musc.
cardiomyocytes
Epithelial
tp63
etv2
mfsd2aa
abcg2a
slc2a1a
tjp1a
tyrp1b
cav1
fabp11a
pecam1
cdh5
kdr
kdrl
epcam
cdh1
cldn7b
krt4
krt8
krt5
krtt1c19e
itga6b
agr2
slc4a4a
prox1a
dll4
fli1a
abcb4
cpb1
amy2a
fabp6
apoba
abcb11b
cyp3a65
fabp10a
endothelium
epidermis
ionocytes
hepatocytes
cholangiocytes
intestine
pancreas
RPE
pigment
other epith.

**fig. S23. Comparative analysis of gene expression between *Danionella cerebrum*, adult and larval zebrafish connective, muscle and epithelial tissues (A)** Dot plots showing expression patterns for a curated set of genes in connective, muscle and epithelial tissues. Larval zebrafish dataset: ZMAP (*80*); adult zebrafish dataset is collated by combining datasets from (*45, 46, 81-89*).

**Figure S24.**

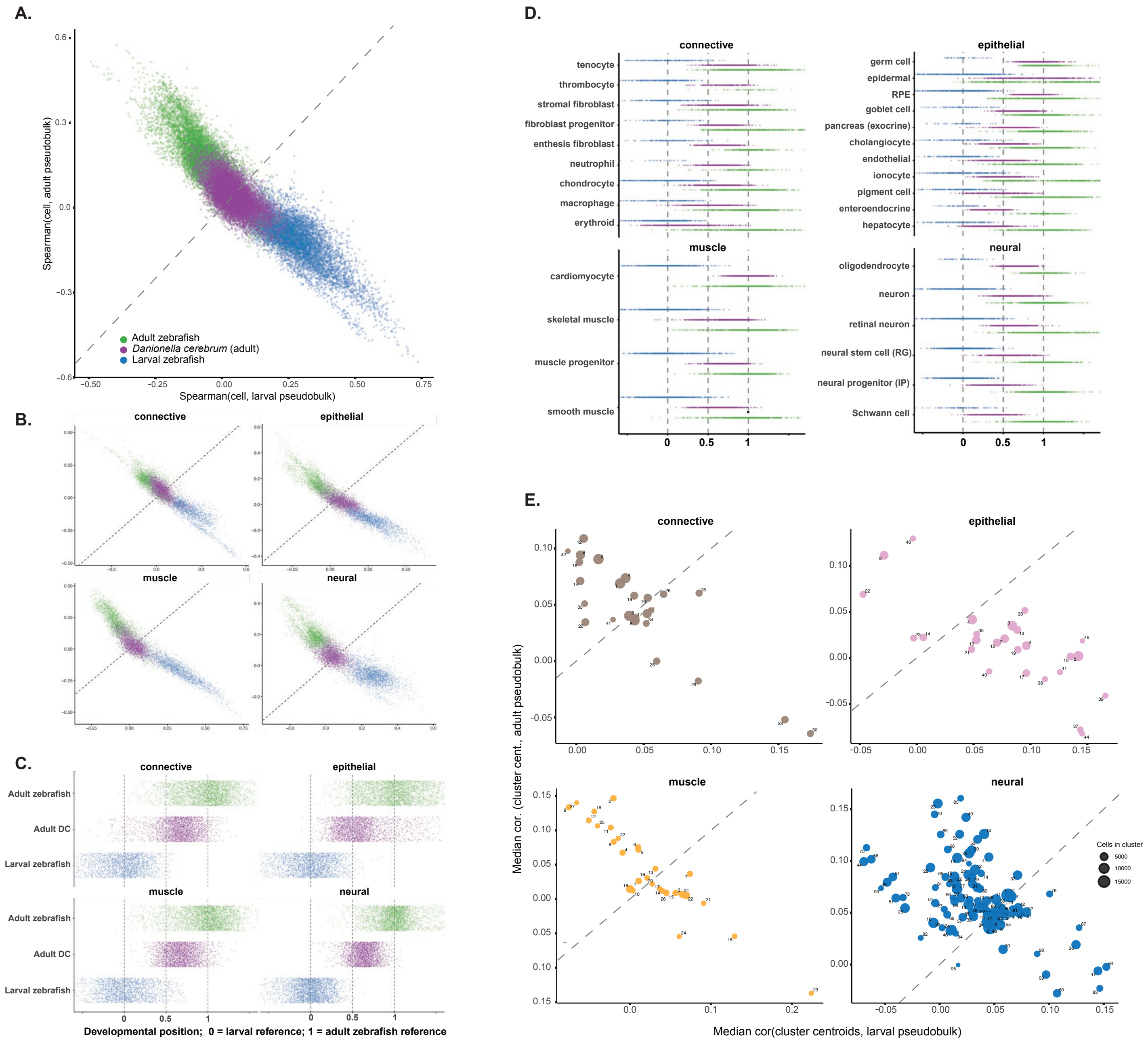

A.
Spearman(cell, adult pseudobulk)
Spearman(cell, larval pseudobulk)
Adult zebrafish
Danionella cerebrum (adult)
Larval zebrafish
B.
connective
epithelial
muscle
neural
C.
connective
epithelial
Adult zebrafish
Adult DC
Larval zebrafish
muscle
neural
Developmental position; 0 = larval reference; 1 = adult zebrafish reference
D.
connective
tenocyte
thrombocyte
stromal fibroblast
fibroblast progenitor
enthesis fibroblast
neutrophil
chondrocyte
macrophage
erythroid
epithelial
germ cell
epidermal
RPE
goblet cell
pancreas (exocrine)
cholangiocyte
endothelial
ionocyte
pigment cell
enteroendocrine
hepatocyte
muscle
cardiomyocyte
skeletal muscle
muscle progenitor
smooth muscle
neural
oligodendrocyte
neuron
retinal neuron
neural stem cell (RG)
neural progenitor (IP)
Schwann cell
E.
connective
epithelial
muscle
neural
Median cor. (cluster cent., adult pseudobulk)
Median cor(cluster centroids, larval pseudobulk)
Cells in cluster
5000
10000
15000

**fig. S24. Comparative analysis of gene expression between *Danionella cerebrum*, adult and larval zebrafish connective, muscle and epithelial tissues (A)** Comparison of all three sources (*Danionella* atlas, zebrafish larval reference atlas (Z-MAP (*80*), and pooled adult reference atlas from (*45, 46, 81-89*) in larval-adult correlation space. Each point is a cell, positioned by its Spearman correlation to the larval zebrafish cell-type pseudobulk (x-axis) versus the adult zebrafish cell-type pseudobulk (y-axis), calculated over that cell type's stage-discriminative genes, and colored by source dataset. Dashed line represents identity: cells above it correlate more strongly with the pooled adult reference and the cells below it more strongly with the larval reference. Cells from all four tissue classes are shown together, representing a "global" view. 3000 cells per tissue class are plotted. **(B)** As in (A) but split by tissue class: neural, connective, epithelial and muscle; overlaid with reference anchors. **(C)** Distribution of "developmental" position by source per tissue class. Position is scaled to ensure that "0" marks the larval reference and "1" marks the pooled adult reference. Each dot is one cell, jittered vertically within its source row. 3000 cells per source dataset are plotted. Peadomorphic shift is read as Danionella atlas cells displaced from the adult reference towards the larval (0) end. **(D)** Developmental position per cell type for all three sources is displayed, separated by tissue class and ordered by ascending median. As in (C), "0" marks the larval end and "1" marks the adult end. Each dot is one cell, and 500 cells per cell type per source dataset are shown. Photoreceptor and amacrine cells are merged into a single "retinal neuron" group and re-scored against the merged reference. **(E)** *Danionella* atlas tissue class cluster centroids in larval-adult correlation space, grouped by tissue class. Each circle represents a single atlas cluster. Clusters > 20 cells are plotted only. Clusters are placed at the median correlation of its cells to the larval (x-axis) and adult (y-axis) cell-type pseudobulk and sized by the number of cells in the cluster. A cluster's position relative to the two "anchors" indicates how larval-like versus how adult-like the cluster is.

**Figure S25.**

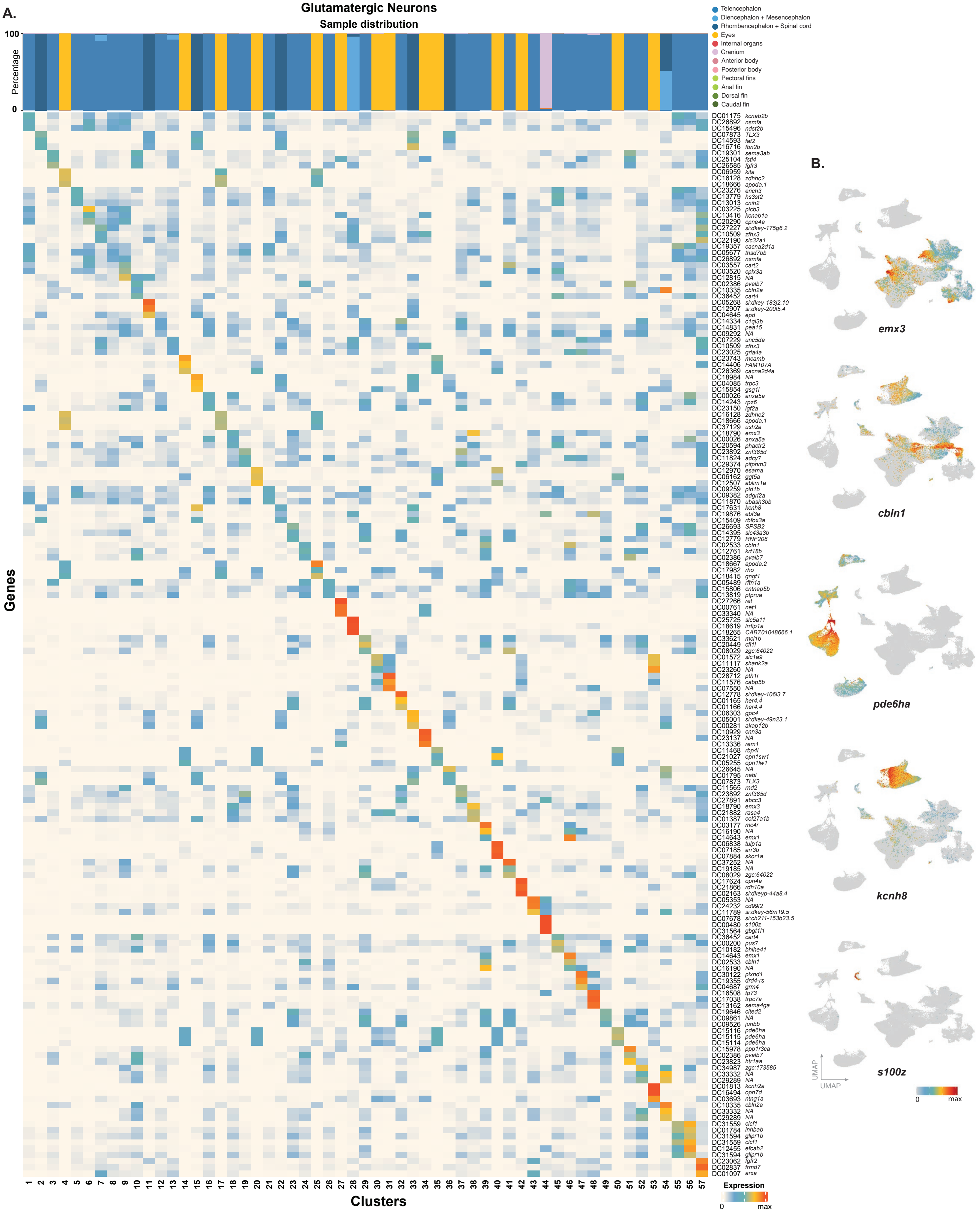
A.
Glutamatergic Neurons
Sample distribution
100
Percentage
0
Telencephalon
Diencephalon + Mesencephalon
Rhombencephalon + Spinal cord
Eyes
Internal organs
Cranium
Anterior body
Posterior body
Pectoral fins
Anal fin
Dorsal fin
Caudal fin
Genes
DC01175 kcnab2b
DC26892 nsmfa
DC15496 ndst2b
DC07873 TLX3
DC14593 fat2
DC16716 fbn2b
DC19301 sema3ab
DC25104 fstl4
DC26585 fgfr3
DC06959 kita
DC16128 zdhhc2
DC18666 apoda.1
DC23276 erich3
DC13779 hs3st2
DC13013 cnih2
DC03225 plcb3
DC13416 kcnab1a
DC20290 cpne4a
DC27227 si:dkey-175g6.2
DC10509 zfhx3
DC22190 slc32a1
DC19357 cacna2d1a
DC05677 thsd7bb
DC26892 nsmfa
DC03557 cart2
DC03520 cplx3a
DC12815 NA
DC02386 pvalb7
DC10335 cbln2a
DC36452 cart4
DC05268 si:dkey-183j2.10
DC12907 si:dkey-200l5.4
DC04645 epd
DC14334 c1ql3b
DC14831 pea15
DC09292 NA
DC07229 unc5da
DC10509 zfhx3
DC23025 gria4a
DC23743 mcamb
DC14406 FAM107A
DC26369 cacna2d4a
DC18984 NA
DC04085 trpc3
DC15854 gsg1l
DC00026 anxa5a
DC14243 rpz6
DC23150 igf2a
DC16128 zdhhc2
DC18666 apoda.1
DC37129 ush2a
DC18790 emx3
DC00026 anxa5a
DC20594 phactr2
DC23892 znf385d
DC11824 adcy7
DC29374 pitpnm3
DC12970 esama
DC06162 ggt5a
DC12507 ablim1a
DC09259 pld1b
DC09382 adgrl2a
DC11870 ubash3bb
DC17631 kcnh8
DC19876 ebf3a
DC15409 rbfox3a
DC26693 SPSB2
DC14395 slc43a3b
DC12779 RNF208
DC02533 cbln1
DC12761 krt18b
DC02386 pvalb7
DC18667 apoda.2
DC17982 rho
DC18415 gngt1
DC05489 rftn1a
DC15806 cntnap5b
DC13819 ptprua
DC27266 ret
DC00761 net1
DC33340 NA
DC25725 slc5a11
DC18619 lrrfip1a
DC18265 CABZ01048666.1
DC33621 mcl1b
DC20449 cfl1l
DC08029 zgc:64022
DC01572 slc1a9
DC11117 shank2a
DC23260 NA
DC28712 pth1r
DC11576 cabp5b
DC07550 NA
DC12778 si:dkey-106l3.7
DC01165 her4.4
DC01166 her4.4
DC06303 gpc4
DC05001 si:dkey-49n23.1
DC00281 akap12b
DC10929 cnn3a
DC23137 NA
DC13336 rem1
DC11468 rbp4l
DC21027 opn1sw1
DC05255 opn1lw1
DC26645 NA
DC01795 nebl
DC07873 TLX3
DC11565 rnd2
DC23892 znf385d
DC27891 abcc3
DC18790 emx3
DC21882 rasa4
DC01387 col27a1b
DC03177 mc4r
DC16190 NA
DC14643 emx1
DC06838 tulp1a
DC07185 arr3b
DC07884 skor1a
DC37252 NA
DC19185 NA
DC08029 zgc:64022
DC17624 opn4a
DC21866 rdh10a
DC02163 si:dkeyp-44a8.4
DC05353 NA
DC24232 cd99l2
DC11789 si:dkey-56m19.5
DC07678 si:ch211-153b23.5
DC00480 s100z
DC31564 gbgt1l1
DC36452 cart4
DC00200 pus7
DC10182 bhlhe41
DC14643 emx1
DC02533 cbln1
DC16190 NA
DC30122 plxnd1
DC19355 drd4-rs
DC04687 grm4
DC16508 tp73
DC17038 trpc7a
DC13162 sema4ga
DC19646 cited2
DC09861 NA
DC09526 junbb
DC15116 pde6ha
DC15115 pde6ha
DC15114 pde6ha
DC15978 ppp1r3ca
DC02386 pvalb7
DC23823 htr1aa
DC34987 zgc:173585
DC33332 NA
DC29289 NA
DC18813 kcnh2a
DC16494 opn7d
DC03693 ntng1a
DC10335 cbln2a
DC33332 NA
DC29289 NA
DC31559 clcf1
DC01784 inhbab
DC31594 glipr1b
DC31559 clcf1
DC12455 efcab2
DC31594 glipr1b
DC23062 fgfr2
DC02837 frmd7
DC01097 arxa
1 2 3 4 5 6 7 8 9 10 11 12 13 14 15 16 17 18 19 20 21 22 23 24 25 26 27 28 29 30 31 32 33 34 35 36 37 38 39 40 41 42 43 44 45 46 47 48 49 50 51 52 53 54 55 56 57
Clusters
Expression
0
max
B.
emx3
cbln1
pde6ha
kcnh8
s100z
UMAP
UMAP
0
max

**fig. S25. Characterization of glutamatergic neurons (A)** Top three markers and sample c for glutamatergic neuron clusters are shown. **(B)** Feature plots of genes enriched in telencephalon, eye, rhombencephalon and spinal cord, and peripheral glutamatergic neuron clusters. Top columns represent the distribution of experimental samples across the clusters.

**Figure S26.**

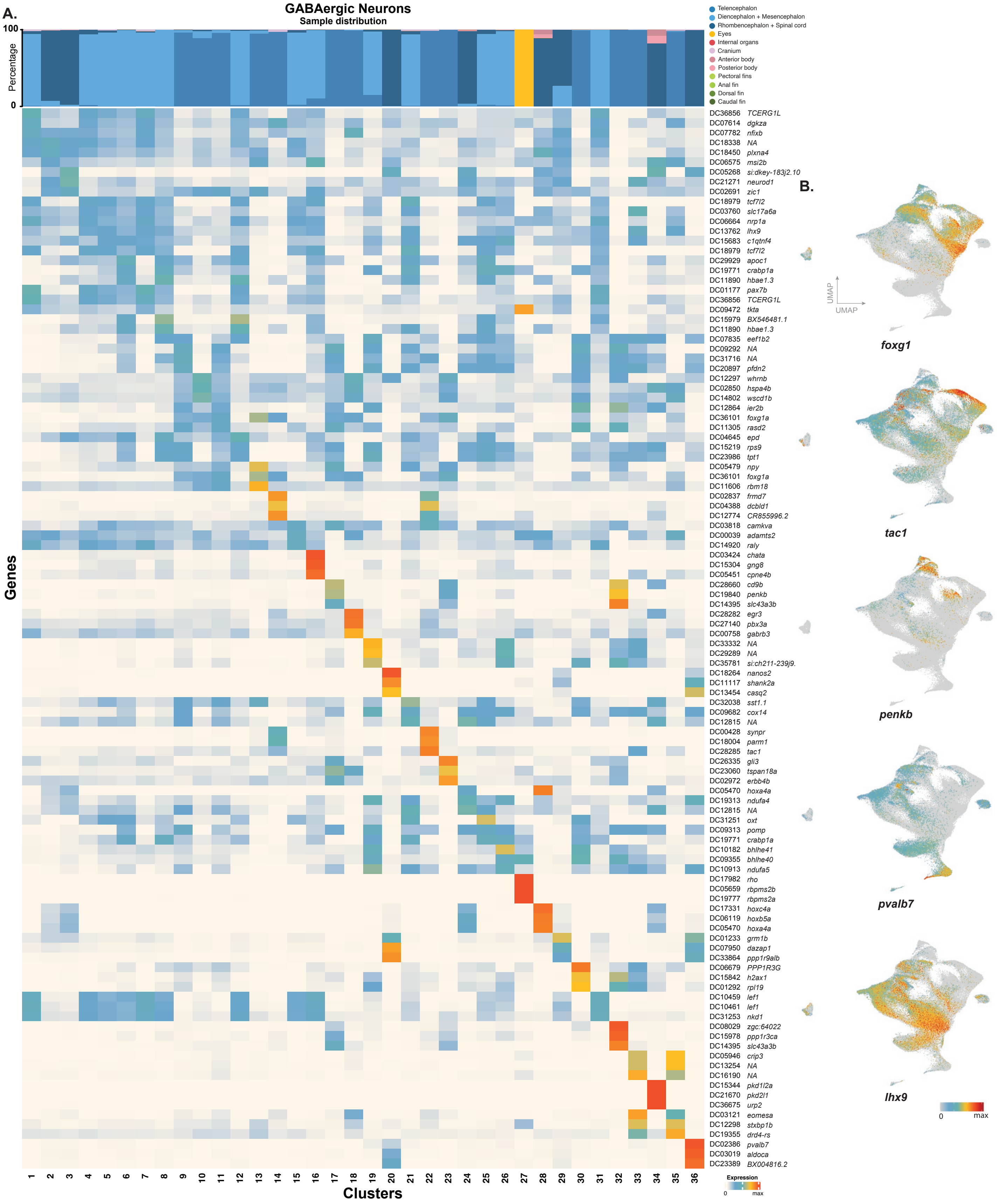

A.
GABAergic Neurons
Sample distribution
Percentage
100
0
Telencephalon
Diencephalon + Mesencephalon
Rhombencephalon + Spinal cord
Eyes
Internal organs
Cranium
Anterior body
Posterior body
Pectoral fins
Anal fin
Dorsal fin
Caudal fin
Genes
DC36856 TCERG1L
DC07614 dgkza
DC07782 nfixb
DC18338 NA
DC18450 plxna4
DC06575 msi2b
DC05268 si:dkey-183j2.10
DC21271 neurod1
DC02691 zic1
DC18979 tcf7l2
DC03760 slc17a6a
DC06664 nrp1a
DC13762 lhx9
DC15683 c1qtnf4
DC18979 tcf7l2
DC29929 apoc1
DC19771 crabp1a
DC11890 hbae1.3
DC01177 pax7b
DC36856 TCERG1L
DC09472 tkta
DC15979 BX546481.1
DC11890 hbae1.3
DC07835 eef1b2
DC09292 NA
DC31716 NA
DC20897 pfdn2
DC12297 whrnb
DC02850 hspa4b
DC14802 wscd1b
DC12864 ier2b
DC36101 foxg1a
DC11305 rasd2
DC04645 epd
DC15219 rps9
DC23986 tpt1
DC05479 npy
DC36101 foxg1a
DC11606 rbm18
DC02837 frmd7
DC04388 dcbld1
DC12774 CR855996.2
DC03818 camkva
DC00039 adamts2
DC14920 raly
DC03424 chata
DC15304 gng8
DC05451 cpne4b
DC28660 cd9b
DC19840 penkb
DC14395 slc43a3b
DC28282 egr3
DC27140 pbx3a
DC00758 gabrb3
DC33332 NA
DC29289 NA
DC35781 si:ch211-239j9.
DC18264 nanos2
DC11117 shank2a
DC13454 casq2
DC32038 sst1.1
DC09682 cox14
DC12815 NA
DC00428 synpr
DC18004 parm1
DC28285 tac1
DC26335 gli3
DC23060 tspan18a
DC02972 erbb4b
DC05470 hoxa4a
DC19313 ndufa4
DC12815 NA
DC31251 oxt
DC09313 pomp
DC19771 crabp1a
DC10182 bhlhe41
DC09355 bhlhe40
DC10913 ndufa5
DC17982 rho
DC05659 rbpms2b
DC19777 rbpms2a
DC17331 hoxc4a
DC06119 hoxb5a
DC05470 hoxa4a
DC01233 grm1b
DC07950 dazap1
DC33864 ppp1r9alb
DC06679 PPP1R3G
DC15842 h2ax1
DC01292 rpl19
DC10459 lef1
DC10461 lef1
DC31253 nkd1
DC08029 zgc:64022
DC15978 ppp1r3ca
DC14395 slc43a3b
DC05946 crip3
DC13254 NA
DC16190 NA
DC15344 pkd1l2a
DC21670 pkd2l1
DC36675 urp2
DC03121 eomesa
DC12298 stxbp1b
DC19355 drd4-rs
DC02386 pvalb7
DC03019 aldoca
DC23389 BX004816.2
1 2 3 4 5 6 7 8 9 10 11 12 13 14 15 16 17 18 19 20 21 22 23 24 25 26 27 28 29 30 31 32 33 34 35 36
Clusters
Expression
0
max
B.
UMAP
UMAP
foxg1
tac1
penkb
pvalb7
lhx9
0
max

**fig. S26. Characterization of GABAergic neurons (A)** Top three markers and sample distribution percentages for GABAergic neuron clusters are shown. **(B)** Feature plots of genes enriched in distinct experimental samples and GABAergic neuron clusters are shown. Top columns represent the distribution of experimental samples across the clusters.

**Figure S27.**

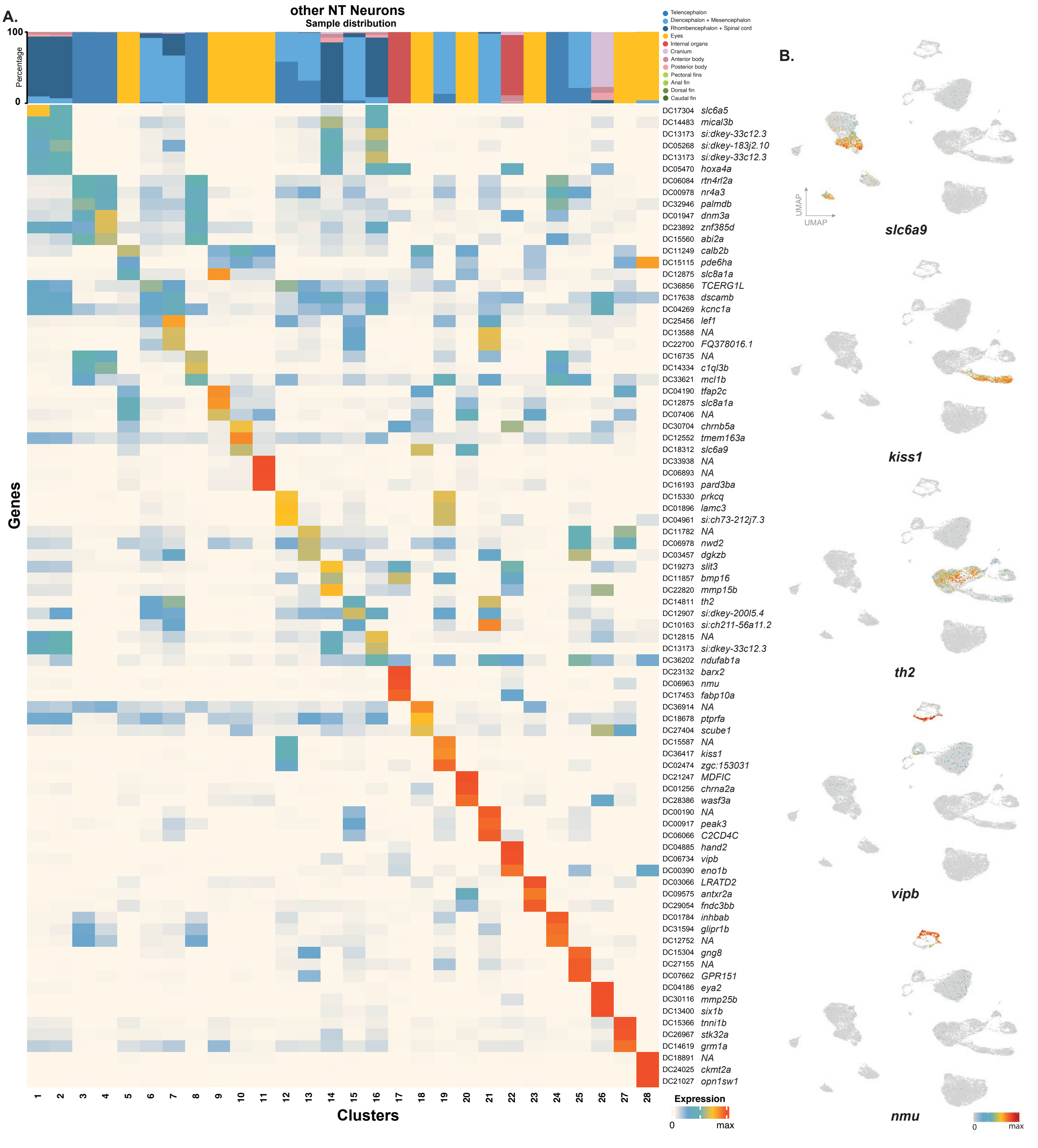
A.
other NT Neurons
Sample distribution
Telencephalon
Diencephalon + Mesencephalon
Rhombencephalon + Spinal cord
Eyes
Internal organs
Cranium
Anterior body
Posterior body
Pectoral fins
Anal fin
Dorsal fin
Caudal fin
100
Percentage
0
Genes
DC17304 slc6a5
DC14483 mical3b
DC13173 si:dkey-33c12.3
DC05268 si:dkey-183j2.10
DC13173 si:dkey-33c12.3
DC05470 hoxa4a
DC06084 rtn4rl2a
DC00978 nr4a3
DC32946 palmdb
DC01947 dnm3a
DC23892 znf385d
DC15560 abi2a
DC11249 calb2b
DC15115 pde6ha
DC12875 slc8a1a
DC36856 TCERG1L
DC17638 dscamb
DC04269 kcnc1a
DC25456 lef1
DC13588 NA
DC22700 FQ378016.1
DC16735 NA
DC14334 c1ql3b
DC33621 mcl1b
DC04190 tfap2c
DC12875 slc8a1a
DC07406 NA
DC30704 chrnb5a
DC12552 tmem163a
DC18312 slc6a9
DC33938 NA
DC06893 NA
DC16193 pard3ba
DC15330 prkcq
DC01896 lamc3
DC04961 si:ch73-212j7.3
DC11782 NA
DC06978 nwd2
DC03457 dgkzb
DC19273 slit3
DC11857 bmp16
DC22820 mmp15b
DC14811 th2
DC12907 si:dkey-200l5.4
DC10163 si:ch211-56a11.2
DC12815 NA
DC13173 si:dkey-33c12.3
DC36202 ndufab1a
DC23132 barx2
DC06963 nmu
DC17453 fabp10a
DC36914 NA
DC18678 ptprfa
DC27404 scube1
DC15587 NA
DC36417 kiss1
DC02474 zgc:153031
DC21247 MDFIC
DC01256 chrna2a
DC28386 wasf3a
DC00190 NA
DC00917 peak3
DC06066 C2CD4C
DC04885 hand2
DC06734 vipb
DC00390 eno1b
DC03066 LRATD2
DC09575 antxr2a
DC29054 fndc3bb
DC01784 inhbab
DC31594 glipr1b
DC12752 NA
DC15304 gng8
DC27155 NA
DC07662 GPR151
DC04186 eya2
DC30116 mmp25b
DC13400 six1b
DC15366 tnni1b
DC26967 stk32a
DC14619 grm1a
DC18891 NA
DC24025 ckmt2a
DC21027 opn1sw1
1 2 3 4 5 6 7 8 9 10 11 12 13 14 15 16 17 18 19 20 21 22 23 24 25 26 27 28
Clusters
Expression
0
max
B.
UMAP
UMAP
slc6a9
kiss1
th2
vipb
nmu
0
max

**Fig. S27. Characterization of neurons of other neurotransmitter classes (A)** Top three markers and sample distribution percentages within clusters of neurons of other neurotransmitter identities, including glycinergic, histaminergic, serotonergic, dopaminergic, neuropeptidergic neurons are shown. **(B)** Feature plots of select genes enriched in distinct experimental samples and clusters are shown. Top columns represent the distribution of experimental samples across the clusters.

**Figure S28.**

**A.** **Neurotransmitter class-specific transcription factors**

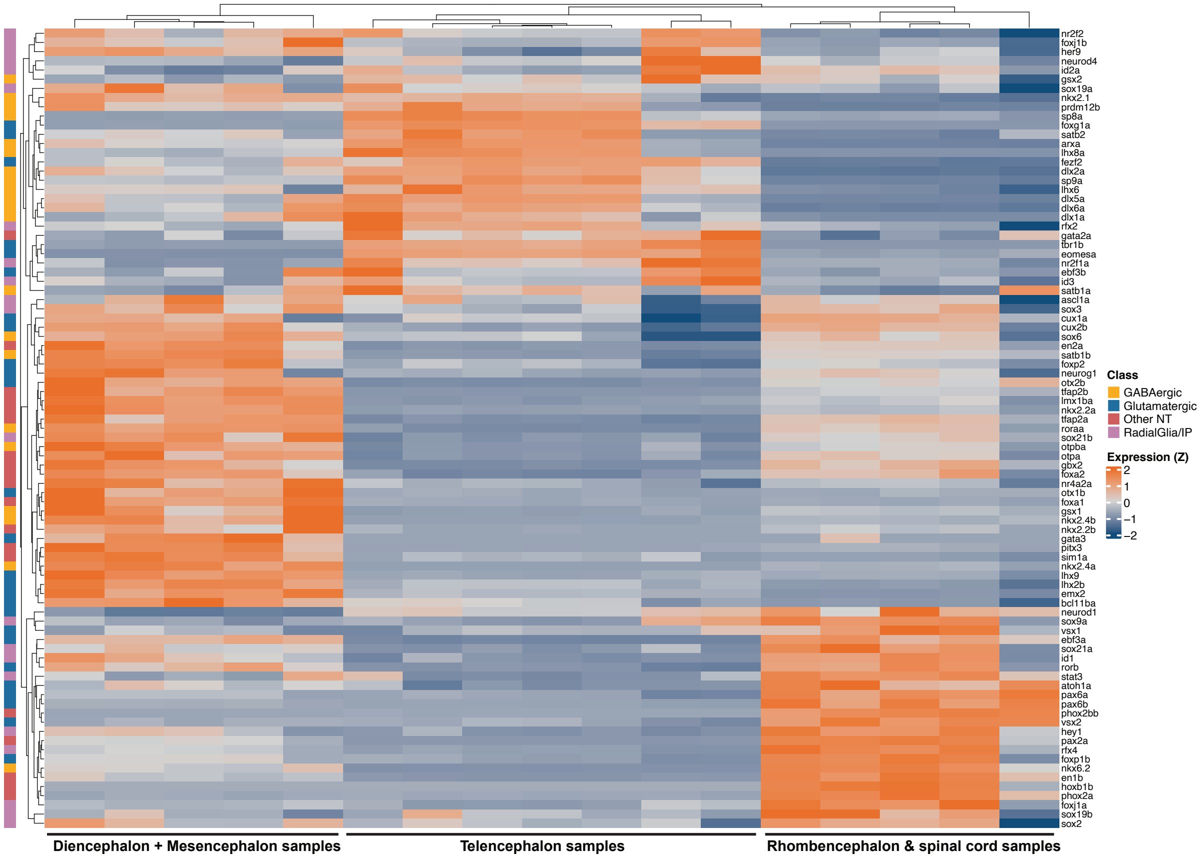

**B.** **Neurotransmitter class-specific NT synthesis/transmission; adhesion/synaptic; cellular homeostasis-related genes**

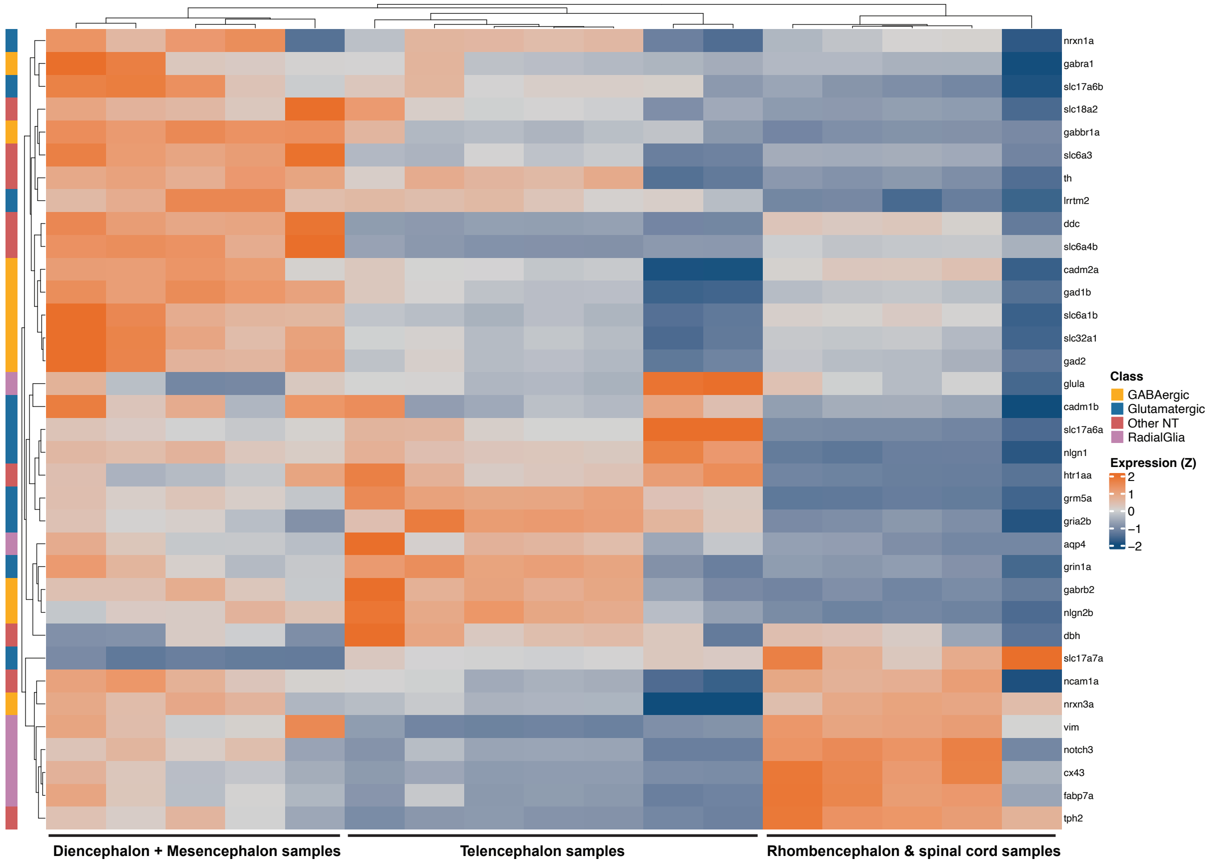

**fig. S28. Transcriptional factors, neurotransmitter synthesis and cell-cell adhesion genes with regional expression in the CNS (A)** Heatmap showing expression of transcriptional factors with regional expression profiles. (B) Heatmap showing genes linked with neurotransmitter synthesis and transmission, cell-cell adhesion genes and cellular homeostasis-related genes with regional expression patterns.

**Figure S29.**

**A.**

Telencephalon vs. Dien-+Mesencephalon (pseudobulk - all cells)

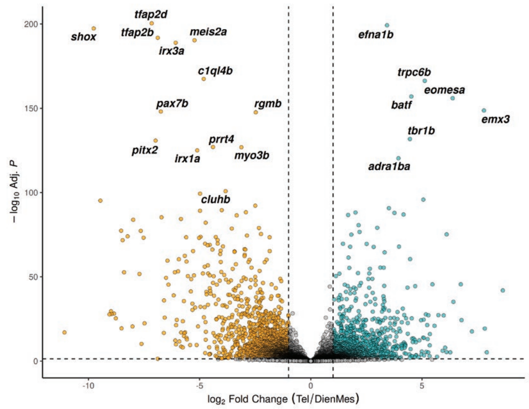

Telencephalon vs. Rhombencephalon + Spinal C. (pseudobulk - all cells)

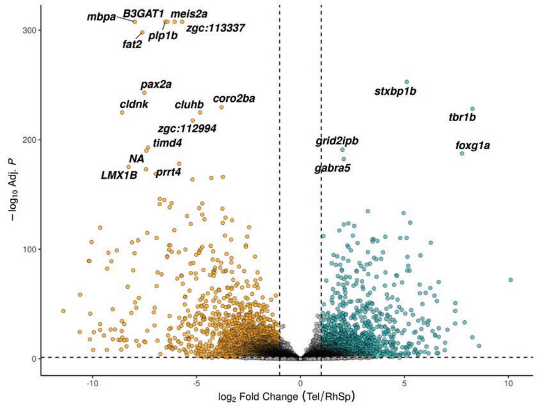

Dien-Mesencephalon vs. Rhombencephalon + Spinal C. (pseudobulk - all cells )

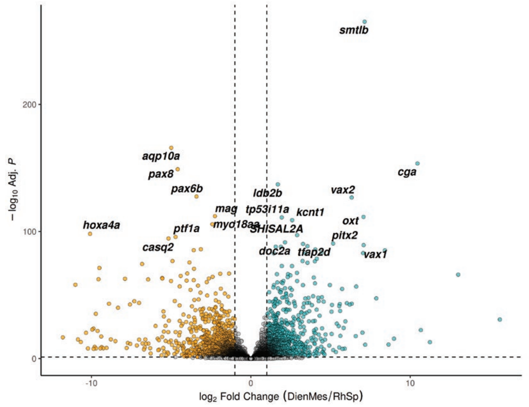

**B.**

Telencephalon vs. Dien-+Mesencephalon (pseudobulk - *snap25a*+)

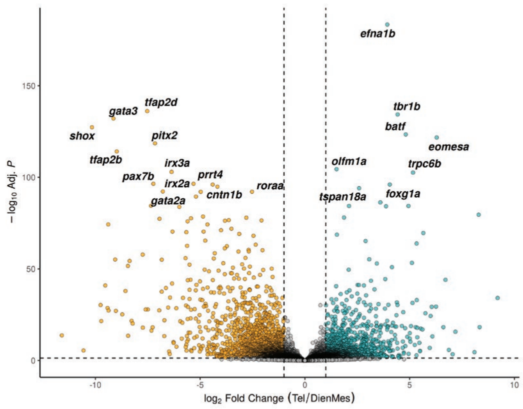

Telencephalon vs. Rhombencephalon + Spinal C. (pseudobulk - *snap25a*+)

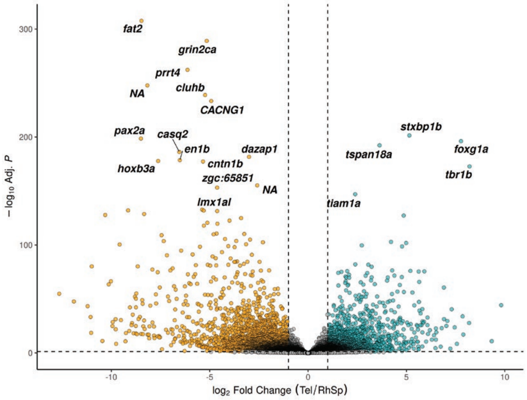

Dien-Mesencephalon vs. Rhombencephalon + Spinal C. (pseudobulk - *snap25a*+)

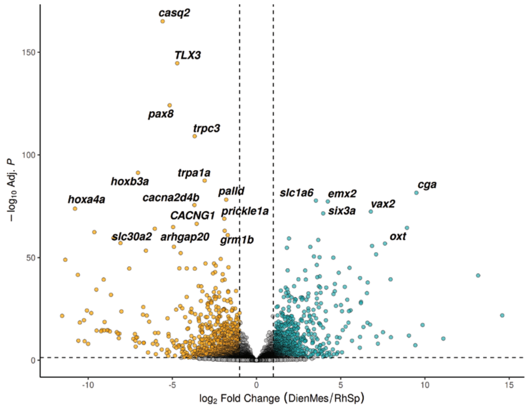

**fig. S29. Differential expression analysis comparing different regions across CNS (A)** Volcano plots showing top differentially expressed genes (pseudobulk analysis) across telencephalon, diencephalon and mesencephalon, and rhombencephalon and spinal cord regions. **(B)** Volcano plots showing top differentially expressed genes (pseudobulk analysis) in *snap25a*+ mature neurons across telencephalon, diencephalon and mesencephalon, and rhombencephalon and spinal cord cells.

Figure S30.

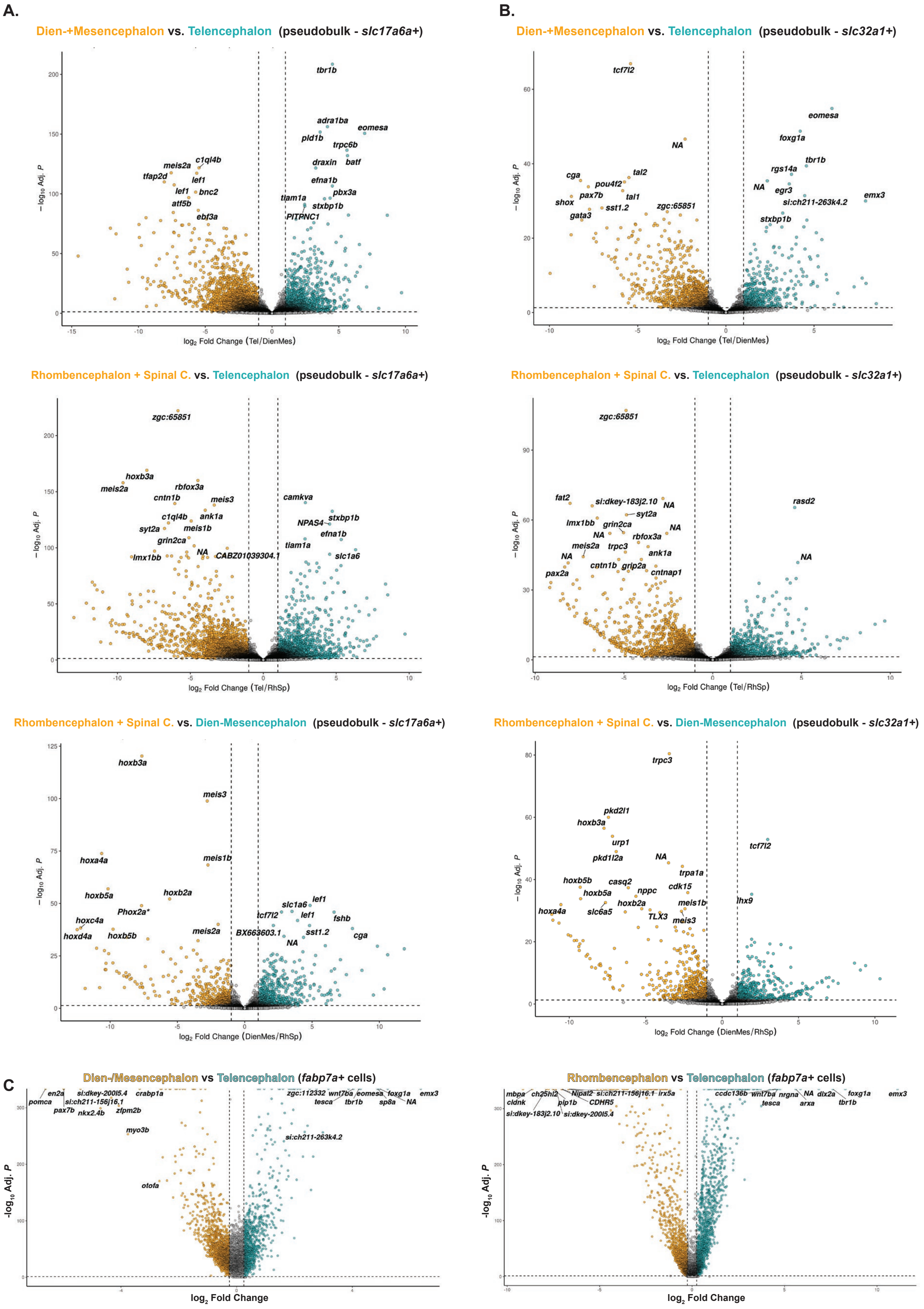

A.
Dien-+Mesencephalon vs. Telencephalon (pseudobulk - slc17a6a+)
−log10 Adj. P
log2 Fold Change (Tel/DienMes)
tbr1b
adra1ba
eomesa
pld1b
trpc6b
draxin
batf
efna1b
pbx3a
tiam1a
stxbp1b
PITPNC1
c1ql4b
meis2a
tfap2d
lef1
bnc2
atf5b
ebf3a
B.
Dien-+Mesencephalon vs. Telencephalon (pseudobulk - slc32a1+)
−log10 Adj. P
log2 Fold Change (Tel/DienMes)
tcf7l2
eomesa
foxg1a
NA
tbr1b
rgs14a
NA
egr3
emx3
si:ch211-263k4.2
stxbp1b
cga
tal2
pou4f2
pax7b
tal1
shox
sst1.2
zgc:65851
gata3
Rhombencephalon + Spinal C. vs. Telencephalon (pseudobulk - slc17a6a+)
−log10 Adj. P
log2 Fold Change (Tel/RhSp)
zgc:65851
hoxb3a
meis2a
rbfox3a
cntn1b
meis3
camkva
c1ql4b
ank1a
NPAS4
stxbp1b
syt2a
meis1b
efna1b
grin2ca
tiam1a
NA
CABZ01039304.1
slc1a6
lmx1bb
Rhombencephalon + Spinal C. vs. Telencephalon (pseudobulk - slc32a1+)
−log10 Adj. P
log2 Fold Change (Tel/RhSp)
zgc:65851
fat2
si:dkey-183j2.10
NA
rasd2
syt2a
lmx1bb
grin2ca
NA
NA
rbfox3a
meis2a
trpc3
ank1a
NA
NA
cntn1b
grip2a
cntnap1
pax2a
Rhombencephalon + Spinal C. vs. Dien-Mesencephalon (pseudobulk - slc17a6a+)
−log10 Adj. P
log2 Fold Change (DienMes/RhSp)
hoxb3a
meis3
hoxa4a
meis1b
hoxb5a
hoxb2a
lef1
slc1a6
Phox2a*
tcf7l2
lef1
fshb
hoxc4a
hoxd4a
hoxb5b
meis2a
BX663603.1
sst1.2
cga
NA
Rhombencephalon + Spinal C. vs. Dien-Mesencephalon (pseudobulk - slc32a1+)
−log10 Adj. P
log2 Fold Change (DienMes/RhSp)
trpc3
pkd2l1
hoxb3a
tcf7l2
urp1
pkd1l2a
NA
trpa1a
hoxb5b
casq2
cdk15
hoxb5a
nppc
lhx9
hoxb2a
meis1b
hoxa4a
slc6a5
TLX3
meis3
C
Dien-/Mesencephalon vs Telencephalon (fabp7a+ cells)
-log10 Adj. P
log2 Fold Change
en2a
si:dkey-200l5.4
crabp1a
zgc:112332
wnt7ba
eomesa
foxg1a
emx3
pomca
si:ch211-156j16.1
tesca
tbr1b
sp8a
NA
pax7b
nkx2.4b
zfpm2b
myo3b
si:ch211-263k4.2
otofa
Rhombencephalon vs Telencephalon (fabp7a+ cells)
-log10 Adj. P
log2 Fold Change
mbpa
ch25hl2
Nipal2
si:ch211-156j16.1
irx5a
ccdc136b
wnt7ba
nrgna
NA
dlx2a
foxg1a
emx3
cldnk
plp1b
CDHR5
tesca
arxa
tbr1b
si:dkey-183j2.10
si:dkey-200l5.4

**fig. S30. Differential expression analysis comparing different regions across CNS (A)** Volcano plots showing top differentially expressed genes (pseudobulk analysis) in *slc17a6a*+ glutamatergic neurons across telencephalon, diencephalon and mesencephalon, and rhombencephalon and spinal cord cells. **(B)** Volcano plots showing top differentially expressed genes (pseudobulk analysis) in *slc32a1*+ GABAergic neurons across telencephalon, diencephalon and mesencephalon, and rhombencephalon and spinal cord cells. **(C)** Volcano plots demonstrating the top differentially expressed genes between *fabp7a*+ radial glia from telencephalon, diencephalon and mesencephalon, and rhombencephalon and spinal cord.

Figure S31.

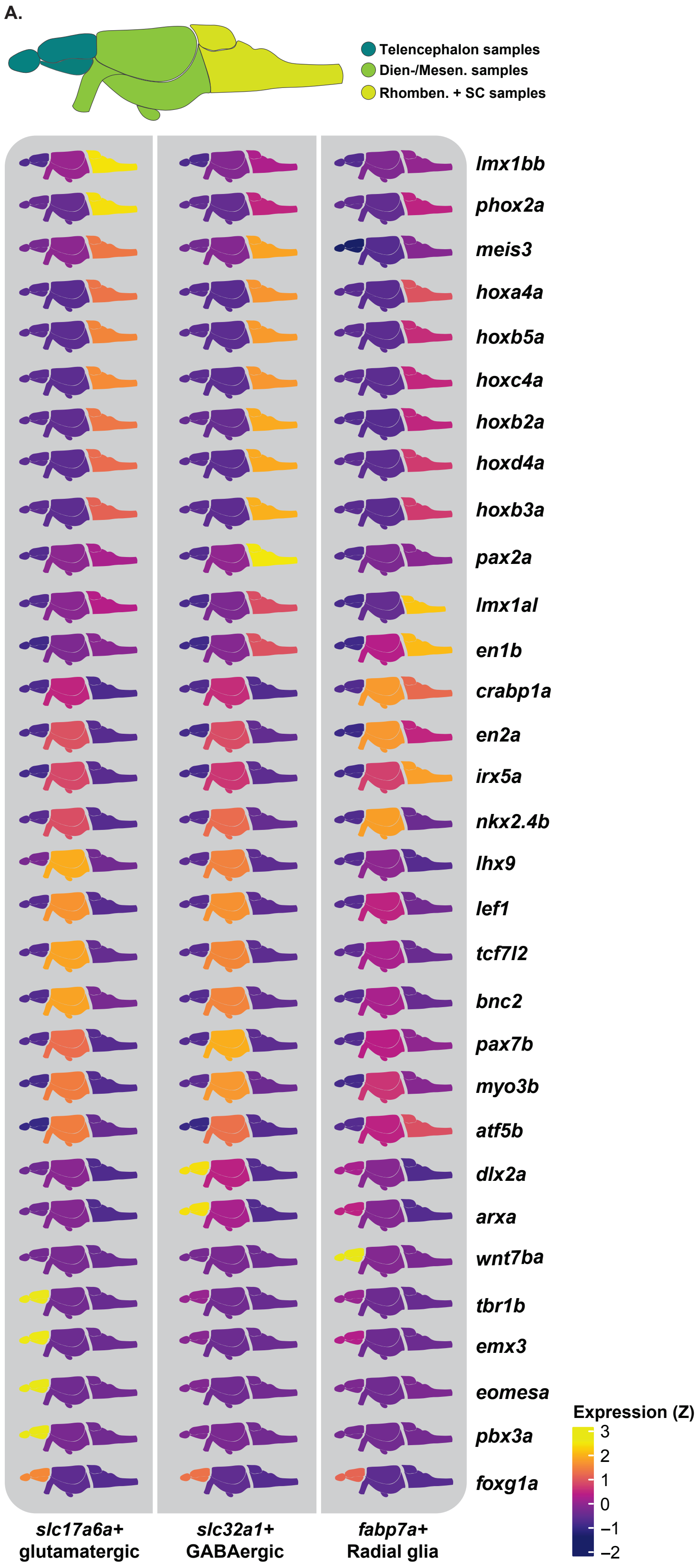

A.
Telencephalon samples
Dien-/Mesen. samples
Rhomben. + SC samples
lmx1bb
phox2a
meis3
hoxa4a
hoxb5a
hoxc4a
hoxb2a
hoxd4a
hoxb3a
pax2a
lmx1al
en1b
crabp1a
en2a
irx5a
nkx2.4b
lhx9
lef1
tcf7l2
bnc2
pax7b
myo3b
atf5b
dlx2a
arxa
wnt7ba
tbr1b
emx3
eomesa
pbx3a
foxg1a
slc17a6a+ glutamatergic
slc32a1+ GABAergic
fabp7a+ Radial glia
Expression (Z)
3
2
1
0
−1
−2

**fig. S31. Top differentially expressed genes in glutamatergic neurons, GABAergic neurons and radial glia across CNS regions. (A)** Expanded summary heatmap showing expression patterns for top differentially expressed genes for *slc17a6a*+ glutamatergic neurons, *slc32a1*+ GABAergic neurons, and *fabp7a*+ radial glia across regions (summary version of this heatmap is presented in Fig. 4I).

**Figure S32.**

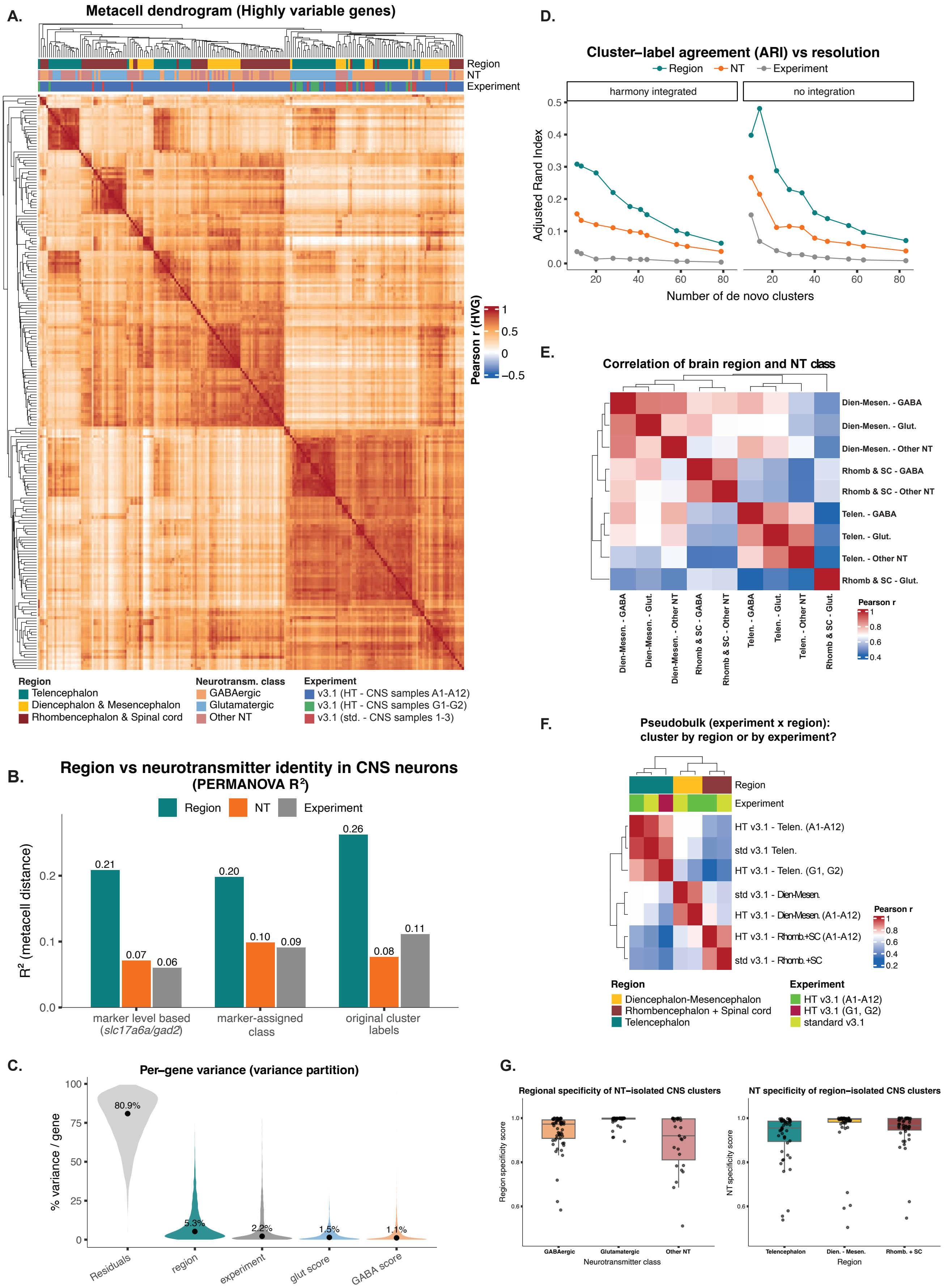

A.
Metacell dendrogram (Highly variable genes)
Region
NT
Experiment
Pearson r (HVG)
1
0.5
0
−0.5
Region
Telencephalon
Diencephalon & Mesencephalon
Rhombencephalon & Spinal cord
Neurotransm. class
GABAergic
Glutamatergic
Other NT
Experiment
v3.1 (HT - CNS samples A1-A12)
v3.1 (HT - CNS samples G1-G2)
v3.1 (std. - CNS samples 1-3)
B.
Region vs neurotransmitter identity in CNS neurons
(PERMANOVA R²)
Region
NT
Experiment
R² (metacell distance)
0.21
0.07
0.06
0.20
0.10
0.09
0.26
0.08
0.11
marker level based (slc17a6a/gad2)
marker-assigned class
original cluster labels
C.
Per−gene variance (variance partition)
% variance / gene
80.9%
5.3%
2.2%
1.5%
1.1%
Residuals
region
experiment
glut score
GABA score
D.
Cluster−label agreement (ARI) vs resolution
Region
NT
Experiment
harmony integrated
no integration
Adjusted Rand Index
Number of de novo clusters
E.
Correlation of brain region and NT class
Dien-Mesen. - GABA
Dien-Mesen. - Glut.
Dien-Mesen. - Other NT
Rhomb & SC - GABA
Rhomb & SC - Other NT
Telen. - GABA
Telen. - Glut.
Telen. - Other NT
Rhomb & SC - Glut.
Pearson r
F.
Pseudobulk (experiment x region):
cluster by region or by experiment?
Region
Experiment
HT v3.1 - Telen. (A1-A12)
std v3.1 Telen.
HT v3.1 - Telen. (G1, G2)
std v3.1 - Dien-Mesen.
HT v3.1 - Dien-Mesen. (A1-A12)
HT v3.1 - Rhomb.+SC (A1-A12)
std v3.1 - Rhomb.+SC
Pearson r
Region
Diencephalon-Mesencephalon
Rhombencephalon + Spinal cord
Telencephalon
Experiment
HT v3.1 (A1-A12)
HT v3.1 (G1, G2)
standard v3.1
G.
Regional specificity of NT−isolated CNS clusters
NT specificity of region−isolated CNS clusters
Region specificity score
NT specificity score
GABAergic
Glutamatergic
Other NT
Neurotransmitter class
Telencephalon
Dien. - Mesen.
Rhomb. + SC
Region

**fig. S32. Contribution of regional origin and neurotransmitter identity to neuronal transcriptomic similarities (A)** Metacell dendrogram showing hierarchical clustering of 206 neuronal metacells, using highly variable genes. Top bars annotate each metacell's brain region, neurotransmitter (NT) class, and experiment of origin. Analysis shows that major branches are organized by region, with different NT classes interleaved within region blocks. **(B)** PERMANOVA analysis showing region versus neurotransmitter contribution to transcriptomic identity. Bars indicate the share of variation in neuronal gene expression profiles explained by brain region, NT class, and experiment (included to account for possible batch effects), where a larger bar represents a larger contribution. NT is separately shown under three increasingly strict ways of analysis: graded marker expression (where each neuron type's expression level of NT genes was used directly as a continuous value with no category assigned), a marker-based category (where a discrete GABA/glut/other NT identity label was assigned when one neurotransmitter predominated and ambiguous types left unlabeled) and the original cluster labels used in this study (where the NT identity inherited from original cluster labels were used as is), to indicate that results of the analysis does not depend on how NT class is defined. Regional contribution appears stronger than NT under all three definitions. **(C)** Per-gene variance decomposition analysis showing, for each of the 3000 highly variable genes, the percentage of expression variance attributable to region, NT marker scores, experiment of origin, and residual variation. For each gene, a linear model (using the variancePartition package in R) partitioned its expression variation across metacells into fractions attributable to each term (region, NT score, or experiment). The sum of squares attributable to region, to each NT class marker score, to experiment, and to the leftover/residual, expressed as percentages that add up to a complete 100%. The violin plot shows the distribution across genes, and the dots indicate medians. Most of the observed variance is "residual" (i.e., driven by currently unexplained fine-grained cell-type differences rather than by region or NT), and region is the strongest component. **(D)** Cluster-label agreement (adjusted Rand index) analysis showing agreement between de novo cell clusters and the region, NT, and experiment labels, measured across increasing clustering resolutions, in the uncorrected/no integration (PCA) and batch-integrated (Harmony, batch=experiment) embeddings. Region agreement exceeds NT at all resolutions tested. **(E)** Pairwise transcriptome correlation of pseudobulk groups, annotated by NT class and brain region, indicating similarities of different NT class neurons within the same regions. **(F)** Pairwise correlation of pseudobulk profiles aggregated per experiment x region (without Harmony integration), labeled by region and experiment/chemistry. Profiles cluster by anatomical region rather than by experiment/chemistry. Additionally, each region's profile in one experiment is most correlated with the same region in the other, rendering regional gene expression signatures as a main biological driver of transcriptomic similarities. **(G)** Distribution of specificity scores for de novo clusters built within a single NT class. Clusters of each NT class scored for region specificity (fraction of cells dominantly present in a region) (left); clusters formed within a specific region scored for NT specificity. Each point is a cluster; boxes indicate median and interquartile ranges.

Figure S33.

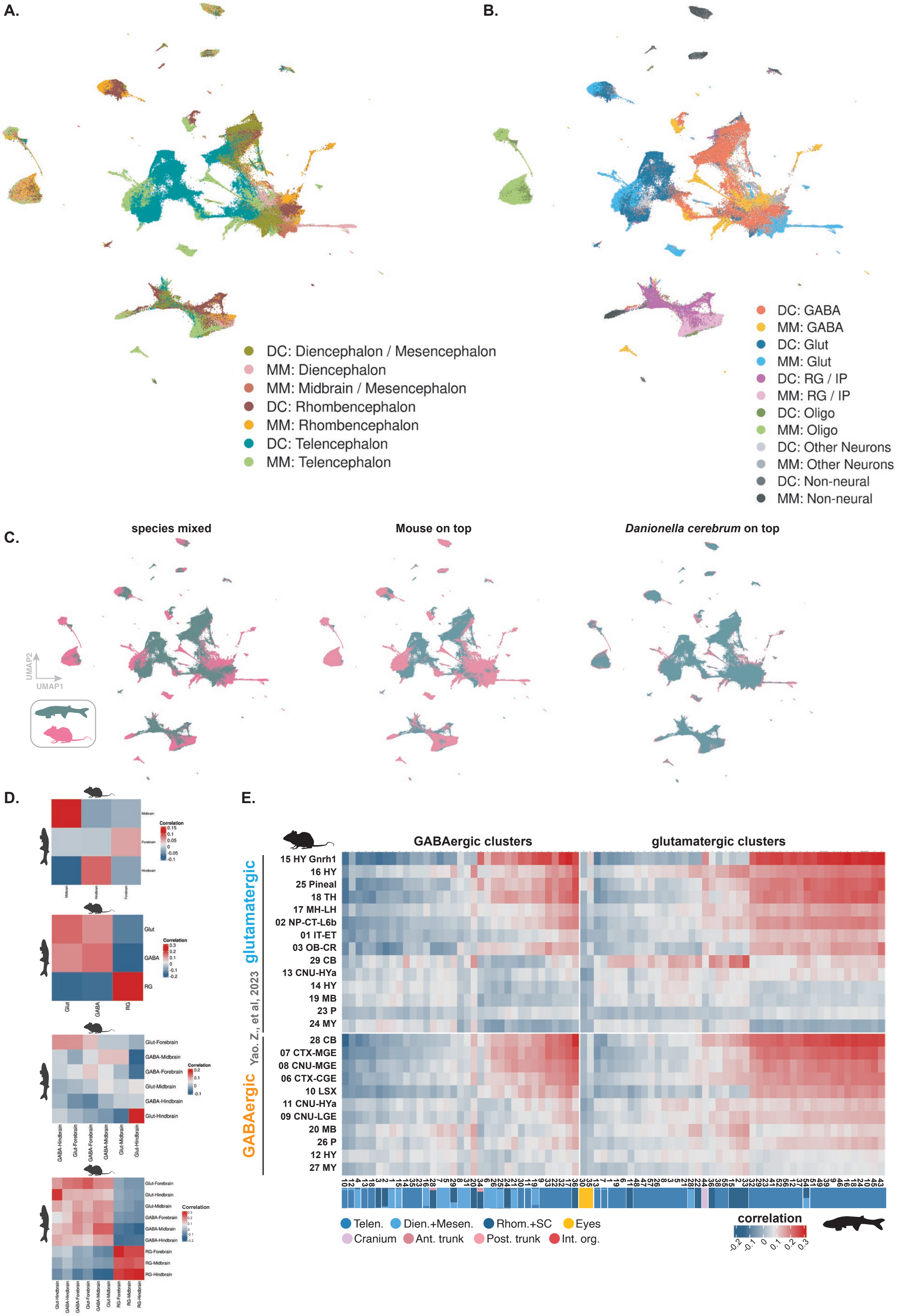
A.
B.
DC: Diencephalon / Mesencephalon
MM: Diencephalon
MM: Midbrain / Mesencephalon
DC: Rhombencephalon
MM: Rhombencephalon
DC: Telencephalon
MM: Telencephalon
DC: GABA
MM: GABA
DC: Glut
MM: Glut
DC: RG / IP
MM: RG / IP
DC: Oligo
MM: Oligo
DC: Other Neurons
MM: Other Neurons
DC: Non-neural
MM: Non-neural
C.
species mixed
Mouse on top
Danionella cerebrum on top
UMAP2
UMAP1
D.
Glut
GABA
RG
Correlation
E.
GABAergic clusters
glutamatergic clusters
glutamatergic
GABAergic
Yao. Z., et al, 2023
15 HY Gnrh1
16 HY
25 Pineal
18 TH
17 MH-LH
02 NP-CT-L6b
01 IT-ET
03 OB-CR
29 CB
13 CNU-HYa
14 HY
19 MB
23 P
24 MY
28 CB
07 CTX-MGE
08 CNU-MGE
06 CTX-CGE
10 LSX
11 CNU-HYa
09 CNU-LGE
20 MB
26 P
12 HY
27 MY
Telen.
Dien.+Mesen.
Rhom.+SC
Eyes
Cranium
Ant. trunk
Post. trunk
Int. org.
correlation
-0.2
-0.1
0
0.1
0.2
0.3

**fig. S33. Comparative analysis between *Danionella cerebrum* and mouse neurotransmitter classes and brain regions (A)** UMAP plots showing cross-species correlation (SAMap) marked by region and **(B)** by neurotransmitter class between mouse (data source (*13*) and *Danionella* neural classes. **(C)** UMAP indicating interspecies mixing between the clusters from mouse and *Danionella* neural classes **(D)** Correlation matrix heatmap (SAMap) showing similarity between distinct regions; neurotransmitter classes; neurotransmitter classes from different regions; and neurotransmitter classes and radial glia from different regions, respectively/from top to bottom **(E)** Correlation matrix heatmap (SAMap) showing similarity between glutamatergic and GABAergic neurons from three distinct regions across *Danionella* CNS.

**Figure S34.**

**A.** **Glutamatergic neurons - cross-species comparsion**

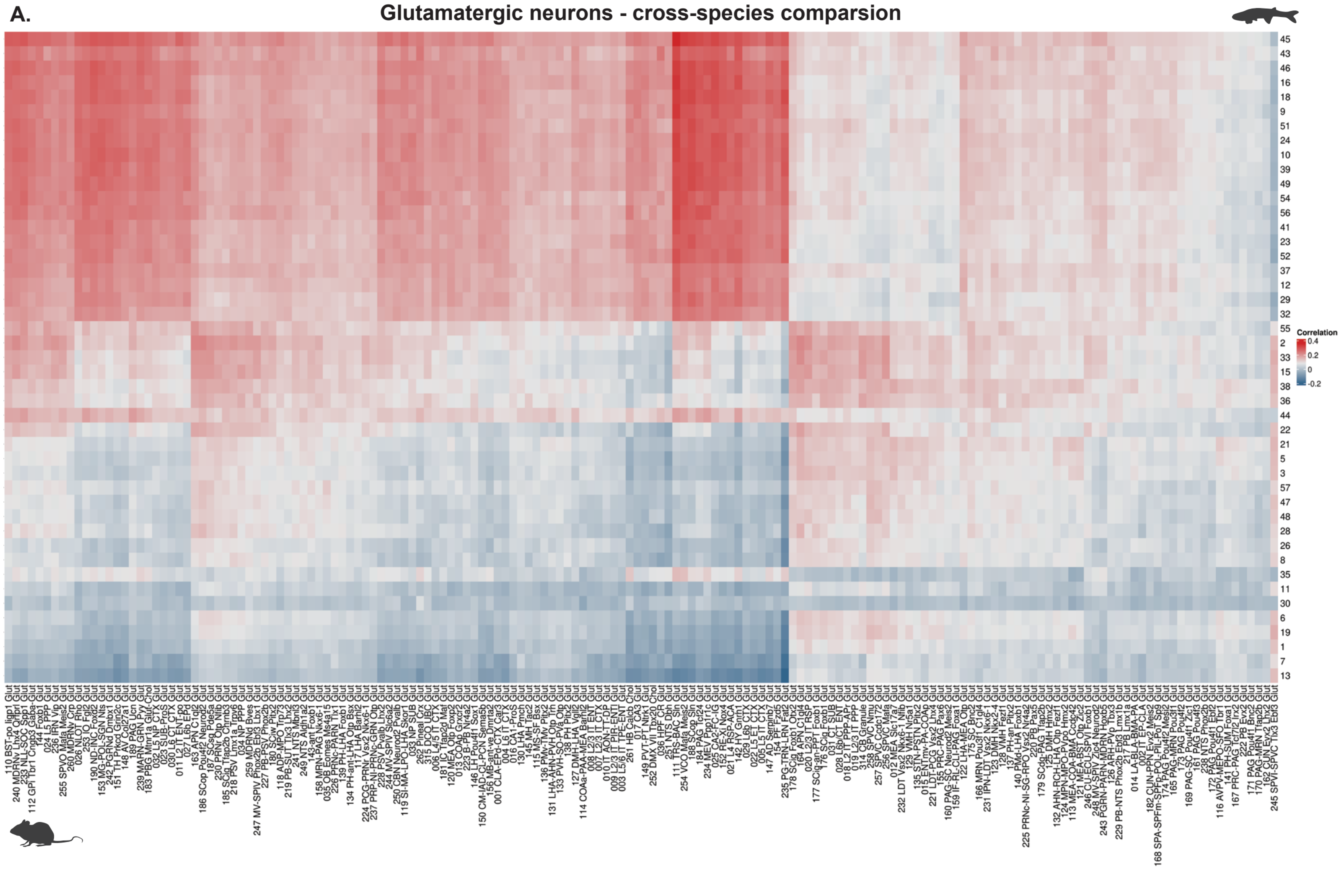

**B.** **GABAergic neurons - cross-species comparsion**

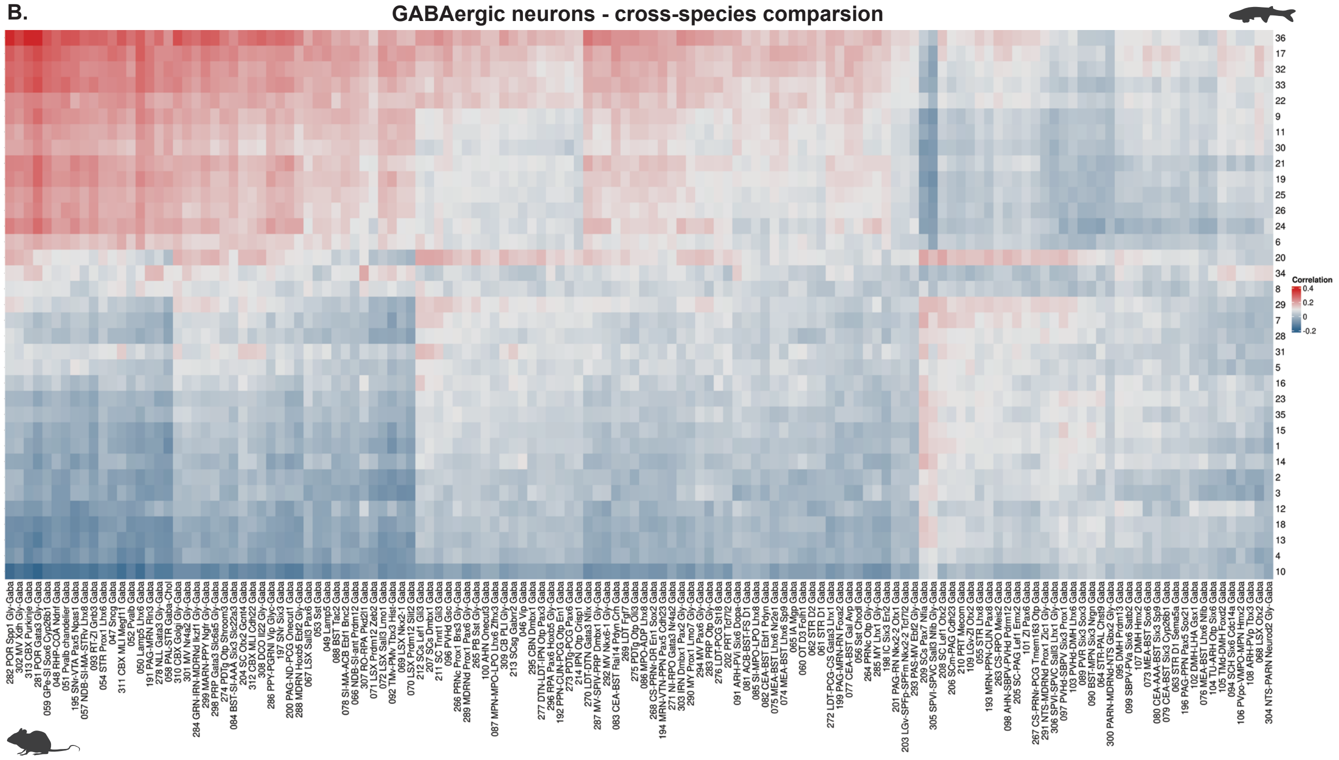

**fig. S34. Cross-species comparisons of *Danionella cerebrum* and mouse glutamatergic and GABAergic neurons (A)** Heatmap showing cross-species correlation (SAMap) between mouse glutamatergic and **(B)** GABAergic neuron subclasses (data source (*13*)) and *Danionella cerebrum* glutamatergic, and **(B)** GABAergic neuron clusters.

Figure S35.

A.

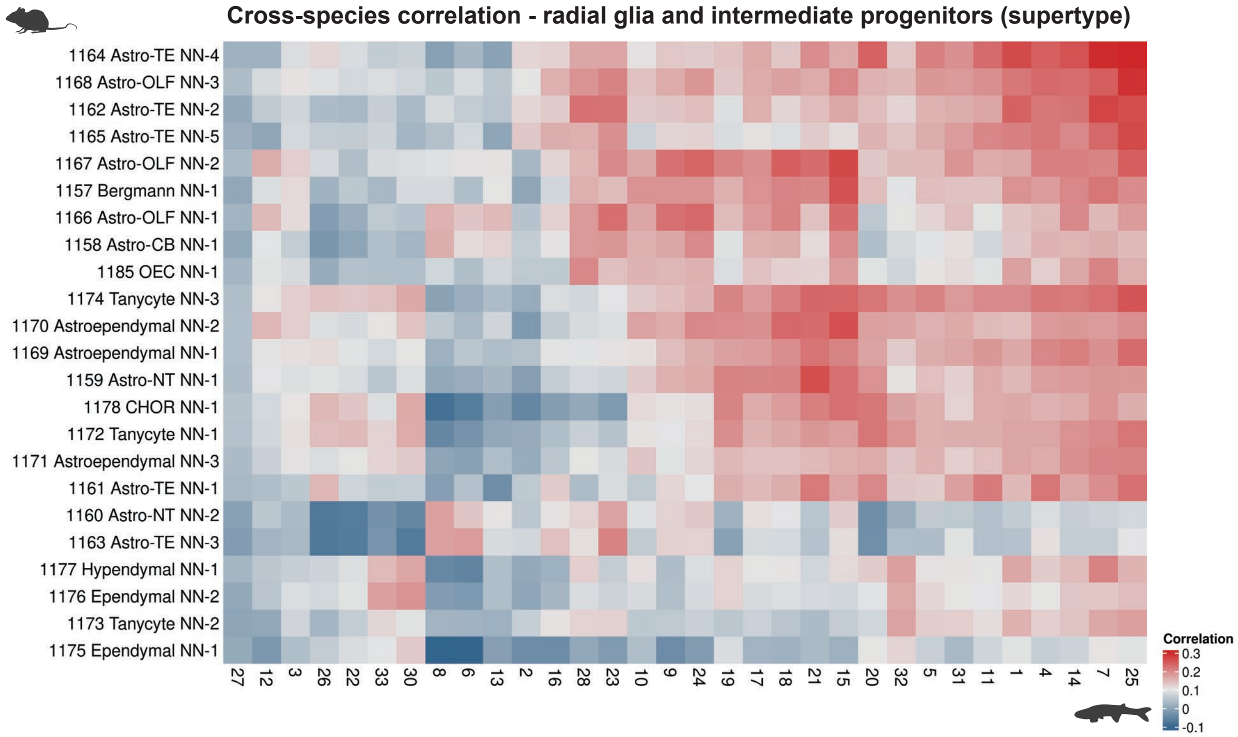

B.

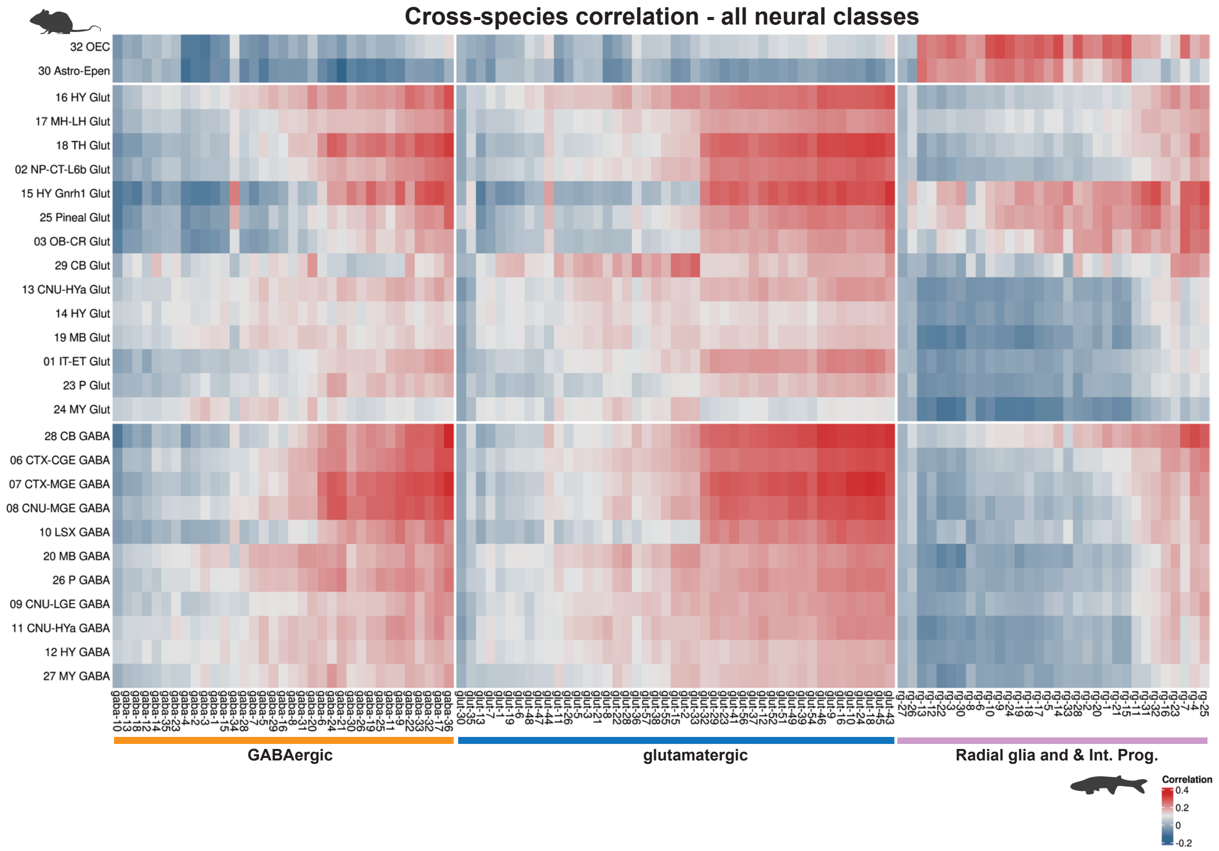

**fig. S35. Cross-species comparison of *Danionella cerebrum* and mouse radial glial and neural progenitor cells (A)** Heatmap showing cross-species correlation (SAMap) between mouse glial supertypes and *Danionella cerebrum* radial glial and intermediate progenitor clusters. **(B)** Integrated heatmap showing cross-species correlation (SAMap) between mouse glutamatergic and GABAergic neuron and glial classes and *Danionella cerebrum* glutamatergic, GABAergic neuron, and radial glia and intermediate progenitor clusters.

Figure S36.

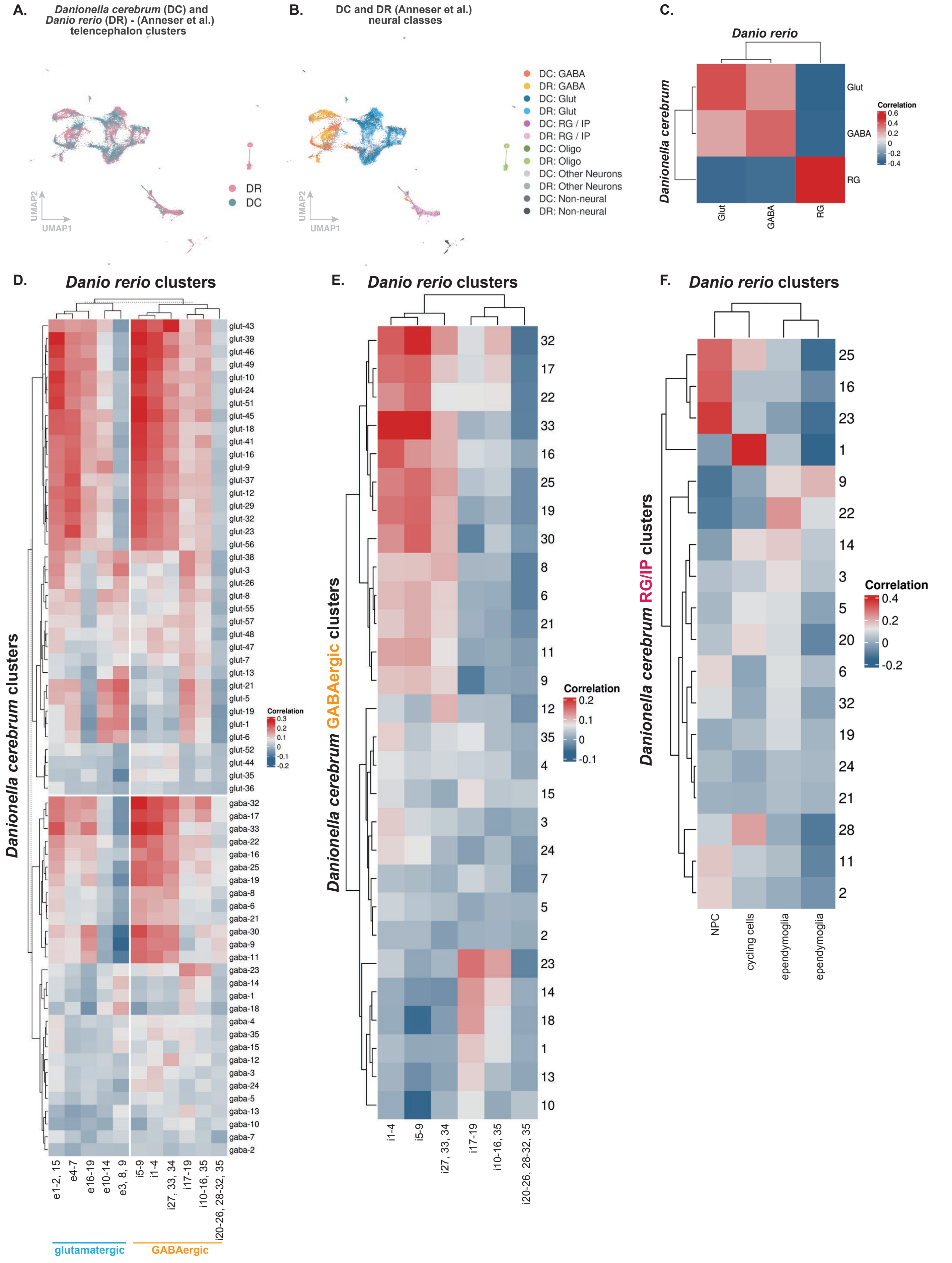
A.
Danionella cerebrum (DC) and Danio rerio (DR) - (Anneser et al.) telencephalon clusters
DR
DC
UMAP2
UMAP1
B.
DC and DR (Anneser et al.) neural classes
DC: GABA
DR: GABA
DC: Glut
DR: Glut
DC: RG / IP
DR: RG / IP
DC: Oligo
DR: Oligo
DC: Other Neurons
DR: Other Neurons
DC: Non-neural
DR: Non-neural
C.
Danio rerio
Danionella cerebrum
Glut
GABA
RG
Correlation
D.
Danio rerio clusters
Danionella cerebrum clusters
e1-2, 15
e4-7
e16-19
e10-14
e3, 8, 9
i5-9
i1-4
i27, 33, 34
i17-19
i10-16, 35
i20-26, 28-32, 35
glutamatergic
GABAergic
E.
Danio rerio clusters
Danionella cerebrum GABAergic clusters
Correlation
F.
Danio rerio clusters
Danionella cerebrum RG/IP clusters
Correlation
NPC
cycling cells
ependymoglia
ependymoglia

**fig. S36. Cross-species comparison of adult *Danionella cerebrum* and adult zebrafish telencephalon** **(A)** UMAP indicating interspecies mixing between the clusters from adult zebrafish telencephalon (L. Anneser et al., 2024; (*42*)) and *Danionella* telencephalon. and **(B)** by neurotransmitter class between adult zebrafish and *Danionella* telencephalic neural classes. **(C)** Correlation matrix heatmap (SAMap) showing similarity between *Danionella* cerebrum an adult zebrafish telencephalic neurotransmitter classes. **(D)** Integrated heatmap showing cross-species correlation (SAMap) between adult zebrafish glutamatergic and GABAergic neuron clusters and *Danionella cerebrum* glutamatergic and GABAergic neuron clusters. **(E)** Heatmap showing cross-species correlation (SAMap) between adult zebrafish GABAergic neuron clusters and *Danionella cerebrum* GABAergic neuron clusters. **(F)** Heatmap showing cross-species correlation (SAMap) between adult zebrafish radial glia/intermediate progenitor (RG/IP) clusters and *Danionella cerebrum* RG/IP clusters.

**Figure S37.**

**A.**

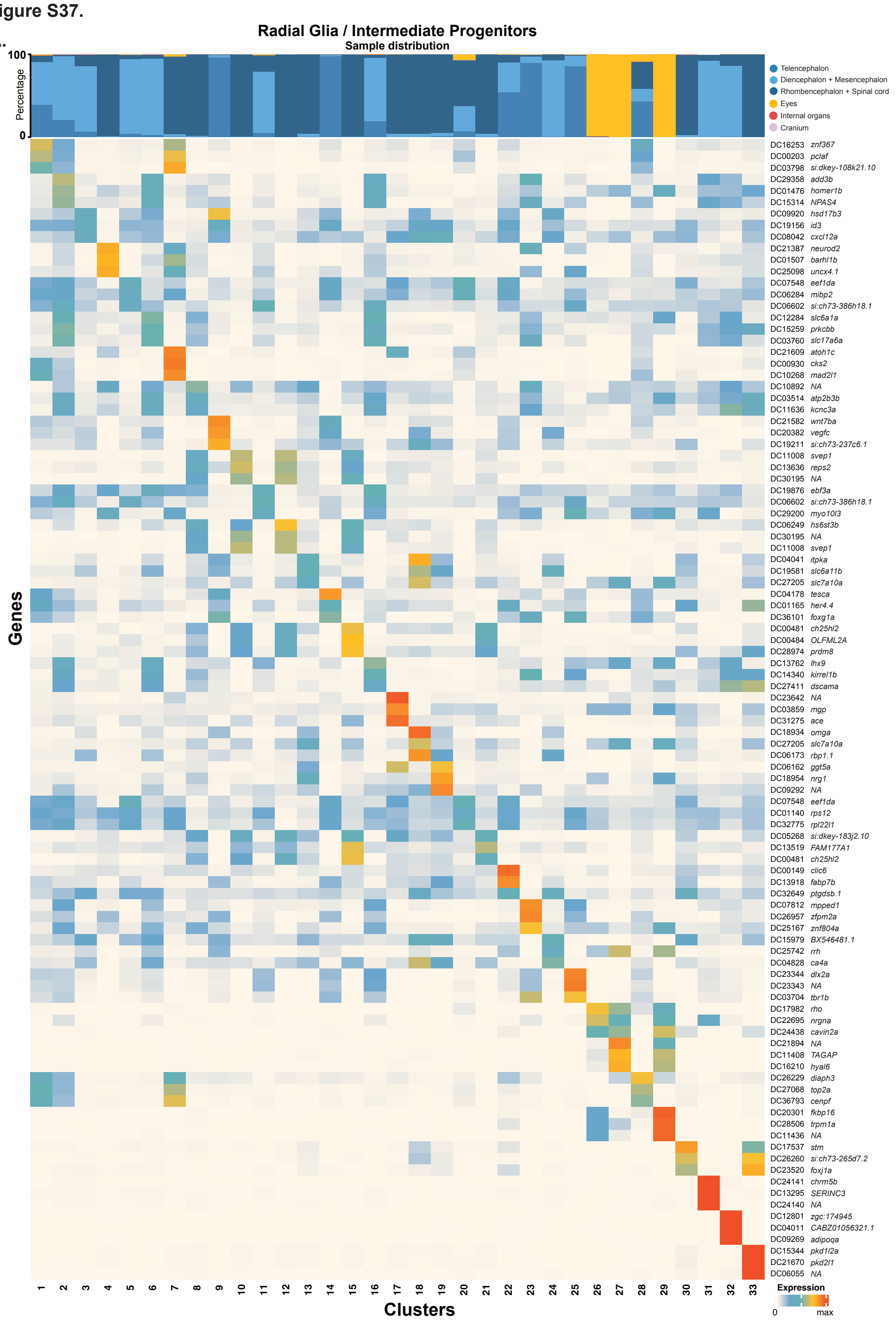

**B.**

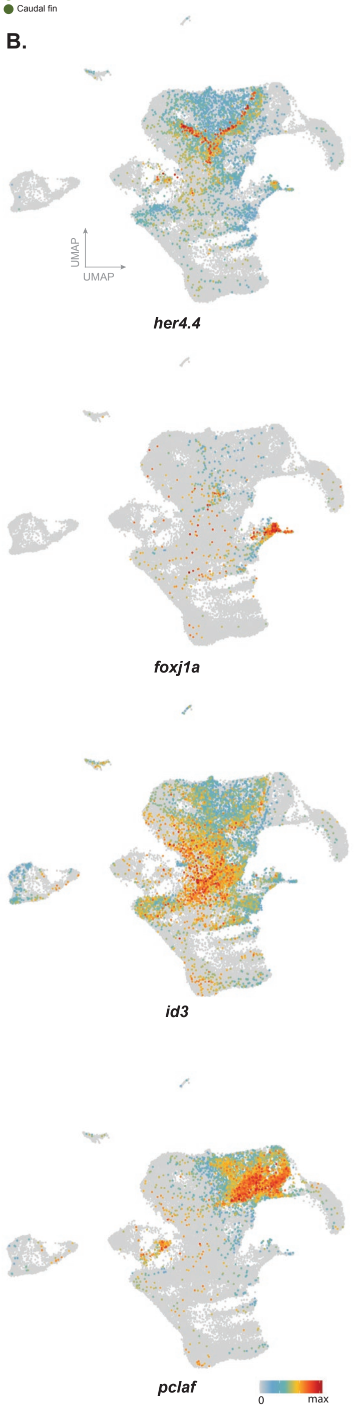

**fig. S37. Characterization of radial glia and neural progenitors (A)** Top three markers and sample distribution percentages of radial glia and intermediate progenitor clusters. **(B)** Features plots of genes marking quiescent and active radial glia, ciliated ependymal cells, and intermediate progenitors.

**Figure S38.**

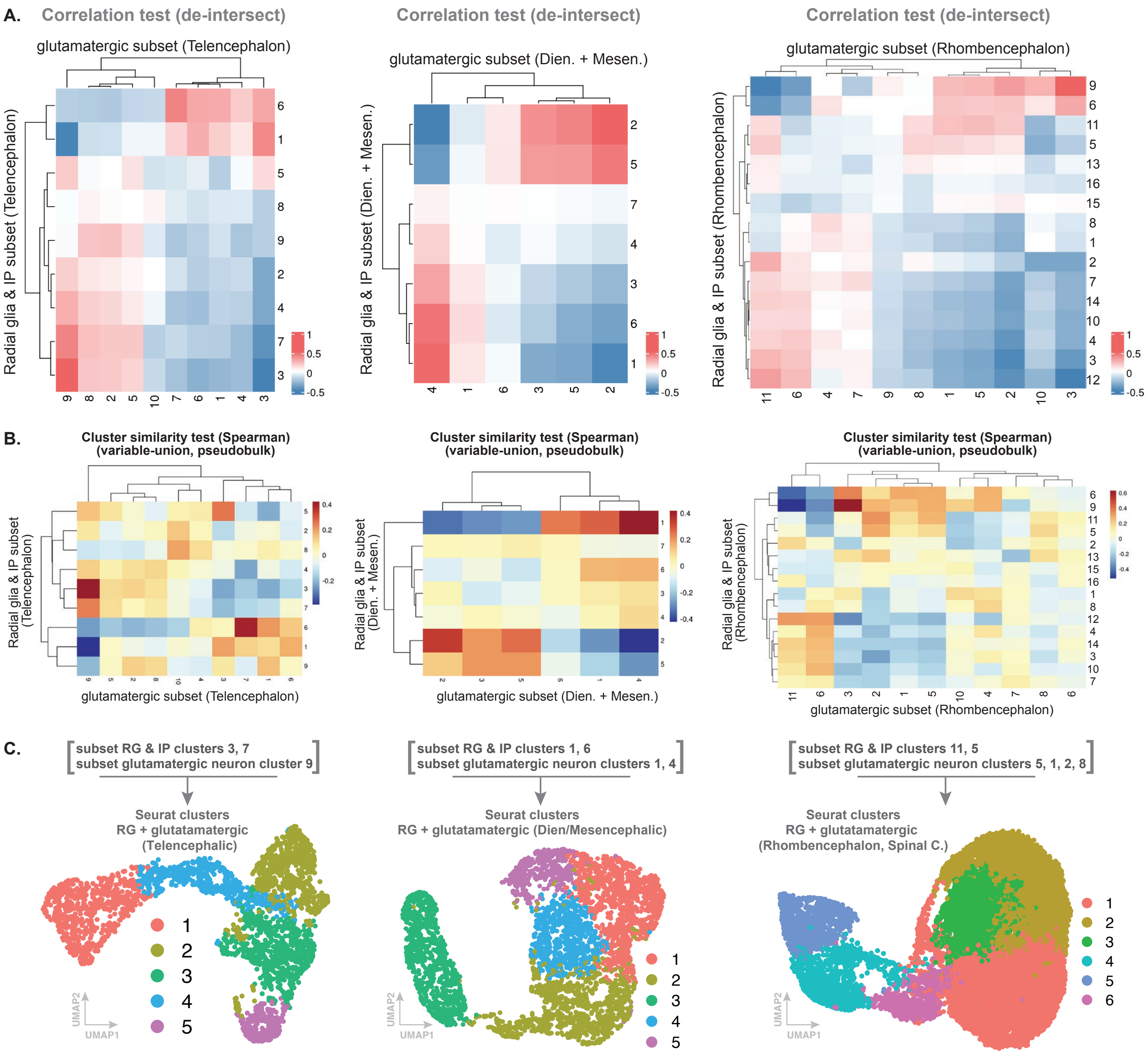
A.
Correlation test (de-intersect)
glutamatergic subset (Telencephalon)
Radial glia & IP subset (Telencephalon)
Correlation test (de-intersect)
glutamatergic subset (Dien. + Mesen.)
Radial glia & IP subset (Dien. + Mesen.)
Correlation test (de-intersect)
glutamatergic subset (Rhombencephalon)
Radial glia & IP subset (Rhombencephalon)
B.
Cluster similarity test (Spearman)
(variable-union, pseudobulk)
Radial glia & IP subset (Telencephalon)
glutamatergic subset (Telencephalon)
Radial glia & IP subset (Dien. + Mesen.)
glutamatergic subset (Dien. + Mesen.)
Radial glia & IP subset (Rhombencephalon)
glutamatergic subset (Rhombencephalon)
C.
subset RG & IP clusters 3, 7
subset glutamatergic neuron cluster 9
Seurat clusters
RG + glutatamatergic
(Telencephalic)
subset RG & IP clusters 1, 6
subset glutamatergic neuron clusters 1, 4
Seurat clusters
RG + glutatamatergic (Dien/Mesencephalic)
subset RG & IP clusters 11, 5
subset glutamatergic neuron clusters 5, 1, 2, 8
Seurat clusters
RG + glutatamatergic
(Rhombencephalon, Spinal C.)
UMAP1
UMAP2

**fig. S38. Correlation analyses for regional neurogenesis pseudotime trajectories (A)** Correlation heatmap generated to isolate populations of cells that consistently co-segregate across distinct datasets, showing transcriptional similarity between glutamatergic neurons from three separate CNS regions and radial glia clusters from corresponding brain regions. **(B)** Heatmaps generated using a separate method, Spearman's correlation test, to detect transcriptional similarity between glutamatergic neurons from three separate CNS regions and radial glia clusters from corresponding brain regions. **(C)** UMAP of cells from combined correlated glutamatergic neuron and radial glia clusters from three CNS regions.

Figure S39.

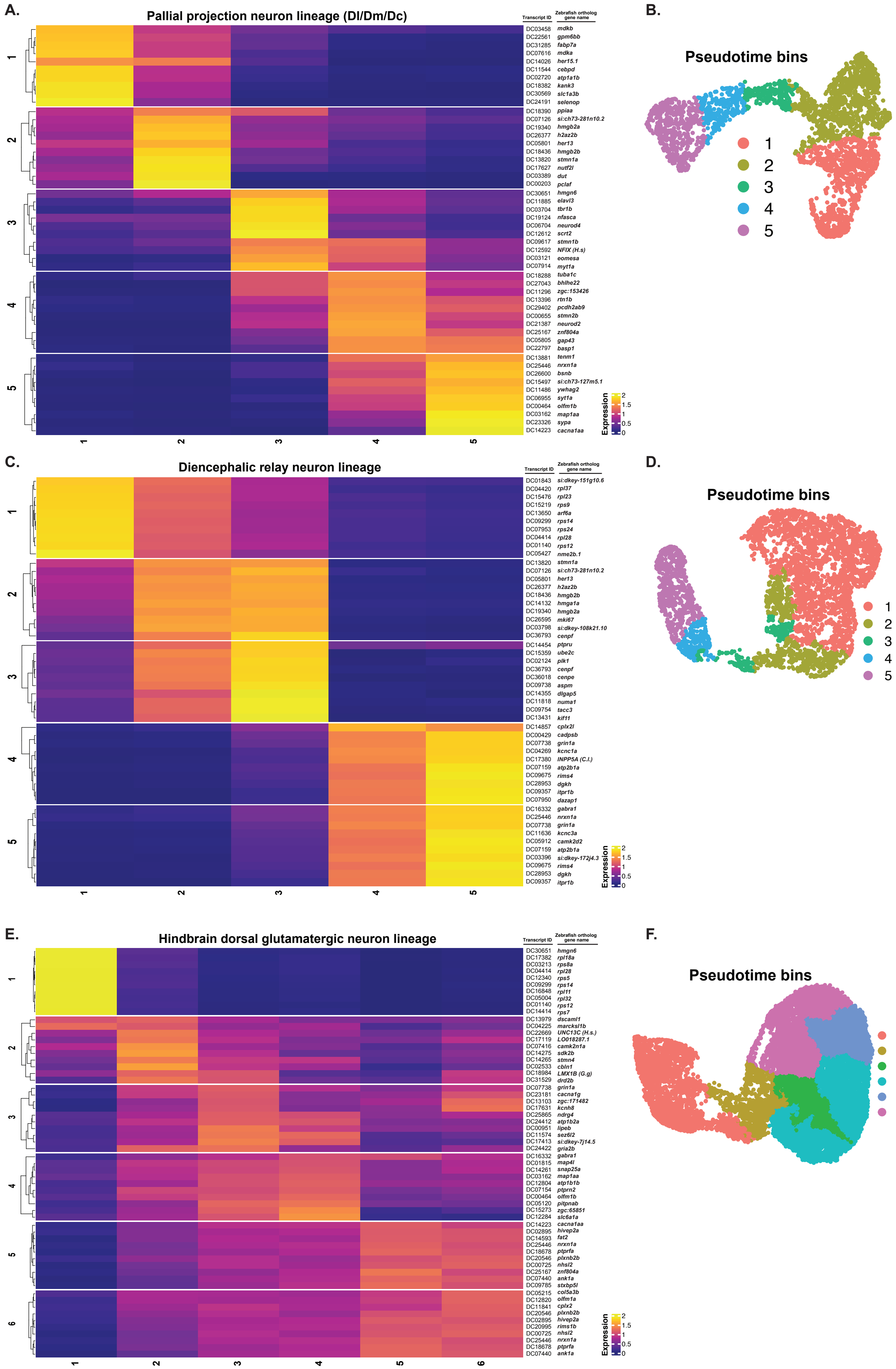
A.
Pallial projection neuron lineage (Dl/Dm/Dc)
Transcript ID
Zebrafish ortholog gene name
DC03458 mdkb
DC22561 gpm6bb
DC31285 fabp7a
DC07616 mdka
DC14026 her15.1
DC11544 cebpd
DC02720 atp1a1b
DC18382 kank3
DC30569 slc1a3b
DC24191 selenop
DC18390 ppiaa
DC07126 si:ch73-281n10.2
DC19340 hmgb2a
DC26377 h2az2b
DC05801 her13
DC18436 hmgb2b
DC13820 stmn1a
DC17627 nutf2l
DC03389 dut
DC00203 pclaf
DC30651 hmgn6
DC11885 elavl3
DC03704 tbr1b
DC19124 nfasca
DC06704 neurod4
DC12612 scrt2
DC09617 stmn1b
DC12592 NFIX (H.s)
DC03121 eomesa
DC07914 myt1a
DC18288 tuba1c
DC27043 bhlhe22
DC11296 zgc:153426
DC13396 rtn1b
DC29402 pcdh2ab9
DC00655 stmn2b
DC21387 neurod2
DC25167 znf804a
DC05805 gap43
DC22797 basp1
DC13881 tenm1
DC25446 nrxn1a
DC26600 bsnb
DC15497 si:ch73-127m5.1
DC11486 ywhag2
DC06955 syt1a
DC00464 olfm1b
DC03162 map1aa
DC23326 sypa
DC14223 cacna1aa
Expression
2
1.5
1
0.5
0
1
2
3
4
5
B.
Pseudotime bins
1
2
3
4
5
C.
Diencephalic relay neuron lineage
Transcript ID
Zebrafish ortholog gene name
DC01843 si:dkey-151g10.6
DC04420 rpl37
DC15476 rpl23
DC15219 rps9
DC13650 arf6a
DC09299 rps14
DC07953 rps24
DC04414 rpl28
DC01140 rps12
DC05427 nme2b.1
DC13820 stmn1a
DC07126 si:ch73-281n10.2
DC05801 her13
DC26377 h2az2b
DC18436 hmgb2b
DC14132 hmga1a
DC19340 hmgb2a
DC26595 mki67
DC03798 si:dkey-108k21.10
DC36793 cenpf
DC14454 ptpru
DC15359 ube2c
DC02124 plk1
DC36793 cenpf
DC36018 cenpe
DC09738 aspm
DC14355 dlgap5
DC11818 numa1
DC09754 tacc3
DC13431 kif11
DC14857 cplx2l
DC00429 cadpsb
DC07738 grin1a
DC04269 kcnc1a
DC17380 INPP5A (C.l.)
DC07159 atp2b1a
DC09675 rims4
DC28953 dgkh
DC09357 itpr1b
DC07950 dazap1
DC16332 gabra1
DC25446 nrxn1a
DC07738 grin1a
DC11636 kcnc3a
DC05912 camk2d2
DC07159 atp2b1a
DC03396 si:dkey-172j4.3
DC09675 rims4
DC28953 dgkh
DC09357 itpr1b
Expression
2
1.5
1
0.5
0
1
2
3
4
5
D.
Pseudotime bins
1
2
3
4
5
E.
Hindbrain dorsal glutamatergic neuron lineage
Transcript ID
Zebrafish ortholog gene name
DC30651 hmgn6
DC17382 rpl18a
DC03213 rps8a
DC04414 rpl28
DC12340 rps5
DC09299 rps14
DC16848 rpl11
DC05004 rpl32
DC01140 rps12
DC14414 rps7
DC13979 dscaml1
DC04225 marcksl1b
DC22669 UNC13C (H.s.)
DC17119 LO018287.1
DC07416 camk2n1a
DC14275 sdk2b
DC14265 stmn4
DC02533 cbln1
DC18984 LMX1B (G.g)
DC31529 drd2b
DC07738 grin1a
DC23181 cacna1g
DC13103 zgc:171482
DC17631 kcnh8
DC25865 ndrg4
DC24412 atp1b2a
DC00951 lipeb
DC11574 sez6l2
DC17413 si:dkey-7j14.5
DC24422 gria2b
DC16332 gabra1
DC01815 map4l
DC14261 snap25a
DC03162 map1aa
DC12804 atp1b1b
DC07154 ptprn2
DC00464 olfm1b
DC05120 pitpnab
DC15273 zgc:65851
DC12284 slc6a1a
DC14223 cacna1aa
DC02895 hivep2a
DC14593 fat2
DC25446 nrxn1a
DC18678 ptprfa
DC20546 plxnb2b
DC00725 nhsl2
DC25167 znf804a
DC07440 ank1a
DC09785 stxbp5l
DC05215 col5a3b
DC12820 olfm1a
DC11841 cplx2
DC20546 plxnb2b
DC02895 hivep2a
DC20995 rims1b
DC00725 nhsl2
DC25446 nrxn1a
DC18678 ptprfa
DC07440 ank1a
Expression
2
1.5
1
0.5
0
1
2
3
4
5
6
F.
Pseudotime bins
1
2
3
4
5
6

**fig. S39. Pseudotime bins for regional neurogenesis trajectories in the adult brain (A)** Heatmap showing top genes representing pseudotime bins (monocle3) representing changes in gene expression in telencephalic projection neurons along pseudotime. **(B)** UMAP showing pseudotime bins for the telencephalon projection neuron lineage. **(C-D)** Heatmap showing top genes from each pseudotime bin and UMAP showing the pseudotime bins from a diencephalic excitatory neuron lineage. **(E-F)** Heatmap showing top genes from each pseudotime bin and UMAP showing the pseudotime bins from a hindbrain excitatory neuron lineage.

Figure S40.

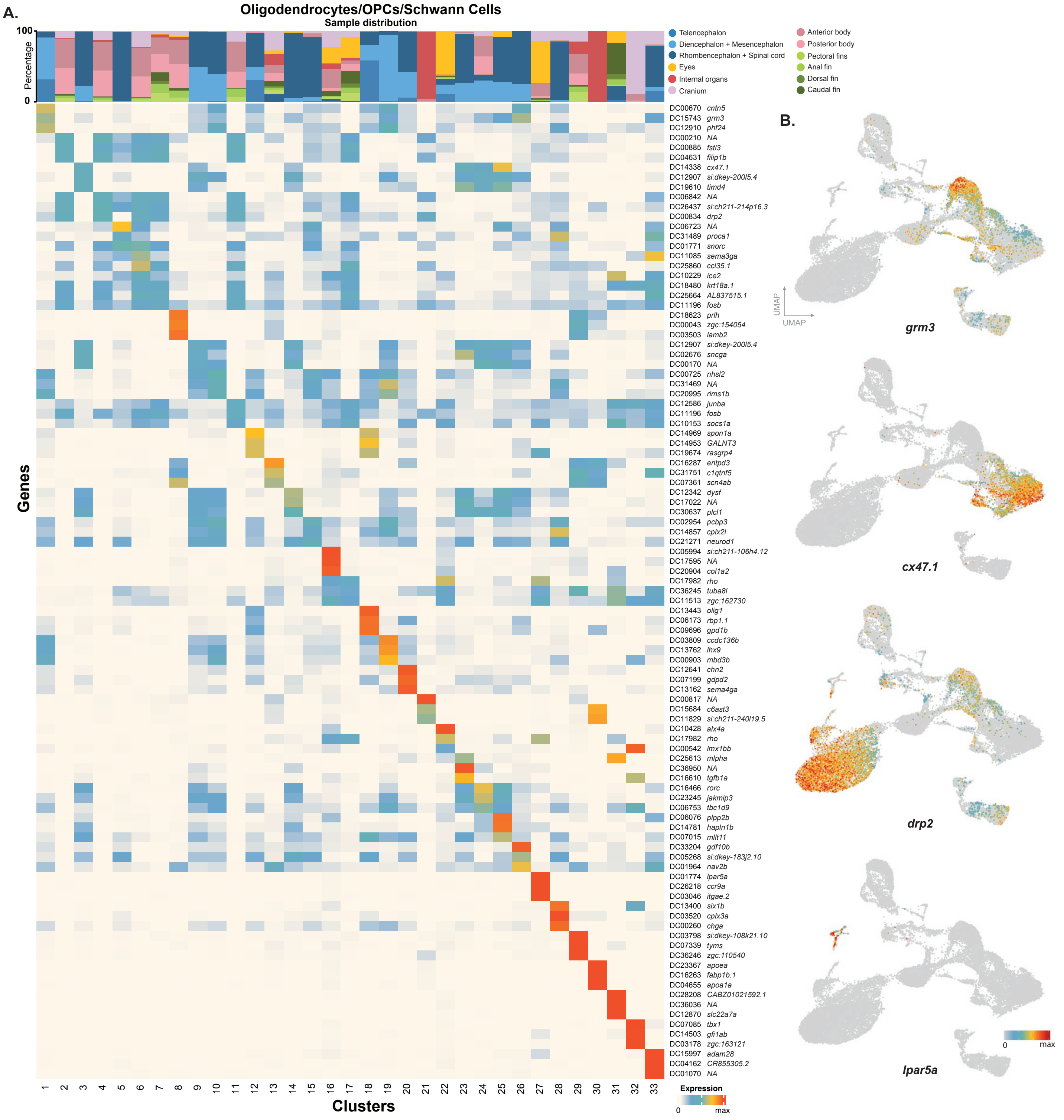
A.
Oligodendrocytes/OPCs/Schwann Cells
Sample distribution
Percentage
Telencephalon
Diencephalon + Mesencephalon
Rhombencephalon + Spinal cord
Eyes
Internal organs
Cranium
Anterior body
Posterior body
Pectoral fins
Anal fin
Dorsal fin
Caudal fin
Genes
Clusters
Expression
B.
UMAP
grm3
cx47.1
drp2
lpar5a

**fig. S40. Characterization of oligodendrocytes, oligodendrocyte progenitor cells (OPCs), and Schwann cells (A)** Top three markers and sample distribution percentages of oligodendrocytes, oligodendrocyte progenitors, and Schwann cells. Top columns represent the distribution of experimental samples across the clusters. **(B)** Feature plots of select genes enriched in distinct experimental samples and clusters are shown.

Figure S41.

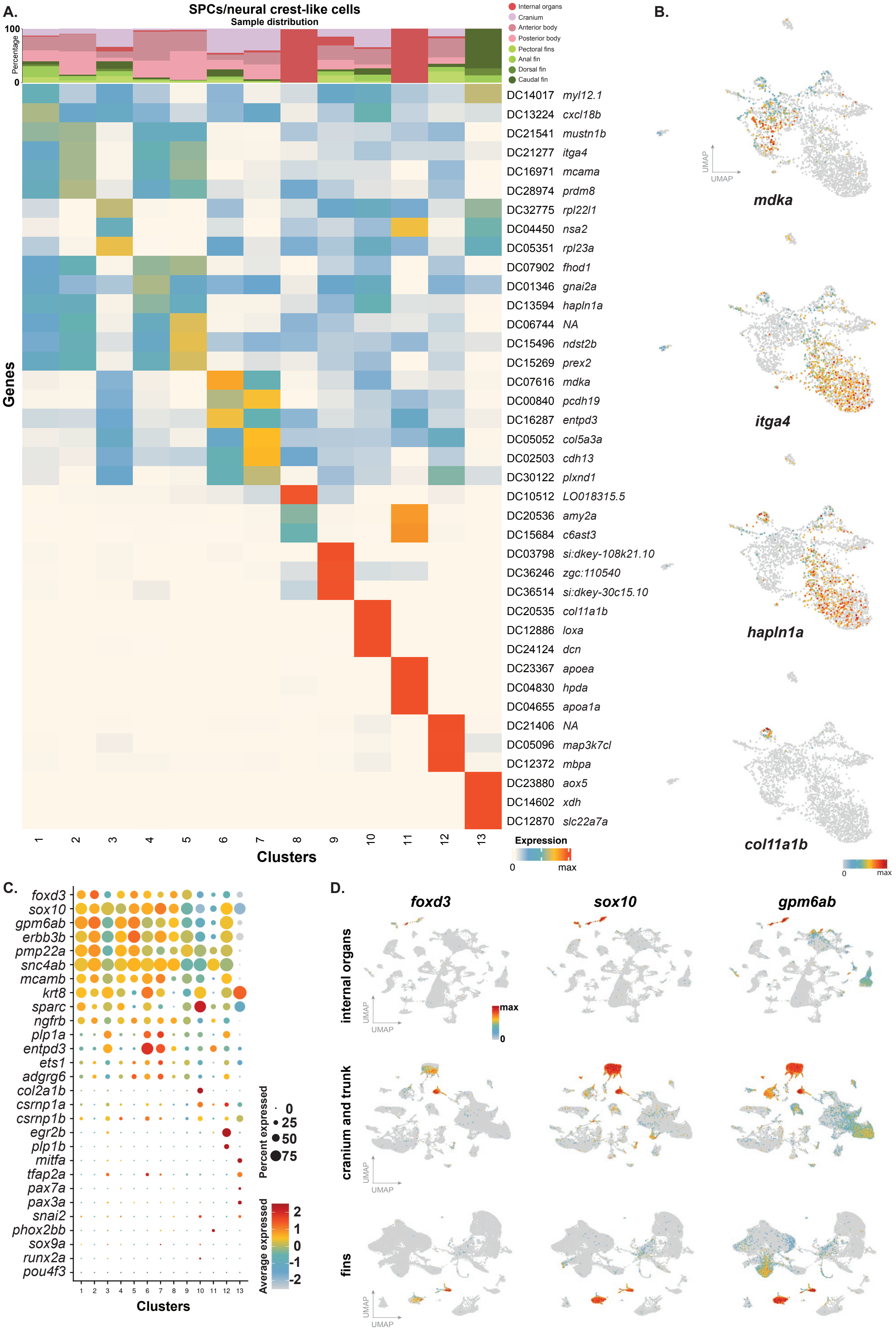

A.
SPCs/neural crest-like cells
Sample distribution
Percentage
100
0
Internal organs
Cranium
Anterior body
Posterior body
Pectoral fins
Anal fin
Dorsal fin
Caudal fin
Genes
DC14017 myl12.1
DC13224 cxcl18b
DC21541 mustn1b
DC21277 itga4
DC16971 mcama
DC28974 prdm8
DC32775 rpl22l1
DC04450 nsa2
DC05351 rpl23a
DC07902 fhod1
DC01346 gnai2a
DC13594 hapln1a
DC06744 NA
DC15496 ndst2b
DC15269 prex2
DC07616 mdka
DC00840 pcdh19
DC16287 entpd3
DC05052 col5a3a
DC02503 cdh13
DC30122 plxnd1
DC10512 LO018315.5
DC20536 amy2a
DC15684 c6ast3
DC03798 si:dkey-108k21.10
DC36246 zgc:110540
DC36514 si:dkey-30c15.10
DC20535 col11a1b
DC12886 loxa
DC24124 dcn
DC23367 apoea
DC04830 hpda
DC04655 apoa1a
DC21406 NA
DC05096 map3k7cl
DC12372 mbpa
DC23880 aox5
DC14602 xdh
DC12870 slc22a7a
1 2 3 4 5 6 7 8 9 10 11 12 13
Clusters
Expression
0 max
B.
UMAP
mdka
itga4
hapln1a
col11a1b
0 max
C.
foxd3
sox10
gpm6ab
erbb3b
pmp22a
snc4ab
mcamb
krt8
sparc
ngfrb
plp1a
entpd3
ets1
adgrg6
col2a1b
csrnp1a
csrnp1b
egr2b
plp1b
mitfa
tfap2a
pax7a
pax3a
snai2
phox2bb
sox9a
runx2a
pou4f3
1 2 3 4 5 6 7 8 9 10 11 12 13
Clusters
Percent expressed
0
25
50
75
Average expressed
2
1
0
-1
-2
D.
foxd3
sox10
gpm6ab
internal organs
cranium and trunk
fins
UMAP
max
0

**fig. S41. Characterization of Schwann cell precursor (SCPs) and neural-crest-like cells (A)** Top three markers and sample distribution percentages of Schwann cell precursors (SCP) and neural-crest-like cells. Top columns represent the distribution of experimental samples across the clusters. **(B)** Feature plots of select genes enriched in distinct clusters are shown. **(C)** Expanded dot plot, including genes previously shown to be expressed in SCPs, and different neural crest lineages (*16, 45, 109-111*). **(D)** Feature plots indicating Schwann cell precursor and neural-crest markers *foxd3*, *sox10* and *gpm6ab* expression in internal organs, cranium and trunk, and fin samples.

Figure S42.

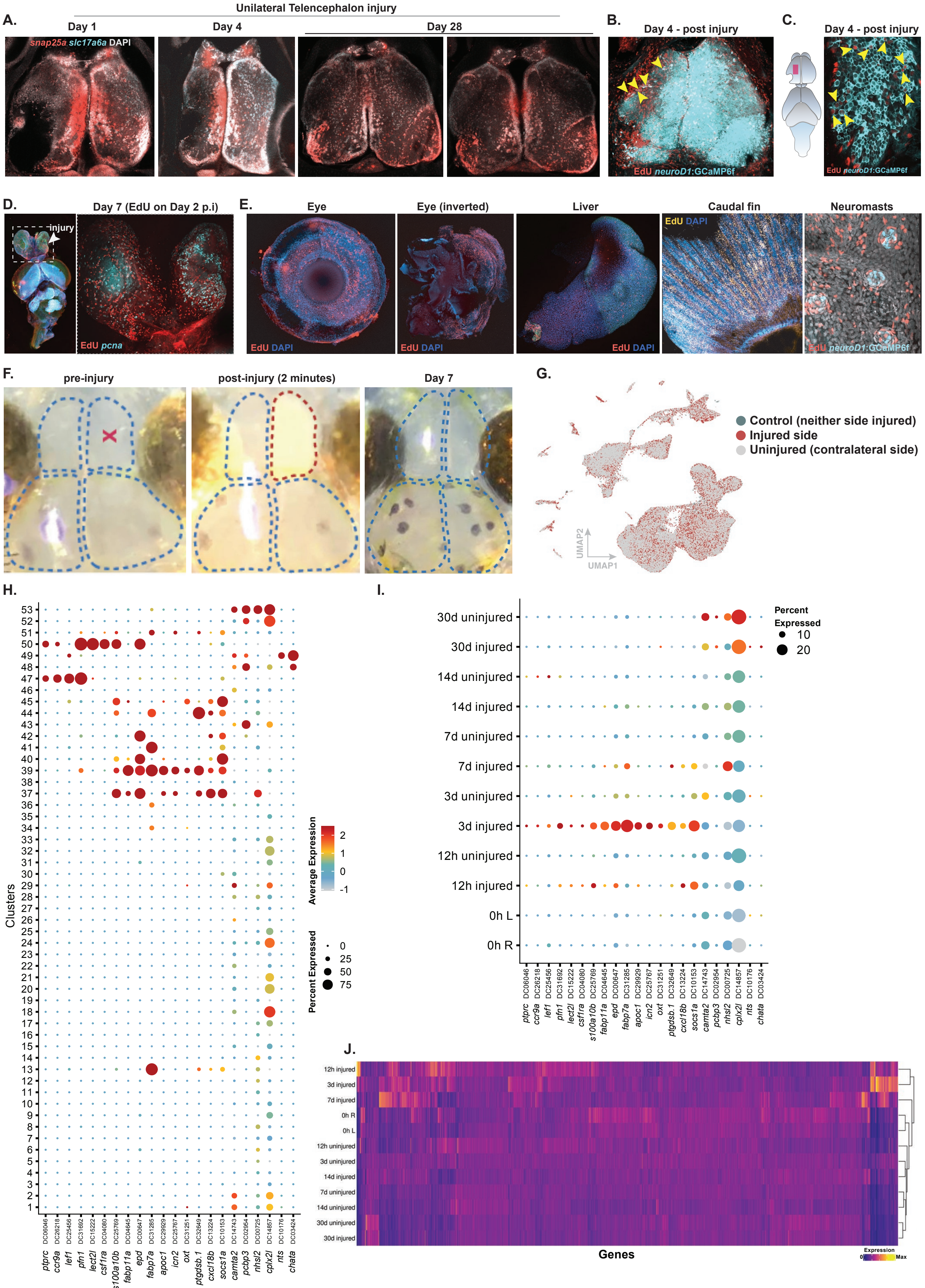
A.
Unilateral Telencephalon injury
Day 1
Day 4
Day 28
snap25a slc17a6a DAPI
B.
Day 4 - post injury
EdU neuroD1:GCaMP6f
C.
Day 4 - post injury
EdU neuroD1:GCaMP6f
D.
Day 7 (EdU on Day 2 p.i)
injury
EdU pcna
E.
Eye
Eye (inverted)
Liver
Caudal fin
Neuromasts
EdU DAPI
EdU neuroD1:GCaMP6f
F.
pre-injury
post-injury (2 minutes)
Day 7
G.
Control (neither side injured)
Injured side
Uninjured (contralateral side)
UMAP1
UMAP2
H.
Clusters
Average Expression
Percent Expressed
ptprc DC06046
ccr9a DC26218
lef1 DC25456
pfn1 DC31692
lect2l DC15222
csf1ra DC04080
s100a10b DC25769
fabp11a DC04645
epd DC00647
fabp7a DC31285
apoc1 DC29929
icn2 DC25767
oxt DC31251
ptgdsb.1 DC32649
cxcl18b DC13224
socs1a DC10153
camta2 DC14743
pcbp3 DC02954
nhsl2 DC00725
cplx2l DC14857
nts DC10176
chata DC03424
I.
30d uninjured
30d injured
14d uninjured
14d injured
7d uninjured
7d injured
3d uninjured
3d injured
12h uninjured
12h injured
0h L
0h R
Percent Expressed
J.
Genes
Expression
Max

**fig. S42. Regeneration in the adult *Danionella cerebrum* brain (A)** Unilateral tissue removal in telencephalon and regeneration. **(B)** EdU labeling of newly integrating neurons into the telencephalon following unilateral injury. **(C)** Zoomed dorsal telencephalon inset from the injured side showing EdU labeling of newly integrating neurons into the telencephalon following unilateral injury. **(D)** Immunostaining showing dividing cells at the site of injury on day 5 following injury (*pcna*) and EdU labeling marking integrated cellular progeny. **(E)** EdU labeling indicates widespread EdU-labeled cellular progeny (labeled 5 days after a 24-hour EdU pulse) across different tissues. **(F)** Unilateral telencephalon injury causes a transient period of opaqueness in the injured lobe. Red "x" marks the injury region. **(G)** UMAP showing cells from the injured and uninjured sides of the telencephalon and cells from the control, uninjured telencephalon. **(H)** Expanded dot plot showing a subset of top genes expressed in each time-point-specific cluster. **(I)** Dot plot showing a subset of top genes expressed in each time point. **(J)** Heatmap of top differentially expressed genes in each time point (list of genes in table S9).

**Figure S43.**

**A.**

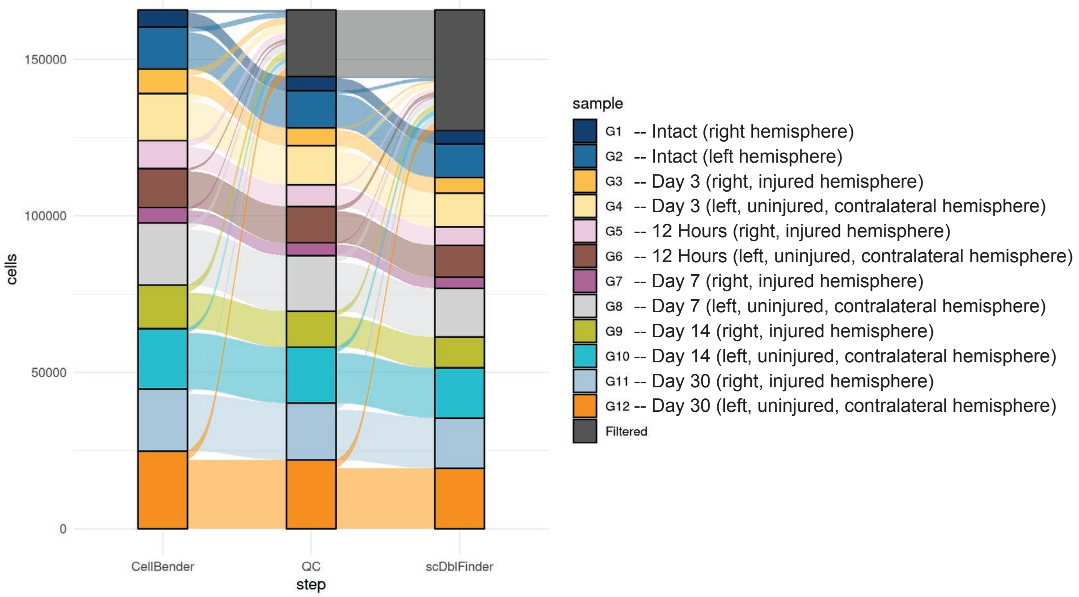

**B.**

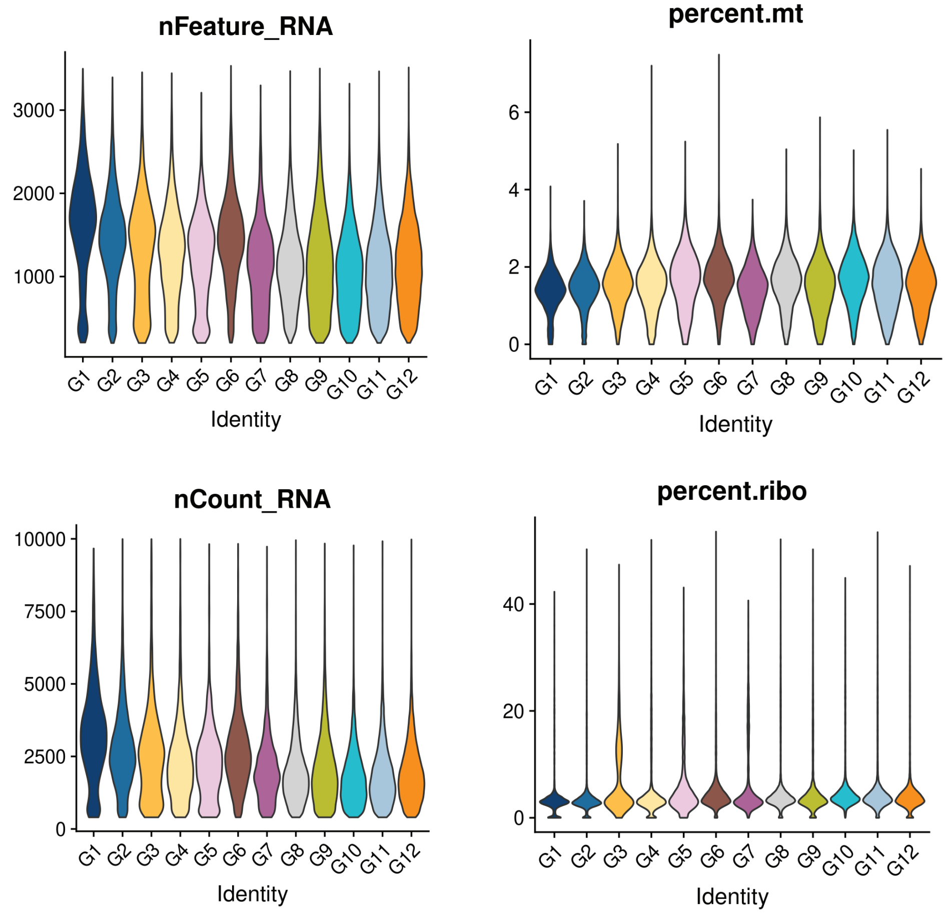

**fig. S43**. **General overview and quality control of the telencephalon regeneration single-cell RNA-seq data (A)** Sankey plot showing computational quality control steps taken and the list of telencephalon regeneration samples **(B)** Violin plots showing the percentage of the number of uniquely expressed genes (nFeatures) and the percentage of the number of RNA molecules (UMIs) detected across all genes (nCounts), and violin plots indicating the percentage of mitochondrial and ribosomal genes detected across these experimental samples.

**Figure S44.**

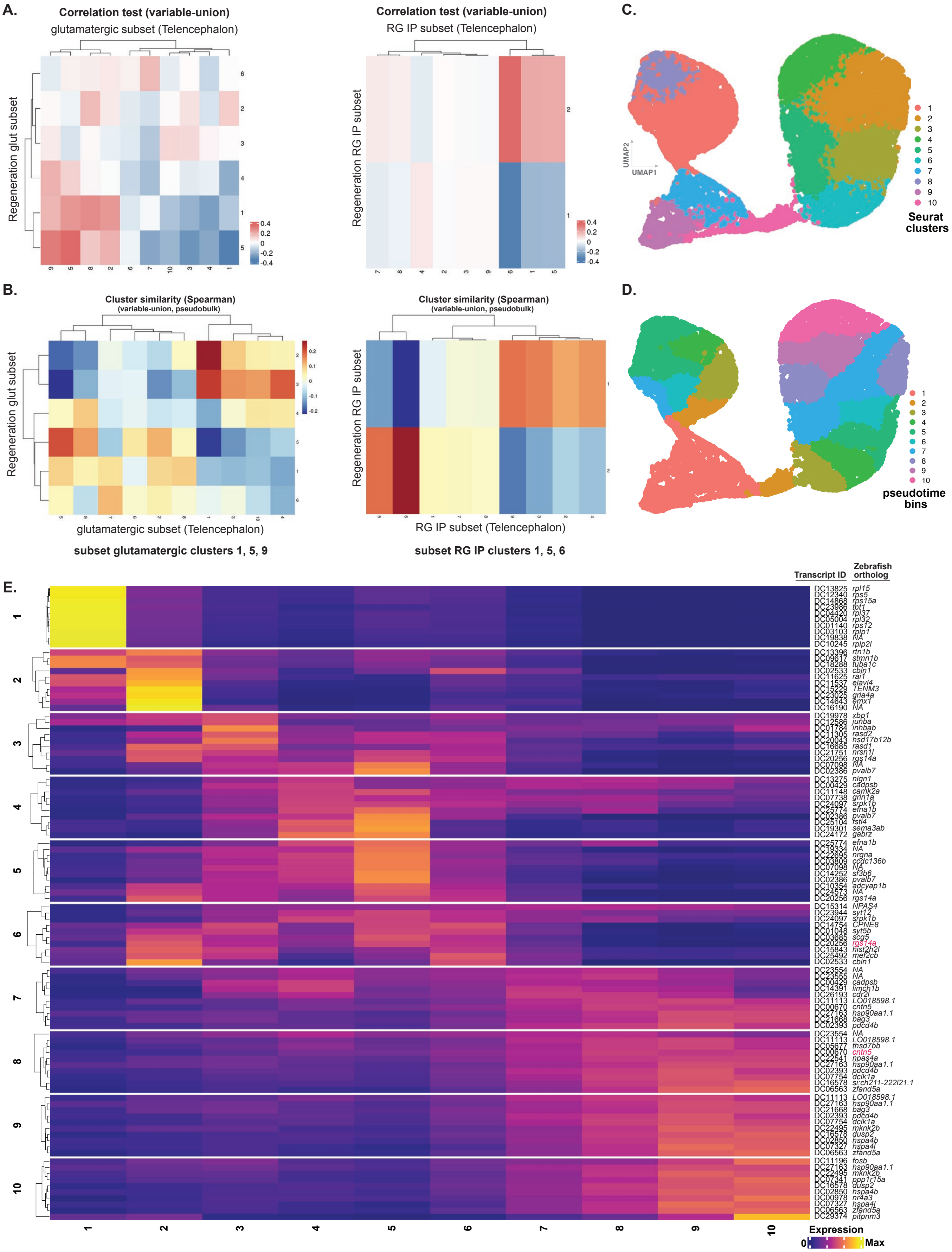
A.
Correlation test (variable-union)
glutamatergic subset (Telencephalon)
Regeneration glut subset
Correlation test (variable-union)
RG IP subset (Telencephalon)
Regeneration RG IP subset
C.
UMAP2
UMAP1
Seurat clusters
B.
Cluster similarity (Spearman)
(variable-union, pseudobulk)
Regeneration glut subset
glutamatergic subset (Telencephalon)
subset glutamatergic clusters 1, 5, 9
Cluster similarity (Spearman)
(variable-union, pseudobulk)
Regeneration RG IP subset
RG IP subset (Telencephalon)
subset RG IP clusters 1, 5, 6
D.
pseudotime bins
E.
Transcript ID
Zebrafish ortholog
Expression
0
Max

**fig. S44. Pseudotime analysis of regeneration in the adult brain (A)** Heatmap showing transcriptional similarity between glutamatergic neurons and glutamatergic neurons from regenerating samples, left; and radial glia clusters and radial glia from regeneration samples. **(B)** Correlation heatmap generated using a separate method, Spearman's correlation test. **(C)** UMAP representation of cells from glutamatergic and radial glia clusters that correspond to regeneration clusters. **(D)** UMAP showing pseudotime bins. **(E)** Heatmap showing top genes from each pseudotime bin from predicted regenerating lineages following unilateral telencephalon injury. Genes marked in red indicated the predicted regenerating glutamatergic lineages.
